# The lower jaw of Devonian ray-finned fishes (Actinopterygii): anatomy, relationships, and functional morphology

**DOI:** 10.1101/2025.01.30.635695

**Authors:** Ben Igielman, Rodrigo Tinoco Figueroa, Robert R Higgins, Stephanie E Pierce, Michael I. Coates, Emily M. Troyer, Vincent Fernandez, Kathleen Dollman, Jing Lu, Min Zhu, Matt Friedman, Sam Giles

## Abstract

Actinopterygii is a major extant vertebrate group, but limited data are available for its earliest members. Here we investigate the morphology of Devonian actinopterygians, focusing on the lower jaw. We use X-Ray Computed Tomography (XCT) to provide comprehensive descriptions of the mandibles of 19 species, which span the whole of the Devonian and represent roughly two thirds of all taxa known from more than isolated or fragmentary material. Our findings corroborate previous reports in part but reveal considerable new anatomical data and represent the first detailed description for roughly half of these taxa. The mandibles display substantial variation in size, spanning more than an order of magnitude. Although most conform to a generalized pattern of a large dentary and one or two smaller infradentaries, XCT data reveal significant differences in the structure of the jaw and arrangement of teeth that may be of functional relevance. We report the presence of a rudimentary coronoid process in several taxa, contributed to by the dentary and/or infradentaries, as well a raised articular region, resulting in a mandible with an offset bite and that functions as a bent level arm. Among the most striking variation is that of tooth morphology: several taxa have heterodont dentary teeth that vary in size and orientation, and multiple variations on enlarged, whorl-like and posteriorly-oriented anterior coronoid dentition are observed. We use these new data to revise morphological characters that may be of phylogenetic significance and consider the possible functional implications of these traits. The observed variation in mandible form and structure suggests previously unappreciated functional diversity among otherwise morphologically homogenous Devonian ray-finned fishes.

## 1. INTRODUCTION

Actinopterygian, or ray-finned, fishes represent half of all living vertebrate species. This ecologically diverse group traces its ancestry deep within the Palaeozoic, over 400 million years ago. The traditional narrative of Palaeozoic ray-finned fish evolution is that the Devonian (419 Ma – 359 Ma) was a period of low abundance and diversity (Trewin, 1986; Sallan & Coates, 2010; Friedman, 2015), with morphological, ecological and lineage diversity substantially increasing in the Carboniferous (359 Ma – 299 Ma; Sallan & Friedman, 2012; Sallan & Coates, 2013; Sallan, 2014). Actinopterygian taxa are few both in terms of number of lineages and taxonomic diversity within individual assemblages; approximately 20 actinopterygian genera are known from the entire Devonian (Giles et al., 2015b; Sallan & Coates, 2010; Henderson et al., 2023), and only five species out of over 50 fishes known from the late Devonian Gogo Lagerstätte are ray-fins (Long & Trinajstic, 2010). During the Devonian, actinopterygians were also morphologically conservative (Sallan & Friedman, 2012), implying that they occupied fewer ecological niches than coexisting sarcopterygians (Friedman, 2015). In contrast, after the End-Devonian Mass Extinction (Hangenberg event), Carboniferous actinopterygians increase in numerical diversity by an order of magnitude (Sallan, 2014; Henderson et al., 2023), exhibiting a much greater variety of body and skull shapes (Sallan & Coates, 2010; Sallan & Friedman, 2012; Sallan & Coates, 2013; Wilson et al., 2021), as well as feeding innovations (Friedman et al., 2019).

This narrative appears to be corroborated by most phylogenetic analyses, which tend to recover a grade of dead-end Devonian lineages with few surviving the end-Devonian mass extinction (Taverne, 1997; Coates, 1999; Swartz, 2009; Giles et al., 2017; but see Cloutier & Arratia, 2004). Stable clades of Devonian taxa are rarely recovered or corroborated across multiple analyses (Gardiner & Schaeffer, 1989; Gardiner et al., 2005; Friedman & Blom, 2006; Long et al., 2008; Choo et al., 2009; Choo, 2012 Giles et al., 2015b) and have few links to Carboniferous and younger lineages (Wilson et al., 2018; Caron et al. 2023). Difficulties in establishing stable phylogenetic relationships amongst Devonian actinopterygians – and to stratigraphically younger taxa – are compounded by the challenging preservation of many specimens, coupled with inaccessible internal anatomy in all but a few exemplary taxa (e.g. Gardiner, 1984). More recently, the application of tomographic techniques has led to the acquisition of detailed internal and external morphological data for both new and previously known Devonian taxa (Giles et al., 2015a,b; Figueroa et al., 2021; Newman et al., 2021; Giles et al., 2023). Incorporation of these data into phylogenetic analyses demonstrates greater anatomical diversity in the Devonian (Lu et al., 2016), broader biogeographical distribution (Figueroa et al., 2021), and greater lineage survivorship across the Hangenberg event (Giles et al., 2023) than previously supposed. Together, these new findings imply extensive cryptic lineage diversification in the Devonian seeding the radiation—and conspicuous morphological innovations—seen in actinopterygians during the Carboniferous. Rather than a collection of evolutionary dead-ends, Devonian taxa now appear central to understanding the early radiation and evolutionary dynamics of Palaeozoic actinopterygians, but diversity patterns still remain a major source of uncertainty.

Crucially, recent studies highlight anatomy with potential significance for informing function, and ecology, as well as relationships. Functional modifications to the lower jaw in particular have been implicated in actinopterygian diversification (Schaeffer & Rosen, 1961; Lauder, 1980; Sallan, 2014). In line with general interpretations of Devonian actinopterygians, the mandibles of the earliest ray-fins are generally held to be structurally conservative, with one or two notable exceptions (e.g. Dunkle & Schaeffer, 1973). Patterns of morphological and adaptive change have been studied in more detail for other early osteichthyan groups, but prior investigations of early actinopterygian lower jaws have generally been confined to primary considerations of morphological characters (Gardiner, 1984; Gardiner & Schaeffer, 1989; Gardiner et al., 2005) or as ‘primitive’ outgroups lacking specialisations for comparison to their sarcopterygian counterparts (Zhu & Yu, 2004) or neopterygian descendants (Lauder, 1980). Here, we use X-Ray CT to carry out a comprehensive review of the mandibles of almost all known Devonian actinopterygian taxa, with the following aims:

1. Describe mandibular anatomy in all Devonian actinopterygian taxa amenable to X-ray CT.
2. Assemble a 3D dataset of Devonian actinopterygian mandibles for use in future functional and morphometric analyses.
3. Synthesise mandibular morphological data into discrete characters for use in phylogenetic analyses and make inferences about jaw evolution in early actinopterygians.
4. Discuss the likely significance of morphological variation for jaw functions and ecological roles of early actinopterygian taxa.

## 2. MATERIALS AND METHODS

### 2.1. Institutional abbreviations

ANSP, The Academy of Natural Sciences of Drexel University, Philadelphia, USA; AM, Australian Museum, Sydney, Australia; AMNH, American Museum of Natural History, New York, USA; BSNS, Bufalo Museum of Science, New York, USA; CMNH, Cleveland Museum of Natural History, Cleveland, USA; IRSNB, Institut royal des Sciences naturelles de Belgique, Brussel, Belgium; IVPP, Institute of Vertebrate Paleontology and Paleoanthropology, Beijing, China; MCZ, Museum of Comparative Zoology, Harvard, USA; MCT, Museu de Ciências da Terra, Rio de Janeiro, Brazil; MGL, Natural History Museum of Lille, Nord, France; MV, Museums Victoria, Melbourne, Australia; NHMD, Natural History Museum of Denmark, Copenhagen, Denmark; NHMUK, Natural History Museum, London, UK; NRM, Swedish Museum of Natural History, Stockholm; PMO, Natural History Museum, University of Oslo, Oslo, Norway.

### 2.2. Taxon sampling

The literature and museum collections were surveyed for species-level records of Devonian actinopterygians that are known from more than isolated or fragmentary material (Supplementary Table 1), capturing a total of 29 taxa (including one as-yet-unnamed specimen that likely represents a new genus). These were assessed to determine suitability for inclusion in the study, with ten taxa ultimately excluded: those in which the mandible is not preserved or is substantially incomplete (?*Howqualepis youngorum*, *Pickeringius acanthophorus, Moythomasia perforata*, *Krasnoyarichthys jesseni*); those from deposits where past work indicated insufficient differentiation between bone and matrix in scans (*Cheirolepis schultzei*); those in which X-ray CT was attempted but ultimately unsuccessful due to poor X-ray penetration and/or incomplete preservation (*Donnrosenia schaefferi, Moythomasia lineata, Moythomasia nitida, Cheirolepis canadensis*; Supplementary Table 2); and those in which scanning was not possible due the size or geometry of the specimen or the host rock and/or extremely flat preservation (*Stegotrachelus finlayi*). In the case of *Cheirolepis trailli*, the problem of low contrast between the fossil and host rock was overcome with the use of synchrotron tomography, but this technique could not be used for all taxa due to beamtime limitations.

### 2.3. X-ray computed tomography and 3D models

Specimens of the 19 taxa included in the study were investigated using lab-based X-ray CT (and, in the case of *C. trailli*, synchrotron CT). Regions of interest focussed on the cranium or jaw region, with the aim of capturing as much anatomical detail as possible. For most taxa, a single specimen (or jaw) was examined, although where material was damaged or incomplete more than one jaw was scanned to supplement the description. In most cases, the left lower jaw was targeted, but where the left mandible was not preserved or otherwise unsuitable the right jaw was scanned and mirrored for ease of comparison.

Full details of the specimens and X-ray CT parameters (voltage, current, exposure, number of projections, number of frames per projection, filtration, voxel size) are given in Supplementary Table 3. Where specimens had already been described using X-ray CT, the original tomographic data was downloaded and the mandible resegmented where necessary.

*Cheirolepis trailli* NHMUK PV P 62908b was examined using propagation phase contrast synchrotron X-ray micro-computed tomography at the BM05 beamline of the European Synchrotron Radiation Facility (ESRF, Grenoble, France). The setup of the beamline for the acquisition was: filtered white beam (Mo 3.93 mm) resulting in a total integrated detected energy of 133 keV; indirect detector comprising a LuAG:Ce 2000µm with reflective layer, 0.615x magnification from optical lenses, PCO.edge 4.2 CLHS detector. The measured pixel size on the detector located at 3.5 m downstream from the sample was 10.14 µm. Given the limited field of view with that setup (2048 x 3040 px^2^, i.e., 20.77 x 3.08 mm^2^), an offset of 9.13 mm was applied on the centre of rotation (equivalent to 900 pixels on the detector), and 25 acquisitions were performed moving the sample by 2 mm along the vertical axis between each scan. Each acquisition consisted of 5100 projections of 50 msec total exposure time (5 frames of 10 msec accumulated), 40 dark current images, and 41 flatfield images (i.e., image of the beam with no sample). The tomography reconstruction was done using the single distance phase retrieval approach included in PyHST2 (Paganin et al., 2002; Mirone et al., 2014), generating individual 32-bit tomogram stacks. Post-processing included: merging of multiple stacks into a single one, change of the dynamic range from 32-bits to 16-bits using the 0.001% saturation value provided by the 3D histogram generated with PyHST2, ring correct on individual images (Lyckegaard et al., 2011), cropping of the data. Additionally, a 2×2×2 binning was generated, providing a smaller version of the dataset for rapid inspection.

CT data were exported as .vol files or .TIFF stacks and cropped to remove empty space surrounding the fossil. Datasets were segmented using Materialise Mimics v.25 and v.26 (Materialise Software, Leuven, Belgium; https://www.materialise.com/en/healthcare/mimics-innovation-suite/mimics). The resulting 3D models were exported into Blender v.2.79 (Blender Project; https://www.blender.org/) to create renders for illustrations. Blender was also used to partially retrodeform the mandibles of *Cheirolepis jonesi* PMO 235.121, *Cheirolepis trailli* NHMUK PV P 1370, and *Limnomis delaneyi* ANSP 23721. Both the left and right mandibles of *Cheirolepis jonesi* are broken into several slightly misaligned fragments (Supplementary Figure 1), and these were realigned into life position. The coronoid series in *Cheirolepis trailli* NHMUK PV P 1370 had been displaced slightly ventrally and rotated such that the tooth-bearing surface faced medially (Supplementary Figure 2); this was repositioned to sit on the medial shelf of the dentary. The left mandible of *Limnomis delaneyi* is fractured in several places (Supplementary Figure 3), and these pieces were realigned.

### 2.4. Data availability

The raw data (projections) for the synchrotron tomography scan of *Cheirolepis trailli* NHMUK PV P 62908b are deposited at: doi.org/10.15151/ESRF-DC-2013017890. Projection series (as .TIFF files), tomogram stacks (as .TIFF files), associated scan metadata (multiple formats), and surface meshes for segmented elements (as .PLY files and one combined .OBJ file) of specimens scanned for this study are archived on MorphoSource and ADMorph following institutional and funder policy; temporary Dropbox links are provided for the PLY files during the review process, and will be replaced with stable Morphosource DOIs upon acceptance. Full details are given in Supplementary Table 4.

## 3. RESULTS

### 3.1 Anatomical overview

#### 3.1.1. Terminology and skeletal anatomy

The lower jaw in Devonian actinopterygians is a complex anatomical unit consisting of an outer and inner series of dermal bones surrounding an endochondral ossification. The lateral (labial) surface of the jaw of Devonian actinopterygians is primarily formed from two dermal bones: the dentary, which is generally larger and tooth-bearing, and the edentulous angular, which is typically smaller and confined posteroventrally. Both of these typically bear the mandibular sensory canal and associated pores; a pit-line is often present on the angular. In some taxa, a third dermal bone, the surangular (sometimes called the supra-angular), may be present dorsal to the angular. These latter bones are typically referred to as infradentary bones, or more infrequently as postdentary bones. The posterodorsal corner of the lateral face of the mandible may be slightly depressed where it is overlapped by the maxilla. Ridges of enamel of various length typically ornament exposed surfaces of these dermal bones, although smooth or porous ornament is present in some taxa.

An additional series of dermal bones is present on the medial surface of the lower jaw: an anterior series of coronoids, the number of which may vary, and a prearticular.

Rarely, an additional, more posterior, prearticular plate may be present. The coronoids and prearticular may be wholly or partly fused to each other, with no observable sutures, in which case the number of coronoids cannot be discerned.

These dermal bones surround chondral ossifications that derive from a cartilage known as Meckel’s cartilage, which spans the length of the jaw. In Devonian actinopterygians, this cartilage may be ossified as a single element or as two endochondral bones; in the latter case, the anterior is referred to as the mentomeckelian and the posterior as the articular.

#### 3.1.2. Exemplar description: *Gogosardina coatesi*

The lower jaw of *Gogosardina* was originally described by Choo et al. (2009) based on articulated and semi-articulated acid-prepared specimens. Here we supplement this description based on X-ray CT of MV P.228269 (Fig. 1), a complete left mandible measuring 26 mm in length. We use this high-resolution scan of a well-preserved specimen to provide an expanded description of a Devonian actinopterygian mandible.

**Fig. 1:**
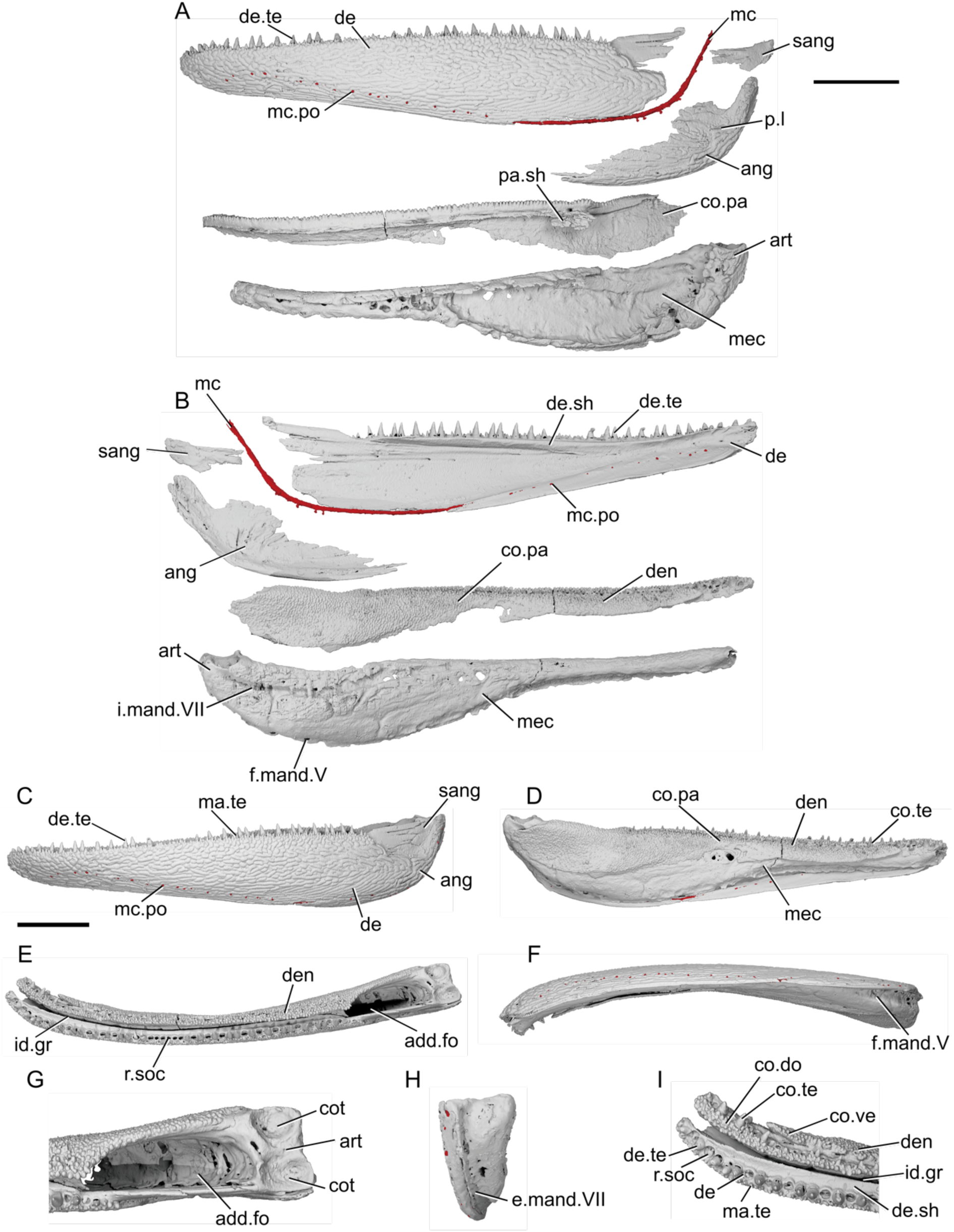
Mandible of *Gogosardina coatesi* MV P.228269. **a**, Left mandible in lateral view, with individual elements separated. **b**, left mandible in medial view, with individual elements separated. **c**, lateral view. **d**, medial view. **e**, dorsal view. **f**, ventral view. **g**, articular region in dorsal view. **h**, posterior view. **i**, anterior region in dorsal view. Scale bar a,b = 5mm. Second scale bar for c-f, h = 5 mm. Panels g, i not to scale. *Abbreviations*: add.fo, adductor fossa; ang, angular; art, articular; co.do, dorsal part of the coronoid; co.pa, coronoid-prearticular; co.te, coronoid teeth; co.ve, ventral part of the coronoid; cot, articular cotyle; de, dentary; de.sh, dentary shelf; de.te, dentary teeth; den, denticles; e.mand.VII, external mandibular branch of the facial nerve; f.mand.V, mandibular branch of the trigeminal nerve; i.mand.VII, internal mandibular branch of the facial nerve; id.gr, interdental groove; ma.te, marginal teeth; max, area overlapped by the maxilla; mc, mandibular canal; mc.po, pores connecting to the mandibular canal; mec, Meckel’s cartilage; p.l, pit line; pa.sh, prearticular shelf; r.soc, replacement socket; sang, surangular.

The dorsal margin of the lower jaw is approximately straight. The mandible is dorsoventrally deepest three-quarters of the way along its anteroposterior length, and gently tapers anteriorly. Posterior to the deepest point, the rear margin curves dorsally. In dorsal and ventral view, the anterior half of the jaw turns gently medially, reaching an angle of approximately 45 degrees from the long axis by the anterior tip of the jaw. Choo et al. (2009) note that the jaw of *Gogosardina* is extremely similar to that of *Mimipiscis* (Gardiner, 1984).

The **dentary** (de, Fig. 1a,b,c,i) is by far the longest element of the lower jaw in *Gogosardina*, spanning approximately 90% of its anteroposterior axis. Along the anterior two thirds of its length, the dentary forms the entire lateral surface of the lower jaw, from the dorsal margin to the ventral margin; the infradentaries contribute to the posterior third. The dentary remains a consistent transverse thickness throughout almost the entire bone. However, the posterodorsal corner is significantly thinner in the region that is overlapped by the maxilla (max, Fig. 1a,c).

The medial surface of the dentary is recessed along most of its length, where it receives Meckel’s cartilage. This depression tapers towards the anterior end. The medial surface also bears a shelf (de.sh, Fig. 1b,i) that originates at the very anterior tip of the jaw and terminates just anterior to the posterior-most teeth. The shelf lies just below the base of the dentary teeth and projects medially with a depth around half the width of the dentary toothrow, other than its anterior and posterior ends, where it rapidly tapers. The shelf is generally very thin, but the thickness bulges for a short section in the middle so that the shelf is pyramidal in cross section. The shelf overlaps Meckel’s cartilage but does not make contact with the dermal tooth-bearing bones on the inner surface of the jaw.

The **angular** (ang, Fig. 1a,b,c) is approximately crescent shaped, and forms the posteroventral margin of the jaw. Anteriorly, the angular forms a distinctly stepped suture with the dentary, resulting from two anterodorsal projections. Anteriorly, the angular has a long, narrow ramus, over half the length of the bone, and extending for almost a third of the total length of the mandible. The anterior portion of the medial surface of the angular has a depression that is continuous with the dentary depression that receives Meckel’s cartilage. Below this, along its ventral margin, the medial surface of the angular is thickened and is deeper than the rest of the bone. This thicker region houses the mandibular canal (mc, Fig. 1a,b).

A small **surangular** (sang, Fig. 1a,b,c) can be identified between the dentary and angular, which was not recognized in the original description (Choo et al., 2009). It is overlapped by both other external dermal elements. It appears to have been interpreted as an anterior flange of the angular by Choo et al. (2009), but is clearly a distinct bone in tomograms. Anteriorly, the surangular interdigitates strongly with the dentary. The surangular is situated entirely within the region overlapped by the maxilla, and completely lacks ornamentation.

Almost the entire lateral surface of the dentary and angular show robust **dermal ornament**. This consists of long, anteroposteriorly directed ridges in the posteroventral region of the jaw, becoming more steeply angled posteriorly. Dorsally and anteriorly, the ornamentation is less regular; the ridges become shorter, are oriented in a greater variety of directions, and close to the dorsal margin the ornamentation resembles randomly arranged tubercles. These appear to grade into the marginal dentition. Tomograms reveal that the ridges forming the ornamentation are mostly hollow. They consist of a series of overlapping, convex bulges of dermal bone housing elongate vacuities, lateral and ventral to the solidly ossified core of dermal bone forming the medial portion of the dentary.

The dentary bears a single row of large teeth (de.te, Fig. 1a,b,c,i) and more numerous marginal teeth (ma.te, Fig. 1c,i). The primary tooth row on the dentary includes 32 preserved teeth, 31 of which are complete and one of which is broken. The teeth are large, sharp, conical and oriented vertically. They are fairly consistent in size along the whole dentary, though the anterior four or five, and posterior two teeth, are smaller than the others. Although these two posteriormost teeth are more comparable in size to the marginal teeth, we identify them as belonging to the primary tooth row as they each have a broad base that appears to be set into a socket.

Replacement pits (r.soc, Fig. 1e,f) lie between most teeth. There is no consistent pattern between teeth and replacement pits: sometimes they alternate, but pits can also be separated by two teeth, three teeth, or a set of four that includes a partially developed tooth. There is also a series of six replacement pits in a continuous line with no teeth. Hypermineralised enameloid caps (acrodin) are difficult to identify in tomograms of this specimen but can be seen on some of the larger teeth.

Lateral to the primary tooth row is a numerous row of much smaller, irregularly sized marginal teeth (ma.te). At their largest, these teeth are barely a quarter of the height of those in the main tooth row. Although they are randomly distributed, no more than two irregular rows of marginal teeth appear to be present before the series grades into the dermal ornament. As with the main dentary tooth row, acrodin is difficult to identify in tomograms but appears to be present on at least the largest teeth.

**Meckel’s cartilage** (mec, Fig. 1a,b,d) is ossified as a single bone spanning the entire anteroposterior length of the jaw. Anteriorly, the ossification is dorsoventrally narrow and subcylindrical. At approximately the midpoint of the jaw, Meckel’s cartilage increases in depth and expands towards the ventral jaw margin, which it reaches in line with the anterior extent of the angular. Posterior to this, Meckel’s ossification closely matches the whole jaw in its full extent and shape. It is exposed medially, ventral to the coronoid and prearticular series, but its lateral extent is entirely covered by the dentary and postdentaries.

Towards the posterior margin of the jaw, Meckel’s ossification increases in width as well as depth and forms the medial surface, posterior margin and floor of the adductor fossa (add.fo, Fig. 1e,g). It also contributes to some of the lower part of the lateral surface of the fossa. Here, Meckel’s cartilage diverges and expands in depth, forming a small dorsal buttress, and a large ventral buttress. Both articulate with the angular.

Posterior to the adductor fossa, the cartilage is robustly ossified and forms the articular region (art, Fig. 1a,b,g), which articulates with the quadrate. The two concave articular cotyles (cot, Fig. 1g) are ovoid, slightly longer than they are wide, and dorsally directed. Almost the entire surface of Meckel’s ossification is covered in perichondral bone, including parts of the Meckelian ossification exposure in the adductor fossa, with unfinished bone restricted to patches in this area. The anterior tip of Meckel’s ossification also lacks perichondral bone, but is instead developed as an ovoid, concave facet that may have articulated with its antimere via a cartilaginous connection.

A deep groove for the internal mandibular branch of the facial nerve (i.mand.VII, Fig. 1g) runs along the medial surface of Meckel’s cartilage, close to, although distinct from, the dorsal margin of the ossification, the path of which it mirrors. It extends from the articular region anterior to the adductor fossa and is roofed by the prearticular. The posteroventral surface of Meckel’s ossification accommodates the external mandibular branch of the facial nerve (e.mand.VII, Fig. 1h) in a broad groove, which fades out approximately in line with the posterior end of the dentary tooth row. Several irregularly spaced foramina are present at the junction between the ventral margin of Meckel’s cartilage and the dermal bones of the jaw, opening into the large, hollow space between Meckel’s cartilage and the lateral and medial dermal bones in the core of the jaw. The posterior two of these foramina reside within the groove for the external mandibular branch of the facial nerve and at least one transmitted the mandibular branch of the trigeminal nerve (f.mand.V, Fig. 1b).

The **inner dermal tooth-bearing bones** extend along the entire length of the medial surface of the jaw. The entire prearticular and coronoid series is fused, with no visible traces of sutures (co.pa, Fig. 1a,b,d). However, in comparable Devonian taxa where sutures remain visible, the prearticular typically forms the posterior half of the series.

The **coronoid** series extends to the very anterior tip of the jaw, mirroring the gentle medial curvature of the dentary. Two primary tooth-bearing surfaces appear to be present: one oriented primarily dorsally and one oriented primarily medially. The dorsal component of the coronoid toothplates (co.do, Fig. 1i) extends to the anterior end of the jaw, but the ventral component (co.ve, Fig. 1i; the “tuberculated lamina” of Gardiner, 1984: p.331) tapers to a gently rounded point a short distance from the anterior end of the jaw. These two regions are separated by a shallow groove that tapers posteriorly, terminating a short distance anterior to the adductor fossa. The coronoid region does not form a lateral shelf. In dorsal view, the tooth-bearing portions of the medial dermal bones are separated from those of the lateral dermal bones by a deep gap (id.gr, Fig. 1e,i), which steadily widens towards the anterior end of the jaw.

The **prearticular** region mainly consists of a large vertical lamina with a slightly convex ventral margin. This lamina is about half of the dorsoventral jaw depth at its deepest point, overlies the dorsal portion of the posterior part of Meckel’s ossification, and extends posteriorly almost to the rear end of the jaw. Dorsally, a narrow horizontal lamina that extends lateral to Meckel’s cartilage and defines the medial and anterior limits of the adductor fossa. At the lateralmost extent of the adductor fossa, the prearticular region forms a short, unornamented horizontal shelf (pa.sh, Fig. 1a) that is overlain by a corresponding horizontal lamina of the maxilla.

Almost the entire dorsal and medial surface of the prearticular and coronoid series is covered with a shagreen of small, blunt, rounded denticles (den, Fig. 1b,d,i). However, the ventral half of what is assumed to be the posteriormost coronoid lacks denticles. Most of these minute denticles are roughly uniform in size, although they are larger on the dorsal surface of the prearticular and coronoids. Additionally, the anterior portion of the dorsal coronoid series bears a row of thirteen larger teeth (co.te, Fig. 1d,i), the largest of which are comparable in size to the primary dentary toothrow. Like the dentary tooth row, replacement pits are interspersed with the larger coronoid teeth, with an uneven distribution as on the dentary. Unlike the dentary tooth row, however, the teeth in the coronoid tooth row are largest anteriorly and decrease in size posteriorly, to the point where they are virtually indistinguishable in size from the surrounding denticles. Three of the anteriormost teeth are distinctly larger and more medially curved than the remainder of the series, resembling miniature fangs. Acrodin appears to be present on at least the larger coronoid teeth.

The **mandibular canal** (mc, Fig. 1a,b) extends through the dentary and angular for almost the entire length of the jaw. It closely follows the posteroventral margin of the angular. Within the dentary, the posterior end of the mandibular canal lies close to the ventral margin. Anterior to this, it rises gently dorsally to reach approximately the dorsoventral midpoint, beyond which it arches and changes direction to run slightly ventrally again. The mandibular canal terminates slightly posterior to the anterior tip of the jaw, in the ventral third of the bone.

The path of the mandibular canal can be traced by a series of pores (mc.po, Fig. 1a,b) that connect to both the inner and outer surface of the jaw. The pores are more numerous and fairly regularly spaced on the lateral face of the dentary (approximately 20 pores) and are more irregularly spaced on the inner face (approximately 14 pores). There are fewer pores on the angular, which are concentrated towards the posterior half of the bone (approximately ten pores on the lateral face). Sensory pores that connect to the canal through the medial face of the dentary lie ventral to Meckel’s cartilage, but those on the angular are level with it.

Additionally, a short, deep pit-line (p.l, Fig. 1a) is present on the lateral surface of the angular, above the mandibular canal though still close to the ventral margin, and just below the posterior extremity of the dentary. This pit-line is not directly connected to the mandibular canal via a distinct canal, but many small tubules are present in this region and some link the pit-line and mandibular canal.

### 3.2 Early Devonian Taxa

#### 3.2.1. Meemannia eos

The isolated left mandible IVPP V14536.5 was assigned to *Meemannia eos* by Zhu et al. (2010) based on its diminutive size (9.4 mm long) and surface ornament of small pores, which is otherwise seen only in partial crania described as *Meemannia eos*. The original description of *Meemannia*, based on external anatomy, noted a peculiar combination of actinopterygian and sarcopterygian characters, and the taxon was interpreted, with caveats, as a stem sarcopterygian (Zhu et al., 2006; 2010). More recent tomographic investigation of the skull roof and partial braincase of *Meemannia* has indicated an actinopterygian affinity (Lu et al., 2016). Consequently, we include the isolated mandible referred to the taxon in our review of actinopterygian lower jaws and redescribe it based on CT data (Fig. 2).

**Fig. 2:**
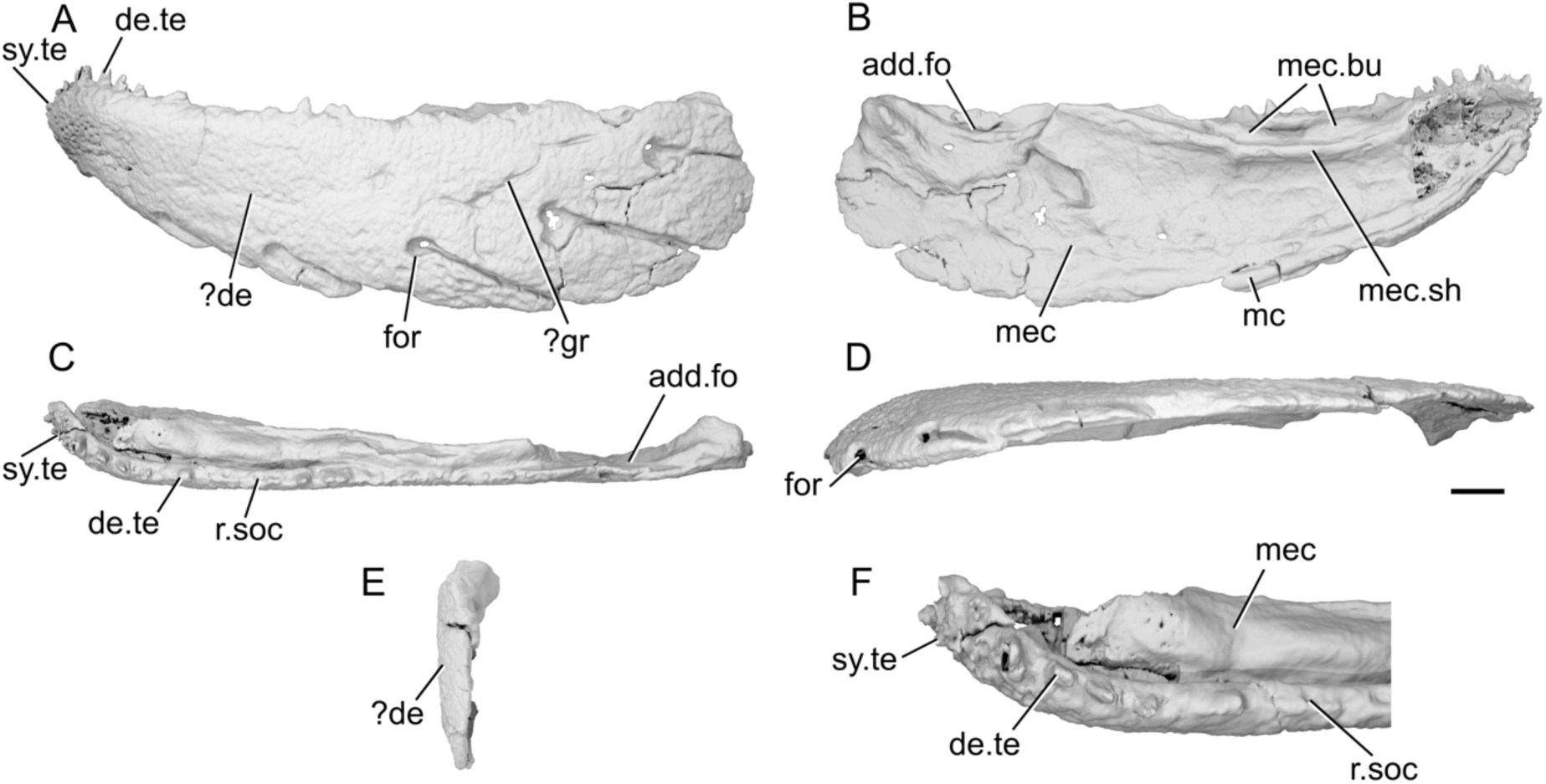
Mandible of *Meemannia eos* IVPP V14536.5. **a**, Left mandible in lateral view. **b**, medial view. **c**, dorsal view. **d**, ventral view. **e**, posterior view. **f**, anterior region in dorsal view. Scale bar = 1 mm. Panel f not to scale. *Abbreviations*: add.fo, adductor fossa; de, dentary. de.te, dentary teeth; for, foramen; gr, groove; mc, mandibular canal; mec, Meckel’s cartilage; mec.bu, bumps on Meckel’s cartilage; mec.sh, shelf on Meckel’s cartilage; r.soc, replacement socket; sy.te, symphysial teeth.

As the entire mandible is tightly fused into a single unit, it is difficult to differentiate separate ossifications. Extensive mechanical preparation has also removed much of the external surface of the dermal bone and perichondrium, including around the sensory pores. Consequently, the extent of many ossifications and the number and shape of teeth is difficult to determine.

The mandible is relatively stout, and approximately three times longer than it is deep. The dorsal margin is very gently concave along its entire length, though slightly more so anteriorly. However, the very anterior tip of the jaw is directed anteroventrally. The ventral margin is more strongly curved than the dorsal margin and the jaw tapers anteriorly, but there is no distinct symphysial reflex. The mandible is fairly straight in dorsal view, and curves gently medially at the anterior end.

It is not possible to confidently identify individual dermal bones contributing to the external surface. A groove in the posterior half of the jaw (?gr, Fig. 2a) descends anteroventrally from the midpoint of the margin of the adductor fossa for approximately a quarter of the length of the jaw, at which point it bifurcates. It is possible that this represents a suture between the dentary and the postdentary series, but as mechanical preparation has penetrated through the matrix and into the bone it is unclear whether this is an original feature or artefact.

Five large foramina (for, Fig. 2a,d) are arrayed along the lateral surface of the jaw. A sixth, anterior foramen notches the ventral margin of the jaw, but has been partially worn away either by preservation or preparation. Although originally described as infradentary foramina (Zhu et al., 2010), it is unclear which dermal ossification these features are borne in. Each more posterior foramen is positioned more dorsally. A narrow, deep groove extends from the posteroventral edge of each foramen to the ventral margin of the mandible, terminating in a thickened ridge that transmitted the mandibular canal (mc, Fig. 2b). This ridge is incomplete posteriorly due to breakage of the specimen. The mandibular canal exits the posterior margin of the mandible approximately midway between the dorsal and ventral margins of the jaw.

Meckel’s cartilage (mec, Fig. 2d,f) appears to be ossified as a single element along the whole length of the jaw. It is closely associated with the lateral dermal bone, but can be identified as chondral bone in tomograms. Only patches of perichondral bone are preserved on the surface of Meckel’s ossification, as some of the thin ossification on the medial surface has been prepared away. Therefore, it cannot be concluded how extensive the perichondral covering originally was. At the posterior extremity of Meckel’s ossification, the adductor fossa (add.fo, Fig. 2b,c) is developed as an elongate depression. Its posterior, lateral and anterior margins are formed by Meckel’s cartilage, which initially increases in height anterior to the fossa before flattening medially to form a horizontal shelf. Its medial extent is not preserved and is not possible to identify the position or orientation of the articular cotyles.

There is no preserved evidence of medial dermal bones, although the dorsal surface of Meckel’s cartilage is developed as a series of low, rounded bumps (mec.bu, Fig. 2b) with a medially directed shelf ventral to the bumps (mec.sh, Fig. 2b). Zhu et al. (2010) interpreted the coronoids as sitting on the dorsal surface of this shelf and the prearticular resting ventral to it, resembling the condition seen in generalized sarcopterygians (Zhu & Yu, 2004). The series of elongate, low, rounded bumps that were originally suggested to separate the coronoid series recall those in the stem osteichthyan *Megamastax* (Choo et al., 2014). Current literature interprets these features as fused coronoid toothplates that facilitated a durophagous ecology (Choo et al., 2014).

A number of moderately large, vertically oriented teeth (de.te, Fig. 2a,c,f) are preserved in a single row on the dorsal surface of the mandible. Nine distinct teeth can be observed, but several more were likely present as preservation and preparation have resulted in substantial damage and wear to the toothrow. The teeth generally increase in size anteriorly, on the symphysis, the toothrow becomes directed medially and ventrally and the teeth decrease in size (sy.te, Fig. 2a,c,f). The anterior seven teeth are positioned close together, followed by a substantial gap with three empty sockets (r.soc, Fig. 2c,f), before an eighth tooth. This suggests that tooth replacement did not occur with regularity. Marginal teeth appear to be absent along the primary tooth row, although a broad field of irregular denticles are visible at the anteriormost tip of the mandible.

### 3.3 Middle Devonian Taxa

#### 3.3.1. Cheirolepis trailli

The description of the mandible of *Cheirolepis trailli* is based on two specimens: NHMUK PV P 62908b (Fig. 3a-d), examined via synchrotron tomography and preserving a nearly complete right lower jaw (preserved length 42 mm); and NHMUK PV P 1370 (Fig. 3e-l; Fig. S1), examined via X-ray CT, preserving a near-complete right lower jaw (preserved length 33 mm) that has undergone some lateral compression, as well as a left lower jaw from which the articular can be segmented. Both specimens are preserved across part and counterpart, with only the part subject to tomography, and as a result some details of the external surface of both cannot be observed. A detailed description of the lower jaw of the taxon is known from external observation of several specimens (Pearson & Westoll, 1979).

**Fig. 3:**
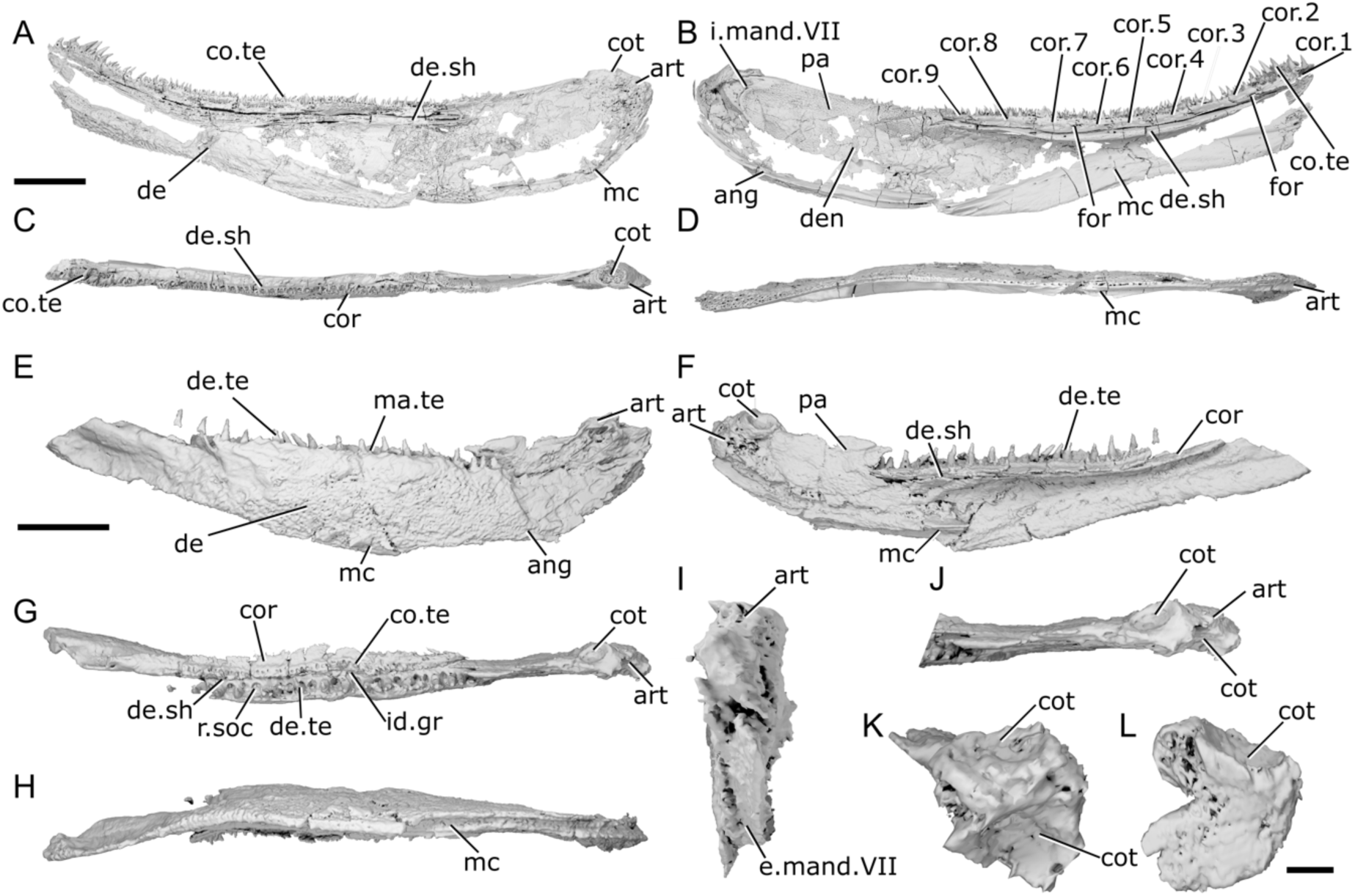
Mandibles of *Cheirolepis trailli*. **a**, Right mandible (mirrored) of NHMUK PV P 62908b in lateral view. **b**, NHMUK PV P 62908b in medial view. **c**, NHMUK PV P 62908b in dorsal view. **d**, NHMUK PV P 62908b in ventral view. **e**, Right mandible (mirrored) of NHMUK PV P 1370 in lateral view. **f**, NHMUK PV P 1370 in medial view. **g**, NHMUK PV P 1370 in dorsal view. **h**, NHMUK PV P 1370 in ventral view. **i**, NHMUK PV P 1370 in posterior view. **j**, NHMUK PV P 1370 articular region in dorsal view. **k**, NHMUK PV P 1370 left articular in dorsal view. **l**, NHMUK PV P 1370 left articular in lateral view. Scale bar: a,b,c,d = 5 mm. e-h = 5 mm. k, l = 1 mm. Panels i, j not to scale. *Abbreviations*: ang, angular; art, articular; co.te, coronoid teeth; cor, coronoid; cot, articular cotyle; de, dentary; de.sh, dentary shelf; de.te, dentary teeth; den, denticles; e.mand.VII, external mandibular branch of the facial nerve; for, foramen; i.mand.VII, internal mandibular branch of the facial nerve; id.gr, interdental groove; ma.te, marginal teeth; mc, mandibular canal; pa, prearticular; r.soc, replacement socket.

The ventral margin is strongly convex in the posterior half, straight in the midsection, and rises towards the anterior extent of the mandible. Similarly, the dorsal margin is gently concave in its posterior third, flat in the midsection, and dorsally inclined anteriorly. This results in a curved profile to the jaw.

Most of the lateral surface of the mandible comprises the dentary (de, Fig. 3a,e), which is ornamented with small pore openings, as well as ridges more ventrally. An angular (ang, Fig. 3b,e) is present in both specimens, though it is incomplete and fragmentary in NHMUK PV P 1370. In neither specimen can the suture between the angular and the dentary be observed easily on the lateral surface, but it is evident in tomograms. The angular is narrow and rod-like, thickened medially where it carries the mandibular canal, and is restricted to the posteroventral margin of the jaw, projecting anteriorly as far as the posterior end of the toothrow. A surangular is more difficult to identify in tomograms of these two specimens due to the mode of preservation, but is apparent on the counterpart of NHMUK PV P 62908b (NMS 1877.30.5), and has been identified on other specimens (Pearson & Westoll, 1979).

Dorsally, the dentary possesses a substantial medial shelf (de.sh, Fig. 3a,b,c,f) that extends underneath and medial to the coronoids. The shelf is thickest approximately at the midpoint of the jaw. The ventral portion of the dentary is thickened into a ridge enclosing the mandibular canal (mc, Fig. 3a,b,d,e,f,h); this arches dorsally in the anterior half of the mandible, and its path is marked by small pores that connect to both the lateral and medial surface of the dentary.

The dentary toothrow is only preserved in NHMUK PV P 1370 and consists of a single row of numerous large, sharp teeth that curve slightly inwards. The teeth (de.te, Fig. 3e,f,g) are densely spaced, with a small number of replacement sockets (r.soc, Fig. 3g) relative to the number of cusps. They are separated from the lateral margin of the dentary by a small ridge. A row of small, medially curved cusps atop this ridge indicates the presence of a marginal toothrow (ma.te, Fig. 3e), the presence of which is corroborated by existing descriptions (Pearson & Westoll, 1979).

Meckel’s cartilage is ossified only in the articular region (art, Fig. 3a,c,d,e,f,g,i,j), and is c-shaped in lateral view. The cartilage is endochondrally ossified, surrounded by a delicate perichondral shell that has been laterally crushed to a greater or lesser extent in both specimens. Only the medial half of the articular, accommodating one cotyle, is preserved in NHMUK PV P 62908b; the cotyle is ovoid, faces dorsally, and situated some way anterior to the posterior margin of the articular. Both the left and right articular are completely preserved in NHMUK PV P 1370, although the right articular—along with the whole of the posterior region of the mandible—has been somewhat laterally compressed. Both cotyles (cot, Fig. 3a,c,f,g,j,k,l) are preserved on both articulars, and the left articular clearly shows the arrangement of the cotyles. The more lateral cotyle is round and faces dorsolaterally. Its posterior margin is almost level with the posterior margin of the articular. The more medial cotyle is positioned anterior and dorsal to the lateral cotyle, and oriented anterodorsally. This morphology is corroborated by the less compressed left articular of NHMUK PV P 1370. A foramen for the internal mandibular branch of the facial nerve (i.mand.VII, Fig. 3b) is present in NHMUK PV P62908b a short distance below the medial cotyle, adjacent to the very posterior tip of the posterodorsal corner of the prearticular. A shallow groove, accommodating the external mandibular branch of the facial nerve (e.mand.VII, Fig. 3i), is evident in NHMUK PV P 1370, where it runs along the posterior margin of the jaw and continues below the prearticular, following a path that runs parallel to the prearticular ventral margin. This groove terminates anteriorly in line with the posterior end of the dentary toothrow.

A series of coronoid plates and a prearticular line the medial surface of the mandible. The coronoid series consists of many small elements each comprising an extensive medial shelf with a slightly concave base and a raised, thickened lateral portion. Each coronoid (cor, Fig. 3b,c,f,g) lies atop the robust medial shelf of the dentary (de.sh, Fig. 3a,b,g,f,g), separated from it by a narrow gap. Nine coronoid elements can be identified in NHMUK PV P62908b, articulating with one another via interdigitating sutures, and at least five in NHMUK PV P 1370, in which sutures between elements are more difficult to identify. A series of elliptical foramina (for, Fig. 3b) pierce the coronoid series, often intercepting sutures between elements. Both the anterior and posteriormost coronoids have a different morphology to the remainder in the series: the posteriormost (cor.9, Fig. 3b) is narrower, and the anteriormost (cor.1, Fig. 3b) is broader, assuming an oval shape that expands towards the midline and curves somewhat ventrally. The coronoids possess an extensive series of teeth on their thickened lateral portion (co.te, Fig. 3a,b,c,d). These teeth are sharp, conical, curve strongly medially, and are approximately half the height of those on the dentary. The dental field is extremely dense, with a single row of larger medial teeth flanked by one-two rows of smaller, sharper lateral teeth. Only a small number of sporadically placed replacement sockets are apparent, most of which are in the posterior half of the series. On the anterior coronoid, the seven or so teeth (plus one empty socket) in the medial row are considerably expanded in size, resembling small fangs, and curve more sharply medially and slightly posteriorly. The surface of each coronoid is smooth medial to the tooth rows, except for irregularly shaped and spaced bumps, which tend to have a narrow neck and broadened and flattened head in section.

The prearticular (pa, Fig. 3b,f) is flat and ovoid, tapering anteriorly, with a rounded posterior margin and straight dorsal margin. It forms the medial margin of the adductor fossa and is deepest ventral to this region, where it is a little over half the depth of the jaw. Minute, flattened denticles (den, Fig. 3b), some of which are almost stellate in shape and resemble dermal ornament, cover the entire face of the prearticular, except for a narrow rim around the posterior margin. These denticles become larger and more pointed dorsally, and at the anterodorsal extent of the prearticular are elongate relative to their width and curve posteriorly, resembling miniature teeth. Anteriorly, the prearticular overlaps a narrow portion of both the posteriormost coronoid and the dentary shelf.

#### 3.3.2. Austelliscus ferox

The only known specimen of *Austelliscus ferox* is MCT890-P, an incomplete left lower jaw (preserved length 70 mm) preserving part of the dentary as a mould. The anterior region of the jaw is complete, but the mandible is broken anterior to the adductor fossa. It was described in detail using X-ray CT by Figueroa et al. (2021), and the description is supplemented here based on reexamination of these surface files (Fig. 4).

**Fig. 4:**
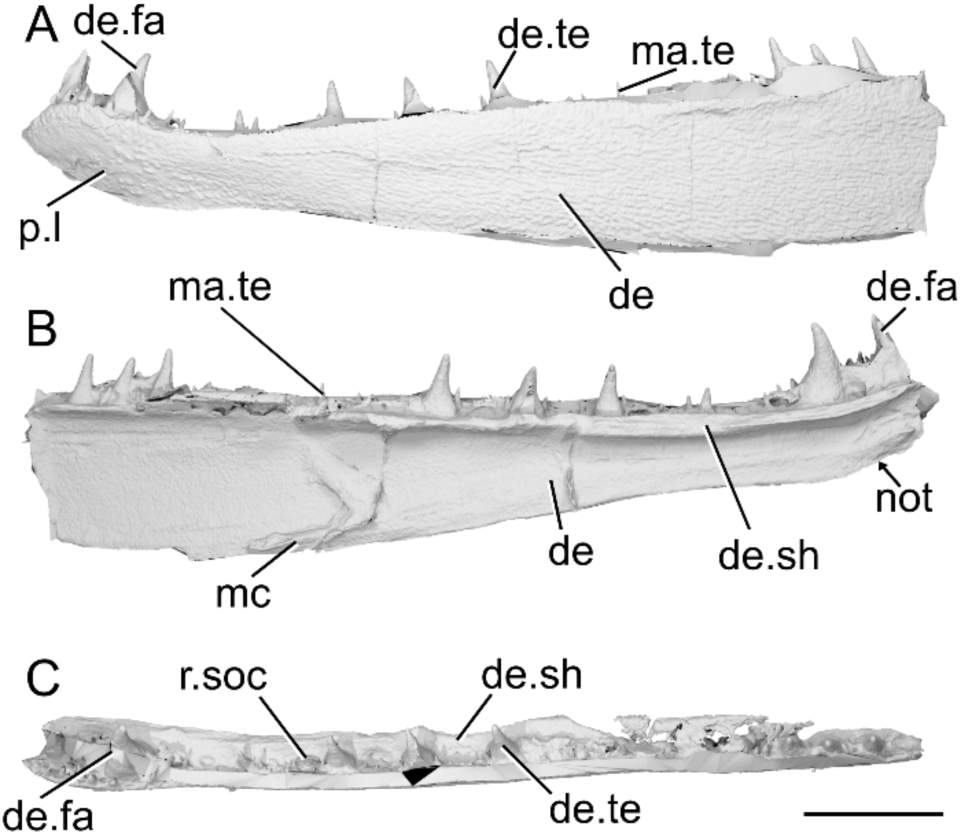
Mandible of *Austelliscus ferox* MCT890-P. **a**, Left mandible in lateral view. **b**, medial view. **c**, dorsal view. Scale bar = 10 mm. *Abbreviations*: de, dentary; de.fa, dentary fang; de.sh, dentary shelf; de.te, dentary teeth; ma.te, marginal teeth; mc, mandibular canal; not, notch; p.l, pit line; r.soc, replacement socket.

The dentary (de, Fig. 4a,b) is relatively shallow, with a dorsal margin that is generally straight and horizontal. A gently concave region is present on the ventral margin with the apex facing dorsally, beginning immediately posterior to the second dentary tooth, and terminating at the third. Posterior to this, the dentary margin is straight and slopes gently. This results in a reflexed anterior margin. A long, well-developed horizontal shelf (de.sh, Fig. 4b,c) extends medially from near the dorsal margin of the dentary, and may have supported the coronoid series. Almost the entire preserved lateral surface of the dentary is ornamented by short, low ridges and grooves, which generally extend anteriorly to posteriorly. In the anterior region of the jaw, where the dentary increases in depth, the ridges give way to randomly arranged tubercles. In this region, a series of approximately five pores forms a short pit-line (p.l, Fig. 4a). The pit-line is crescent shaped, with the apex directed dorsally, and indicates the anterior position of the mandibular canal.

The dentary bears two distinct toothrows. Medially, a series of eight very large, sharp, recurved teeth (de.te, Fig. 4a,c) sit on the horizontal shelf; consequently, their bases are positioned ventral to the dorsal margin of the dentary. The teeth are arranged in groups: as a pair anteriorly, then two sets of three at the midpoint and posterior end of the preserved section of dentary. Within each group, antero-posteriorly elongate replacement sockets (r.soc, Fig. 4c) alternate with the teeth, and several consecutive sockets separate the groups. The anterior two teeth are considerably larger than the other six, are more medially curved with the tip oriented somewhat posteriorly, and appear fang-like (de.fa, Fig. 4); the remaining teeth are less strongly recurved, and slope slightly anteriorly. Lateral to the large toothrow is an additional row of smaller teeth (ma.te, Fig. 4a,). These teeth are conical, vary in size (but are always smaller than those of the primary toothrow), and are irregularly spaced. Posteriorly, this row appears to merge into the larger tooth row.

A series of low ridges, suggested by Figueroa et al. (2021) as marking the course of the mandibular canal (mc, Fig. 4b), are present along much of the medial face of the dentary. Posteriorly, the canal is positioned close to the ventral margin of the jaw, then rises to the mid-height of the jaw in the anterior third. Close to the anterior end of the jaw, the canal arches dorsally, meeting the crescent-shaped pit line on the lateral surface.

#### 3.3.3. Howqualepis rostridens

The lower jaw of *Howqualepis rostridens* was originally described by Long (1988) based on external examination of mouldic fossils and positive copies cast in moulds. Here we redescribe it based on CT scans of two specimens; AMF65495 (Fig. 5a), which includes a mold of the lateral surface of a well-preserved right mandible (43 mm long); and MV P.160801, Fig. 5b-k), which is also mouldic (although some original material remains within the matrix), and contains a complete left (79 mm long) and partial anterior right mandible, both split across part and counterpart.

**Fig. 5:**
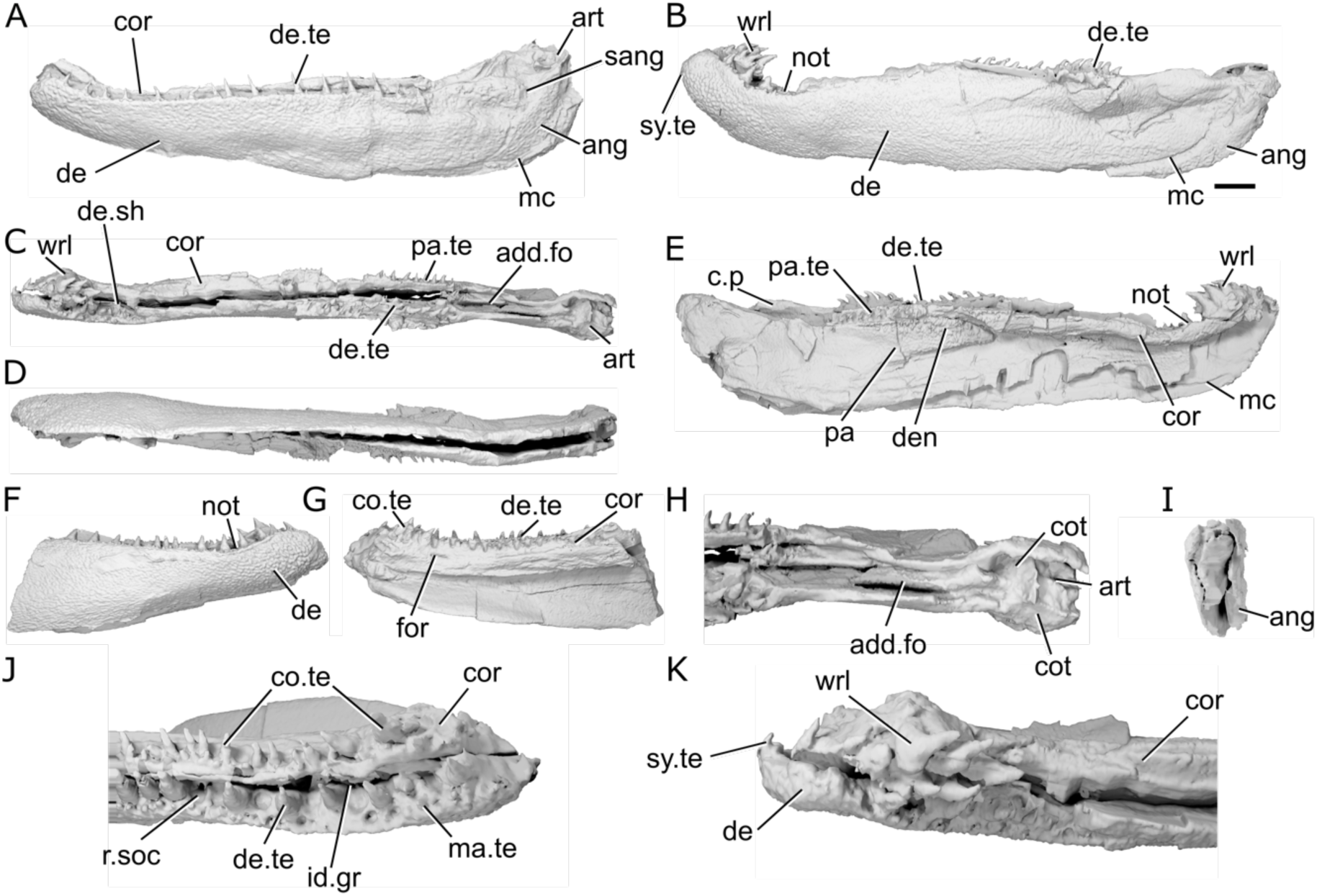
Mandibles of *Howqualepis rostridens*. **a**, Right mandible (mirrored) of AM F 65495 in lateral view. **b**, reconstructed left mandible of MV P.160801 in lateral view. **c**, MV P.160801 left mandible in dorsal view. **d**, MV P.160801 left mandible in ventral view. **e**, MV P.160801 left mandible in medial view. **f**, right mandible of MV P.160801 in lateral view. **g**, MV P.160801 right mandible in medial view. **h**, MV P.160801 articular region of left mandible in dorsal view. **i**, MV P.160801 left mandible in posterior view. **j**, MV P.160801 anterior region of right mandible in dorsal view. **k**, MV P.16080 anterior region of left mandible in dorsal view. Scale bar a-g, i 5 mm. Panels h, j, k not to scale. *Abbreviations*: add.fo, adductor fossa; ang, angular; art, articular; c.p, coronoid process; co.te, coronoid teeth; cor, coronoid; cot, articular cotyle; de, dentary; de.te, dentary teeth; de.sh, dentary shelf; den, denticles; id.gr, interdental groove; ma.te, marginal teeth; mc, mandibular canal; not, notch; pa, prearticular; pa.te, prearticular teeth; r.soc, replacement socket; sang, surangular; sy.te, symphysial teeth; wrl, whorl-like teeth.

The ventral margin of the jaw is slightly convex along its entire length. The dorsal margin is straight along most of the jaw, but has an overall concave profile as it turns upwards at both the anterior and posterior ends of the jaw. A slight notch (not, Fig. 5b,e,f) on the dorsal margin of the dentary immediately precedes the upturn on both mandibles of MV P.160801. This results in the jaw expanding in depth posteriorly, and being oriented anterodorsally at the anterior end.

The dentary (de, Fig. 5a,b,f) occupies almost the entire lateral surface of the jaw. It possesses a medial shelf (de.sh, Fig. 5c) that appears to underly at least the lateral part of the medial dermal tooth-bearing bones, although the full medial extent of the shelf is unknown due to the mouldic preservation. An interdentary groove (id.gr, Fig. 5j) appears to be present between the dentary toothrow and the medial dermal bones.

A crescentic angular (ang, Fig. 5a,b,i) is present, with a long, thin anterior ramus that extends to the anterior margin of the adductor fossa and a thicker posterodorsal ramus. It reaches the posterior end of the jaw, but does not reach as high as the dorsal margin of the articular. A small surangular (sang, Fig. 5a) is present. It is mostly visible in medial view, where it projects dorsally slightly above the level of the adductor fossa to form a modest coronoid process (c.p, Fig. 5e).

Most of the lateral surface of the jaw does not show a distinct pattern of ornamentation. Instead, it is rugose and almost entirely covered by shallow pits. This covering is continued into the posterodorsal region of the jaw that would have been overlapped by the maxilla, which is barely depressed. Ridges of ornament parallel to the long axis of the mandible are present on the posterodorsal ramus of the angular.

The mandibular canal (mc, Fig. 5a,b,e) follows the ventral margin of the angular and dentary in the posterior half of the mandible before rising gently dorsally. Close to the anterior margin, a short series of closely vertically spaced pores are present, indicating that at the very anterior tip of the jaw the mandibular canal is directed downwards. This results in a curved shape for the anterior section of the canal, with the apex of the curve directed dorsally.

The dentary bears a row of large, sharp, pointed teeth (de.te, Fig. 5a,b,c,d,g,j) that curve medially. AM F 65495 shows the entire toothrow, with nineteen teeth present, although individual tooth morphology is not as clear as in MV P.160801. Teeth are present along the entire surface of the jaw, including in the notch preceding the dorsal upturn of the mandible. The teeth generally form a consistent pattern alternating with replacement tooth sockets (r.soc, Fig. 5j), but one set of two and one set of three adjacent teeth are present. At the anterior extremity of the jaw, the tooth-bearing surface of the dentary thickens medially and faces posteriorly, and consequently the teeth in this region are also oriented somewhat posteriorly. Across the jaw, teeth are generally largest in the mid-region and become smaller anteriorly and posteriorly. The teeth are larger posteriorly, and smaller anteriorly, though there are two smaller, partially erupted teeth in the middle of the toothrow. Much of the toothrow in the left jaw of MV P.160801 is poorly preserved, but at the posterior end of the dentary, immediately anterior to the adductor fossa, are six adjacent, small teeth that are strongly recurved.

Lateral to the main tooth row, an additional row of smaller teeth (ma.te, Fig. 5j) lie directly on top of the ridge formed by the dorsolateral margin of the dentary. These teeth are fairly straight and pointed, have dorsally directed tips. Empty replacement sockets are sporadically present along the row. At the anteriormost tip of the left mandible of MV P.160801, on the jaw symphysis, a single medially-directed tooth (sy.te, Fig. 5b,k) is present. A narrow interdental groove (id.gr, Fig. 5j) is present between the tooth-bearing portions of the dentary and medial dermal series.

The medial surface of the jaw is only completely preserved on the left jaw of MV P.160801. The coronoids and prearticular are ossified separately, although the number of coronoids cannot be determined. Both consist predominantly of a medially facing shelf, with a flat crest level with the middle of the dentary teeth forming the dorsal surface.

Although individual coronoids (cor, Fig. 5a,c,e,g,j) cannot be separated, the series as a whole is shallow and reaches the anterior tip of the jaw. The anteriormost coronoid thickens medially anterior to the dorsal upturn of the mandible, and a narrow, elongate foramen is present near its posteromedial margin. The prearticular (pa, Fig. 5e) is elongate and deep, occupies the medial surface of the posterior half of the jaw, and tapers anteriorly to terminate in a triangular point that extends beneath the posterior coronoid. A narrow, curved groove separates the posterior coronoid from the prearticular.

A complex arrangement of teeth and denticles covers the dorsal and medial surfaces of the coronoids and prearticular. One to two rows of small, sharp teeth run close to the dorsal margin of the series (co.te, pa.te, Fig. 5c,e,j), with a larger row positioned immediately medial to them. All of these teeth have medially oriented cusps, and empty replacement sockets are infrequently present. A groove ventral to the larger row separates the dorsal and medial surfaces of the medial dental series. Beneath the groove, a field of variably sized denticles (den, Fig. 5e) is present across much of the remainder of the prearticular. These denticles are typically larger dorsally and decrease in size ventrally, fading out towards the lower margins of the prearticular.

In both jaws of MV P.160801, the anteriormost coronoid teeth are enlarged. The anteriormost coronoid region of the left mandible of MV P.160801 bears three rows of teeth, each of which is arranged antero-posteriorly, resembling a whorl (wrl, Fig. 5b,c,e,k). The teeth in each row are strongly posteriorly curved, with their tips facing posteriorly, and generally increase in size posteriorly. Teeth in the most medial row are substantially larger and more robust than those in the central row, which are again larger than those in the lateralmost row. This whorl was originally considered part of the dentary tooth row by Long (1988).

Meckel’s cartilage is ossified only in the articular region (art, Fig. 5a,c,h), and is partially preserved in AMF65495 and complete in the left jaw of MV P.160801. Two cotyles (cot, Fig. 5h) are present. The more medial cotyle is positioned anteriorly, with a rounded surface that faces dorsally and slightly medially. The lateral cotyle is harder to identify due to the mouldic preservation of the specimen, but appears to face dorsally and be slightly more elongate, as well as positioned more dorsal and posterior to the lateral cotyle. The shape of the adductor fossa (add.fo, Fig. 5c,h) is difficult to discern due to the crushed nature of MV P.160801, but it was clearly narrow and elongate, with parallel lateral margins.

#### 3.3.4. Cheirolepis jonesi

Both jaws of *Cheirolepis jonesi* have been previously described via X-ray CT of PMO 235.121 and external description of other specimens by Newman et al. (2021). We supplement this existing description by re-examining both mandibles (left mandible preserved length: 55 mm; right mandible preserved length: 53 mm) from the existing tomograms and partially retrodeforming the right mandible (Fig. 6; Fig. S2).

**Fig. 6:**
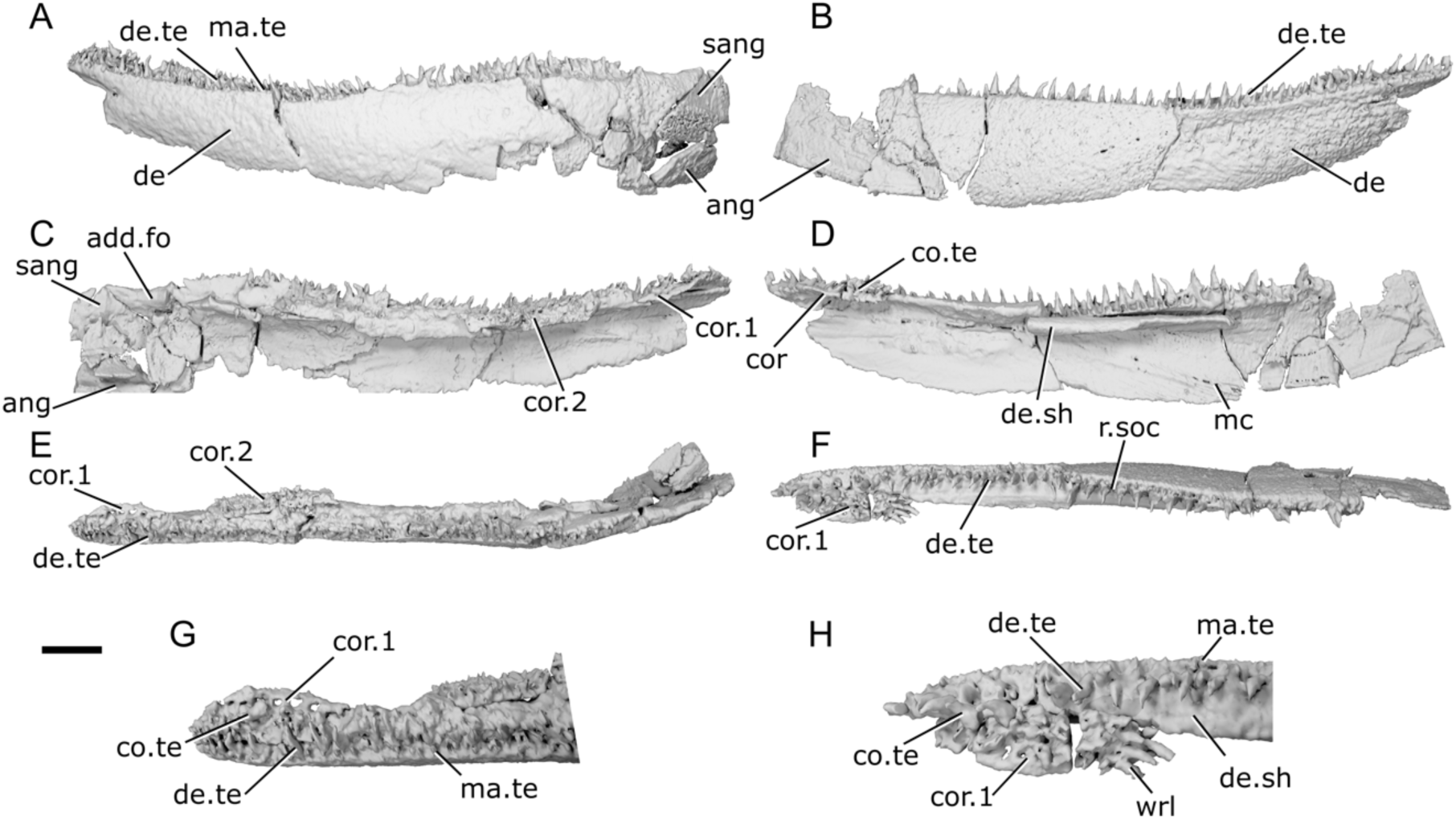
Mandibles of *Cheirolepis jonesi* PMO 235.121. **a**, Left mandible in lateral view. **b**, right mandible in lateral view. **c**, left mandible in medial view. **d**, right mandible in medial view. **e**, left mandible in dorsal view. **f**, right mandible in dorsal view. **g**, anterior region of left mandible in dorsal view. **h**, anterior region of right mandible in dorsal view. Scale bar a-f = 5 mm. Panels g,h not to scale. *Abbreviations*: add.fo, adductor fossa; ang, angular; co.te, coronoid teeth; cor, coronoid. de, dentary; de.sh, dentary shelf; de.te, dentary teeth: ma.te, marginal teeth; mc, mandibular canal; r.soc, replacement socket; sang, surangular; wrl, whorl-like teeth.

The ventral margin of the jaw is gently convex along its whole length. It is curved slightly more strongly in the anterior half. The dorsal margin bulges slightly in the posterior half—although the articular region is not preserved and its height is not known—and is concave in the anterior half. At the anterior end of the mandible, the dentary tapers rapidly towards the toothrow. This results in a jaw profile that curves upwards towards the anterior end. Some ornamentation is evident, particularly in the anterior half of the right jaw, and appears to comprise both ridges and pores. However, the resolution of the scan does not allow for a more detailed description.

The dentary (de, Fig. 6a,b) makes up the majority of the lateral surface of the jaw. Its dorsolateral surface is developed as a narrow vertical ridge, with all teeth positioned medial to this ridge. A robust medial dentary shelf (de.sh, Fig. 6d,h) is present below the level of the toothrow. The shelf is well preserved on the right dentary, displaying a slightly convex dorsal margin that peaks in depth in the middle of the shelf and thins medially. This shape corresponds to the chevron-shaped base of the coronoids. However, the mode of preservation is such that the medial shelf of the dentary and the coronoids of the left jaw cannot be differentiated.

Two infradentaries are present at the posterior end of the jaw. The large, plate-like angular (ang, Fig. 6a,b,c) is partially overlapped by the dentary. It forms all the posterior and a third of the ventral margin of the mandible; viewed medially, where more of it is exposed, it extends for almost half the length of the jaw. The dorsal margin of the angular curves to underlie the medial dentary shelf. A small, wedge-shaped surangular (sang, Fig. 6a,c) is present on the lateral surface of the mandible, close to its posterodorsal margin. It partially overlaps the angular and has an unornamented region where it is overlapped in turn by the maxilla.

The dentary possesses a toothrow of many narrow, tall and sharp teeth that curve medially towards their tips (de.te, Fig. 6a,b,e,f,g,h). These teeth are very densely spaced; replacement sockets (r.soc, Fig. 6f) are present occasionally, but teeth are generally adjacent to one another. Marginal dentition (ma.te, Fig. 6a,g,h) is also present and densely spaced, such that it is difficult to tell whether it is arranged in one or two rows. As is typical for marginal dentition, the teeth are notably smaller than the main dentary teeth, but the marginal teeth are relatively large, reaching half the height of the main dentary teeth in many cases.

Coronoid elements (cor, Fig. 6c,d,e,f,g,h) are associated with the right mandible, but have become separated from the dentary shelf and their exact number is thus difficult to determine. As previously mentioned, the left coronoids cannot be differentiated from the dentary, with the exception of the anteriormost coronoid. A corresponding element is also partially articulated with the anterior end of the right mandible. This anteriormost coronoid (cor.1, Fig. 6c,d,e,f,g,h) is elongate and ovoid, tapering both anteriorly and posteriorly. It bears many sharp, posteriorly recurved teeth arranged in anteroposterior rows that resemble a whorl (wrl, Fig. 6h) posteriorly. Empty sockets are preserved in alignment on the left anterior coronoid, and a narrow, non-denticulated strip runs along the medial edge of this coronoid. At least two additional coronoid elements are preserved, restricted to the anterior portion of the right jaw; comparison with *Cheirolepis trailli* (Fig. 3) and the length of the gap suggests the true number was likely higher. The coronoids are entirely restricted to above the medial shelf of the dentary and their dorsal surface is covered in small, sharp teeth. These teeth are arranged somewhat randomly, rather than being arranged into a row, and are curved medially. Nothing is known of the prearticular, as preservation of the posterior and medial part of the jaw is particularly poor. Similarly, no ossifications of Meckel’s cartilage can be observed, although the lateral margins of the adductor fossa are indicated on the medial surface of the left jaw (add.fo, Fig. 6c).

The mandibular canal (mc, Fig. 6d) originates posteriorly in the angular, at the ventral margin of the jaw. Its path can be traced via a distinct bulge on the medial surface of the mandible, with a few small foramina opening onto both the lateral and medial surfaces. As the mandibular canal moves anteriorly, it enters the dentary and immediately deviates diagonally upwards. Around one-third of the way along the dentary it plateaus, continuing along a straight path to the anterior end of the jaw.

### 3.4 Late Devonian Taxa

#### 3.4.1. Mimipiscis toombsi

The description of the mandible of *Mimipiscis toombsi* is primarily based on X-ray CT of NHMUK PV P 56495 (Fig. 7a,b,d-h), a complete, acid-prepared mandible (16 mm long). A second specimen, NHMUK PV P 53249 (Fig. 7c) an articulated acid-prepared cranium; mandible length 25 mm), was also imaged using X-ray CT, but the lower quality of the tomography data means that many finer features are not visible, and this specimen is only referred to when necessary to highlight variation between the specimens. The mandible of *M. toombsi* has previously been extensively described by Gardiner (1984) based on external description of acid-prepared specimens, as well as more recently by Choo (2012).

**Fig. 7:**
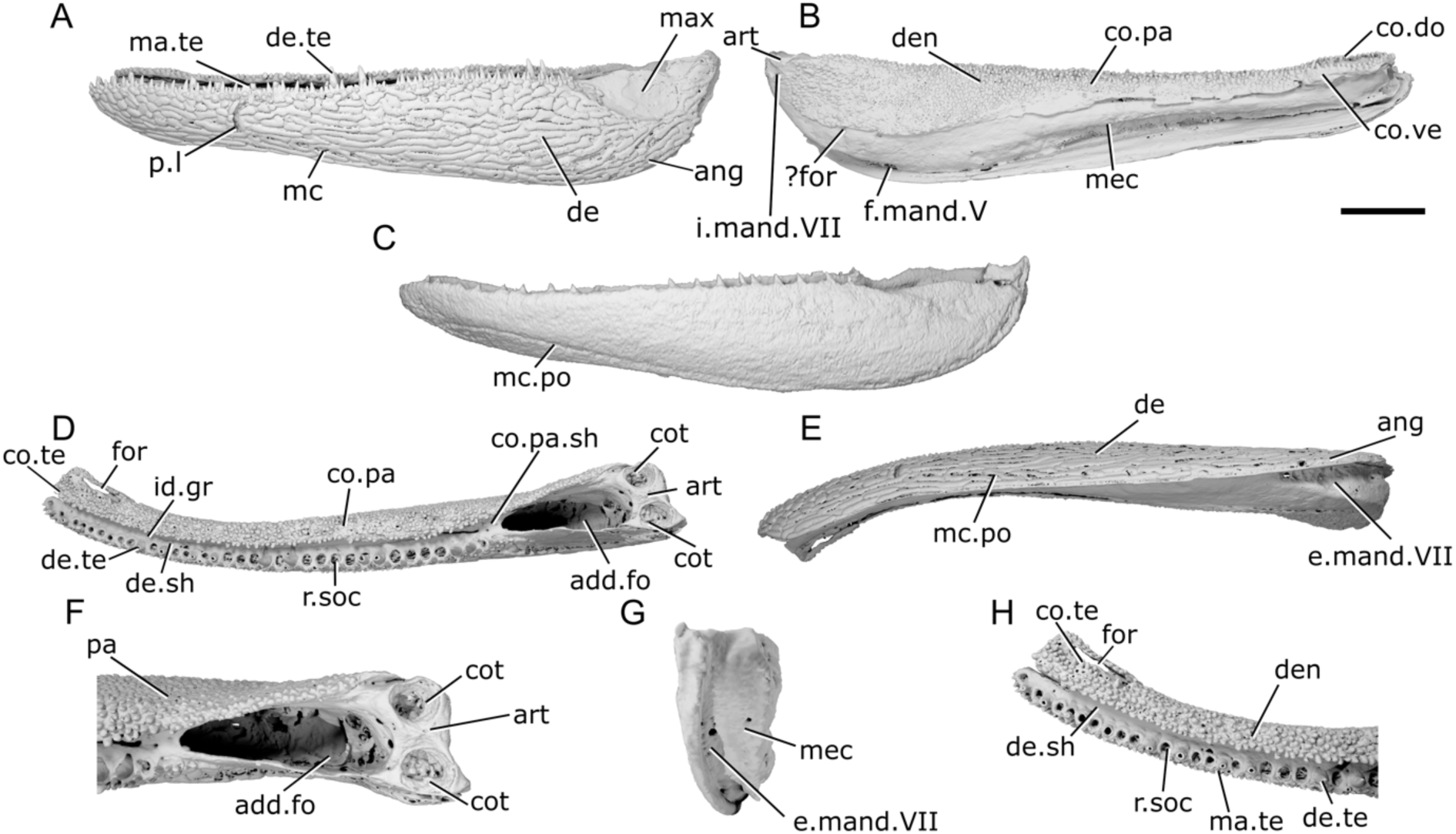
Mandibles of *Mimipiscis toombsi.* **a**, Right mandible (mirrored) of NHMUK PV P 56495 in lateral view. **b**, NHMUK PV P 56495 in medial view. **c**, right mandible (mirrored) of NHMUK PV P 53249 in lateral view. **d**, NHMUK PV P 56495 in dorsal view. **e**, NHMUK PV P 56495 in ventral view. **f**, NHMUK PV P 56495 articular region in dorsal view. **g**, NHMUK PV P 56495 in posterior view. **h**, NHMUK PV P 56495 anterior region in dorsal view. Scale bar = 2 mm. Panels f, h not to scale. *Abbreviations*: add.fo, adductor fossa; ang, angular; art, articular; co.do, dorsal part of coronoid; co.pa.sh, coronoid-prearticular shelf; co.pa, coronoid-prearticular plate; co.te, coronoid teeth; co.ve, ventral part of coronoid; cot, cotyle; de, dentary; de.sh, dentary shelf; de.te, dentary teeth; den, denticles; e.mand.VII, external mandibular branch of the facial nerve; f.mand.V, mandibular branch of the trigeminal nerve; for, foramen; i.mand.VII, internal mandibular branch of the facial nerve; id.gr, interdental groove: max, area overlapped by the maxilla; ma.te, marginal teeth; mc, mandibular canal; mc.po, pores connecting to the mandibular canal; mec, Meckel’s cartilage; pa, prearticular; p.l, pit line; r.soc, replacement socket.

The dorsal margin of the lateral surface of the jaw is straight along most of its length. It turns upwards slightly at the anterior end. Near the posterior end of the jaw, above the adductor fossa, the dorsal margin is sinusoidal, so it forms a low crest. The ventral margin is convex in the posterior half of the jaw. In the anterior half, the ventral margin is straight and slopes upwards, so that the jaw narrows towards the anterior end. Close to the anterior end, the ventral margin turns upwards more sharply, resulting in the whole jaw being angled slightly dorsally, and tapering towards the tip. Viewed dorsally, the jaw is straight in its posterior two-thirds, and gently curves medially in the anterior third. In the posterior portion of the jaw, on the lateral surface, a shallow, crescentic depression lateral to the adductor fossa (max, Fig. 7a) accommodated the ventral extension of the maxilla. This area lacks ornamentation.

The lateral surface of the jaw is strongly ornamented, such that it largely obscures the suture between the dentary and infradentary. The ornament is deep and consists of somewhat irregularly arranged grooves and ridges that are generally oriented from anterior to posterior but break into broad tubercles more dorsally. Near the dorsal margin, these tubercles become more pointed and grade imperceptibly into the marginal teeth. Most of the lateral surface of the jaw is occupied by the dentary (de, Fig. 7a,e), which has a slight dorsal rim at its lateral extent but otherwise has a flat dorsal margin that extends into a stubby medial shelf (de.sh, Fig. 7d,h). This shelf overlies both the Meckel’s cartilage and a splint-like coronoid-prearticular shelf (co.pa.sh, Fig. 7d), but is separated from the tooth-bearing regions of the dermal ossifications by a substantial interdental groove (id.gr, Fig. 7d) along its entire length. A narrow angular (ang, Fig. 7a,e) represents the only infradentary. It occupies the posteroventral margin of the jaw and is largely overlapped by the dentary. Dorsally, the angular contributes to the posterior half of the maxillary overlap area. More ventrally, the angular is much narrower, but extends a little way anterior to the adductor fossa.

The main dentary tooth row preserves 17 teeth (de.te, Fig. 7a,d,h). They are short, squat and conical in shape, almost vertically oriented but with a slight medial curve. The anteriormost teeth are the smallest in the row, with the largest teeth found in the middle region of the jaw, generally decreasing in size again posteriorly. Generally, most teeth alternate with replacement sockets (r.soc, Fig. 7d,h). However, there are two pairs of adjacent teeth, and three sections of four or five adjacent replacement tooth sockets. An additional field of small marginal teeth (ma.te, Fig. 7a,h), loosely organized into two rows, is present lateral to the main dentary tooth row. It comprises three types of teeth: sharp, pointed teeth that are approximately half the size of the main dentary teeth; smaller, slightly blunter teeth interspersed between the sharper teeth, generally in pairs; and a lateralmost irregular row of low, rounded, laterally directed cusps that interdigitate with the dermal ornament tubercles. Replacement sockets are rare between the marginal teeth, and not observed between the lateralmost cusps.

The medial, dermal tooth-bearing bones ossified as a single element encompassing the coronoids and prearticular (co.pa, Fig. 7b,d). This coronoid-prearticular plate has two components: a medial surface, which bears teeth or denticles and extends further down the medial face of the mandible posteriorly; and a lateral shelf (co.pa.sh, Fig. 7d) extending between the dentary shelf and Meckel’s cartilage, which thins to a splint anteriorly but becomes more laterally extensive posteriorly to form the anterior margin of the adductor fossa (add.fo, Fig. 7d,f). The medial tooth-bearing surface is dorsally extensive, reaching to slightly below the tips of the main dentary teeth. The anterior end of the coronoid-prearticular plate broadens medially and flattens, ending in a rectangular, anteromedially facing surface. Close to the medial margin of this surface, a narrow, elongate opening (for, Fig. 7d,h) extends all the way through the plate. At least one canal opens horizontally to the lateral surface of the coronoid. A dorsal and ventral component to the anterior coronoid is just perceptible, and substantially less developed than in *Gogosardina coatesi* (Fig. 1i) or previously described mandibles of *M. toombsi* (Gardiner, 1984). Although shallow anteriorly, the coronoid-prearticular plate increases in depth posteriorly, reaching its maximum level with the midpoint of the adductor fossa. Its posterior margin is rounded, and its dorsal margin projects slightly above the dorsal margin of the adductor fossa. A narrow canal for the internal mandibular branch of the facial nerve (imand.VII, Fig. 7b) notches the posterodorsal corner of the coronoid-prearticular plate, piercing the mandible and running between the plate and Meckel’s cartilage.

Almost the entire medial dermal series is covered in a shagreen of randomly arranged, blunt denticles (den, Fig. 7b,h). Most of the denticles are similar in size, though a few randomly dispersed cusps are larger, and there is a general decrease in size close to the ventral and posterior margins of the plate. A single row of slightly larger, very blunt denticles runs along the dorsal margin of the coronoid-prearticular series. A few larger, sharper teeth are present around the margins of the elongate opening (for, Fig. 7d,h) through the anteriormost coronoid; these teeth are smaller than the main dentary tooth row but notably larger than the denticles, and they curve towards the coronoid cavity.

The path of the mandibular canal in NHMUK PV P 56495 is barely traceable in lateral view, marked only by a few widely spaced pores (mc.po, Fig. 7c). These are positioned near the posterior and ventral margin of the mandible in its posterior half, but arch slightly anteriorly in the anterior half. Approximately one third of the way along the jaw, the canal travels vertically and terminates in a curved pit line (p.l, Fig. 7a); there is no trace of the canal within the mandible more anteriorly. A few sporadically placed pores also open onto the medial surface of the dentary.

The path of the mandibular canal differs somewhat in NHMUK PV P 53249 (Fig. 7c), although we note that the lower quality of the scan gives the erroneous impression that the canal is open in a groove in the anterior half of the mandible. In this specimen, the mandibular canal extends from the posterodorsal margin of the mandible almost to its extreme anterior end. The path can be traced by a combination of pores and grooves on the lateral surface of the jaw, with a small number of pores opening onto the medial surface. As in NHMUK PV P 56495, the canal initially closely follows the posterior and ventral margin of the mandible in a thickened portion of the angular and, subsequently, the dentary. Approximately one-third from the anterior end of the jaw, the canal deviates from the ventral jaw margin and gently rises to the mid-height of the jaw. Unlike in NHMUK PV P56495, the canal continues, exiting the anterior margin of the dentary via two foramina level with the termination of Meckel’s cartilage.

Meckel’s cartilage (mec, Fig. 7b,g) is ossified as a single element along the entire length of the jaw, terminating in an unfinished cup with limited endochondral ossification at the anterior end of the jaw. In the anterior half of the jaw, Meckel’s is ossified as a shallow perichondral shell restricted to the dorsal half of the mandible. From roughly the midpoint of the jaw to the articular region, it extends from beneath the coronoid-prearticular shelf to almost the ventral margin of the jaw. There is little in the way of endochondrally ossified bone in this region. A large, ovoid foramen for the mandibular branch of the trigeminal nerve (f.mand.V, Fig. 7b) is present at the ventral margin of Meckel’s cartilage where it articulates with the dentary, immediately posterior to the deepest point of the jaw. The posteroventral margin of Meckel’s cartilage bears a deep, wide groove (emand.VII, Fig. 7e,g), which continues forward a short distance to beneath the deepest point of the jaw.

The articular region (art, Fig. 7b,d,f) consists of more densely ossified endochondral bone, extending from the anterior end of the medial margin of the adductor fossa to approximately halfway along the lateral margin of the adductor fossa. The articular is mostly coated in perichondral bone, but the bone surface is unfinished in some patches within the adductor fossa, and inside the medial articular cotyle. The adductor fossa (add.fo, Fig. 7d,f) comprises just under 20% of the length of the jaw. It is widest posteriorly, and tapers quite strongly anterior, resulting in a triangular shape in dorsal view Two articular cotyles (cot, Fig. 7d,f) are present on the articular: both are close to circular in shape, deep, and face directly dorsally. They are almost laterally aligned, although the more medial coronoid is positioned slightly more anteriorly.

#### 3.4.2. Mimipiscis bartrami

The lower jaw of *Mimipiscis bartrami* (Fig. 8) has previously been described by Gardiner (1984) as *M. toombsi* based on external morphology of acid prepared specimens, prior to Choo (2012) recognizing two distinct species within the genus. Our description is based on NHMUK PV P 53254, a near-complete acid-prepared mandible missing only the anterior tip (preserved length 25 mm). As with NHMUK PV P 53249 (Fig. 7c), the low quality of the tomography data does not allow for detailed observation of minute features, so we express caution when interpreting these.

**Fig. 8:**
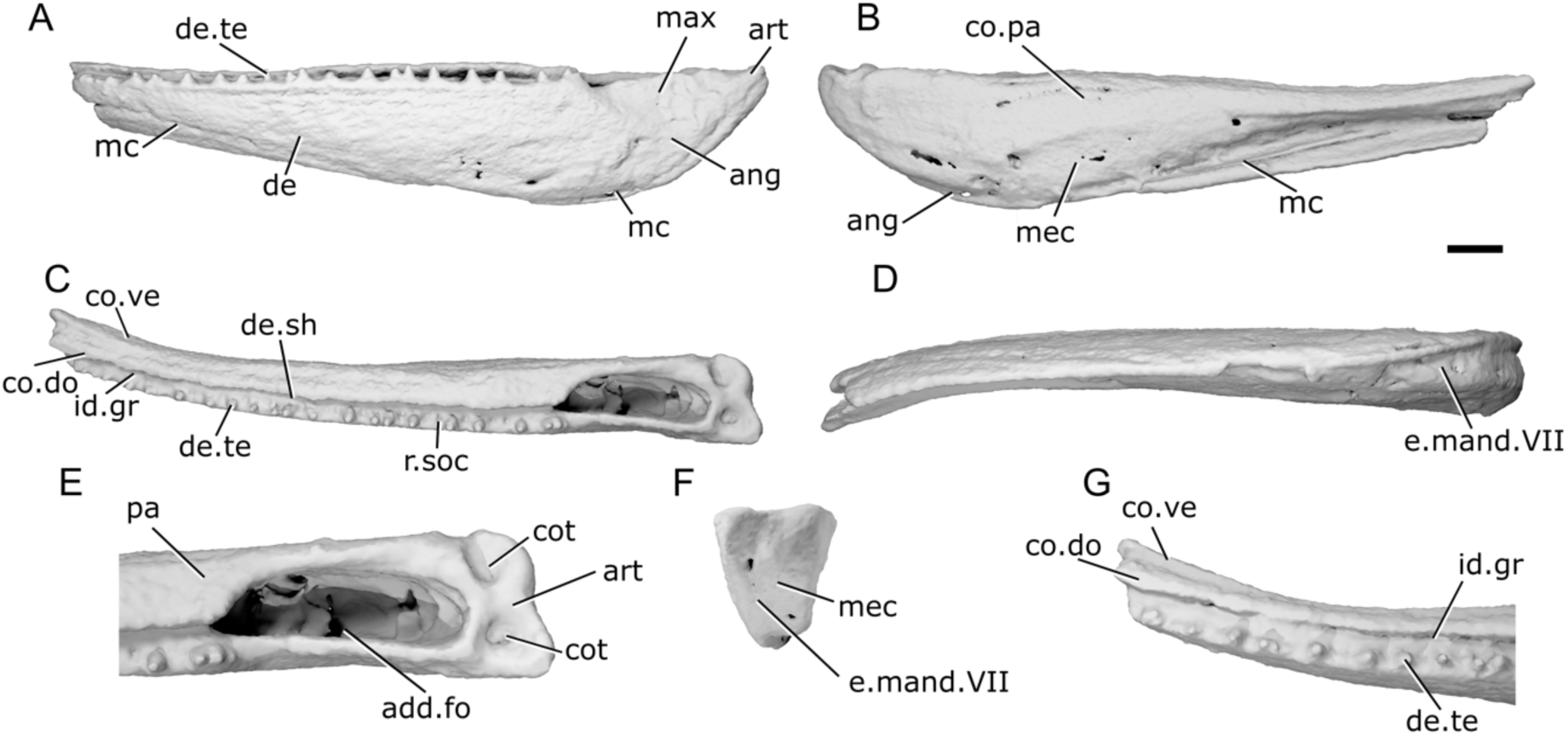
Mandible of *Mimipiscis bartrami* NHMUK PV P 53254. **a**, Right mandible (mirrored) in lateral view. **b**, medial view. **c**, dorsal view. **d**, ventral view. **e**, articular region in dorsal view. **f**, posterior view. **g**, anterior region in dorsal view. Scale bar = 2 mm. Panels e, g, not to scale. *Abbreviations*: add.fo, adductor fossa; ang, angular; art, articular; co.do, dorsal part of the coronoid; co.pa, coronoid-prearticular; co.ve, ventral part of the coronoid; cot, cotyle; de, dentary; de.sh, dentary shelf; de.te, dentary teeth; e.mand.VII, external mandibular branch of the facial nerve; id.gr, interdental groove; max, area overlapped by the maxilla; mc, mandibular canal; mec, Meckel’s cartilage; pa, prearticular; r.soc, replacement socket.

The dorsal margin of the lower jaw of *M. bartrami* is almost completely straight. The ventral margin bulges below the position of the adductor fossa, then slopes upwards steadily towards the anterior end of the jaw. In dorsal view, the jaw is mostly straight, and curves very gently medially in its anterior third. The adductor fossa has a straight lateral margin in dorsal view, and the medial margin is straight along most of its length, giving it an ovoid outline.

The dentary (de, Fig. 8a) comprises most of the lateral dermal surface of the lower jaw in *M. bartrami*. In axial section, the dentary bears a stubby medial shelf (de.sh, Fig. 8c) that lies over both Meckel’s cartilage and a corresponding lateral shelf of the coronoid-prearticular plate. This results in an appreciable interdental groove (id.gr, Fig. 8c,g) between the dentary and medial dermal tooth rows. The angular (ang, Fig. 8a,b) is a small, narrow, crescentic element restricted to the very posteroventral margin of the jaw. There is no evidence of a separate surangular. Ornament is difficult to describe in detail given the poor quality of the scan, but appears to consist of irregular ridges angled roughly antero-posteriorly. This is corroborated by external photographs of the mandible of *M. bartrami* that show strong ornament of elongate ridges, shorter ridges and tubercles more dorsally (Choo, 2012: figs. 5a,6a,7a). The postderodorsal corner of the lateral surface of the jaw also has a concave depression in the region that would have been overlapped by the maxilla (max, Fig. 8a).

A row of teeth (de.te, Fig. 8a,c,g) extends along the full length of the dentary. The teeth are short, sharp and robust with a wide base. They are oriented vertically and not recurved. The teeth are relatively densely spaced with a small number of replacement sockets (r.soc, Fig. 8c); many teeth are adjacent to one another, and there are more teeth than replacement sockets. At least one row of smaller teeth lies lateral to the main dentary tooth row, and comparison with previously described material suggests that these interdigitate with denticle-like dermal ornament (?den; Choo, 2012: fig. 7)

The medial dermal bones (co.pa, Fig. 8b) are ossified as a single prearticular-coronoid series without visible sutures, although this may be an artefact of scan quality. They comprise approximately a third of the depth of the jaw in the anterior half and increase to cover a little over half of the depth in the posterior region. The coronoid-prearticular consists of a relatively flat vertical medial surface and a horizontal lateral shelf that contact another at a sharp corner. The medial shelf reaches dorsally almost as far as the tips of the dentary teeth, but does not extend above them. Anteriorly, the coronoid-prearticular plate is separated into dorsal (co.do, Fig. 8c,g) and ventral (co.ve, Fig. 8c,g) components by a shallow groove, which tapers posteriorly and disappears shortly behind the midlength of the coronoid region. Teeth on the medial dermal bones are difficult to make out in tomograms due to the poor quality of the scan, but larger, low-crowned teeth appear to be present on more dorsal regions of the coronoid-prearticular plate, with minute denticles on the more ventral portions.

Meckel’s cartilage (mec, Fig. 8b,f) is ossified throughout the entire length of the jaw. In the anterior region, it is restricted to a dorsal position in between the dentary and medial dermal bones. In the posterior half of the jaw, Meckel’s cartilage extends ventrally below the level of the medial dermal bones, nearly reaching the ventral margin of the jaw. At the posterior end of the jaw, Meckel’s cartilage forms the articular region (art, Fig. 8a,e). Two small, round articular cotyles (cot, Fig. 8e) are present. They both face primarily dorsally, though the more medial cotyle is slightly further forward and faces slightly dorsomedially.

The path of the mandibular canal (mc, Fig. 8e,b) can be traced by a small number of pores on the lateral surface of the mandible. One cluster of pores can be observed very close to the posterior end of the jaw, close to the ventral margin. A further series of roughly six pores indicate that the canal sloped anteriorly, then plateaued close to the anterior end of the mandible. As with NHMUK PV P 53249, we note that the apparent groove in the anterior portion of the mandible is an artefact of low scan quality; the canal was enclosed laterally.

#### 3.4.3. Moythomasia durgaringa

Description of the mandible of *Moythomasia durgaringa* (Fig. 9) is based on X-ray CT of AMNH FF 11598, a near-complete but somewhat disarticulated complete fish (mandible length: 30 mm). The jaw of this taxon has previously been described by Gardiner (1984) based on acid-prepared material, with subsequent treatment by Choo (2015).

**Fig. 9:**
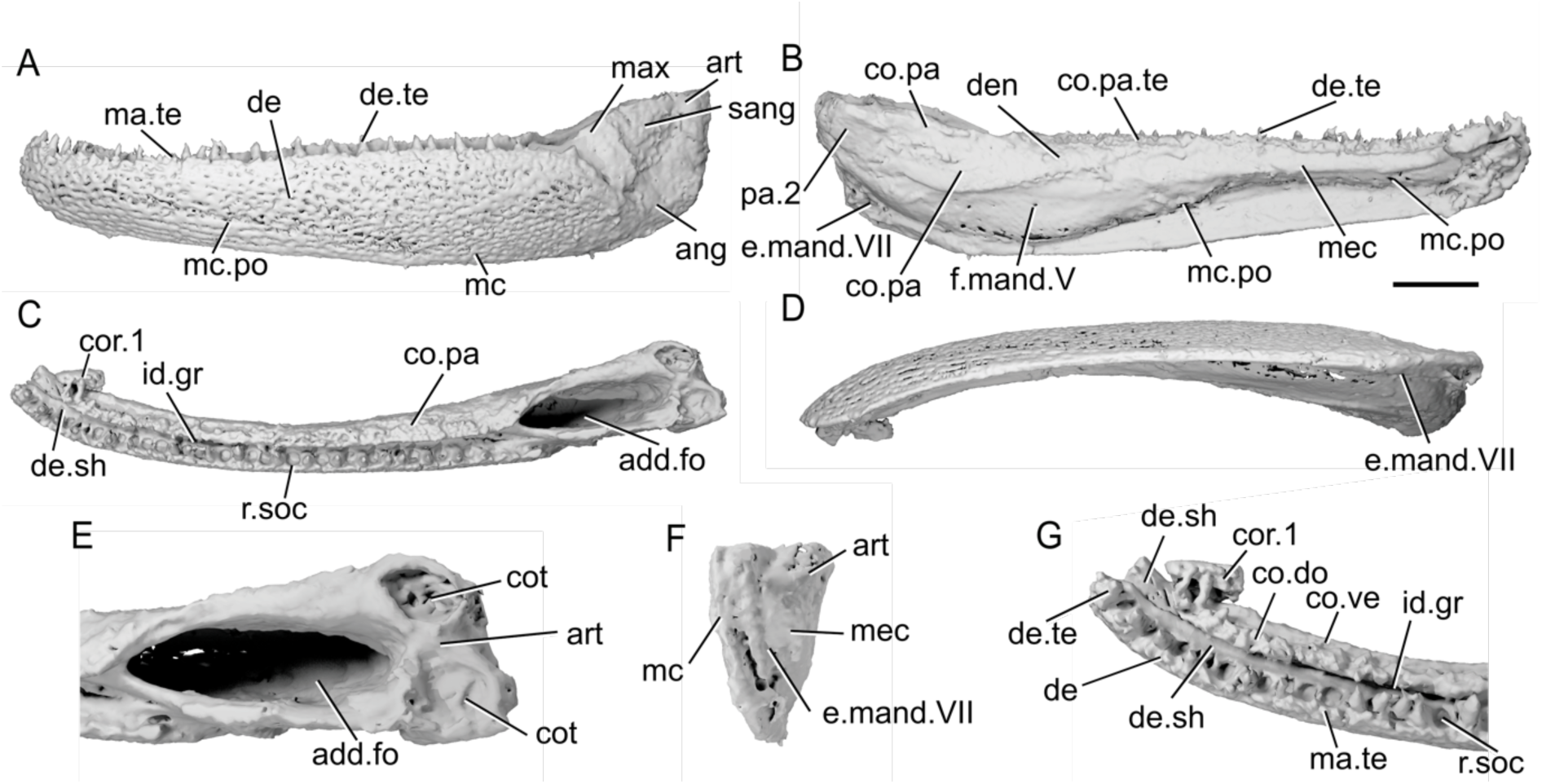
Mandible of *Moythomasia durgaringa* AMNH FF 11598. **a**, Right mandible (mirrored) of AMNH FF 11598 in lateral view. **b**, medial view. **c**, dorsal view. **d**, ventral view. **e**, articular region in dorsal view. **f**, posterior view. **g**, anterior region in dorsal view. Scale bar = 5 mm. Panels e, g not to scale. *Abbreviations*: add.fo, adductor fossa; ang, angular; art, articular; co.do, dorsal part of the coronoid; co.pa, coronoid-prearticular; co.pa.te, coronoid-prearticular teeth; co.ve, ventral part of the coronoid; cor, coronoid; cot, cotyle; de, dentary; de.sh, dentary shelf; de.te, dentary teeth; den, denticles; e.mand.VII, external mandibular branch of the facial nerve; f.mand.VII, mandibular branch of the trigeminal nerve; id.gr, interdental groove; ma.te, marginal teeth; max, area overlapped by the maxilla: mc, mandibular canal; mc.po, pores connecting to the mandibular canal; mec, Meckel’s cartilage; r.soc, replacement socket; pa.2, posterior prearticular; sang, surangular.

The overall profile of the mandible in *Moythomasia durgaringa* is straight, with a blunt anterior margin and raised articular region. The dorsal margin of the dentary is slightly concave in its anterior half. This results in a morphology in which the jaw bulges at the anterior tip, but only marginally. At the posterior margin of the mandible, both the dorsal and ventral margins turn upwards. The mandible is concave along almost its entire length in dorsal view, though is straight for a short portion at the midpoint of the jaw. A large ovoid depression on the lateral surface of the mandible, close to its posterodorsal end, represents the maxilla overlap area (max, Fig. 9a). This encompasses the surangular, much of the dorsal half of the angular, and a short section of the dentary.

The dentary (de, Fig. 9a) is large, making up the entire height of the jaw for approximately 85% of mandible length. A short horizontal shelf (de.sh, Fig. 9c,g) projects from the medial surface of the dentary near its dorsal margin. This shelf abuts, but does not substantially under-or overlie, the dermal medial tooth-bearing bones, except immediately anterior to the adductor fossa. As a result, there is a wide interdental groove (id.gr, Fig. 9c,g) between the teeth of the inner and outer tooth-bearing bones. The angular (ang, Fig. 9a) is also large, with a robust anterior ramus that projects to just beyond the anterior end of the adductor fossa and makes up more than half of the jaw height beneath the adductor fossa. The angular also possesses a robust dorsal ramus that reached to the posterior third of the adductor fossa and comprises the posterior and dorsal portions of the lateral mandibular surface. A small surangular (sang, Fig. 9a) is confined entirely within the maxillary overlap area. It is roughly triangular, with a straight anterior margin and a gently curved dorsal margin that increases in height posteriorly.

Almost the entire lateral surface of the dentary is ornamented, excepting only the posterior region. The ornamentation consists of deep pits and pores that mostly connect to small vacuities below the surface of the dentary that are not linked to the main path of the mandibular canal. In the anteroventral corner, the pits are aligned into approximately four parallel horizontal lines. These lines converge posteriorly to a point approximately midway along the dentary, and the ornament comprises irregular pores dorsal and posterior to this region. The angular appears unornamented, other than a slightly rugose surface at the posterior end.

Twenty teeth (de.te, Fig. 9a,b,g) are present on the dentary, their bases originating slightly ventral to its dorsal margin. Each tooth is fairly robust, conical, and oriented vertically. The teeth become slightly smaller anteriorly, but the posteriormost tooth is also small. Teeth mostly alternate with replacement tooth sockets, though a few pairs of adjacent teeth or sockets are present, as well as one series of four teeth in a row. A single row of much smaller, blunt teeth (ma.te, Fig. 9a,g) lie lateral to the main tooth row and grade into the dermal ornament.

The medial dermal tooth-bearing bones are ossified as a single coronoid-prearticular series (co.pa, Fig. 9b,c). The anterior tip of the series, corresponding with the anterior part of the first coronoid (cor.1, Fig. 9c,g), has broken off and rotated medially out of life position. The prearticular region of the series extends posteriorly as far as the articular cotyle, and lines the medial margin of the adductor fossa. It is shallow, extending less than half the way down the medial surface of the mandible. A small, oblong, separately ossified prearticular (pa.2, Fig. 9b) lies medial and ventral to the articular cotyle, as previously described in this taxon by Gardiner (1984: fig. 95).

Dorsally, the prearticular-coronoid plate is raised into a distinct ridge that projects almost as high as the tips of the dentary teeth. The ridge bears several irregular rows of low, blunt teeth (co.pa.te, Fig. 9b), all smaller than those on the dentary. In the anterior third of the mandible these teeth are bounded ventrally by a shallow groove that separated the dorsal (co.do, Fig. 9g) and ventral (co.ve, Fig. 9g) components of the coronoids. Dorsal to this ridge, the anteriormost (displaced) coronoid bears two enlarged teeth. Smaller, more rounded denticles (den, Fig. 9b) are present on the more ventral portions of the entire plate. In the anterior half of the mandible, a short lateral coronoid shelf abuts the horizontal shelf of the dentary. More posteriorly, in the prearticular region anterior to the adductor fossa, the shelf becomes more laterally extensive and underlies that of the dentary.

A mandibular canal (mc, Fig. 9a,f) is present along the whole length of the jaw. Posteriorly, it begins at the posterodorsal corner of the mandible, entering the angular immediately behind the lateral condyle and travelling in a narrow canal along the very margin of the angular. It passes into the dentary level with the anterior margin of the adductor fossa, along the ventral margin. More anteriorly, it gently curves upwards before arching sharply dorsally and then, in the anterior third of the jaw, ventrally. It terminates through an opening at the anteroventral tip of the jaw. There is frequent communication between the mandibular canal and the lateral surface of the dentary via a high density of surface pits and pores (mc.po, Fig. 9a,b). However, no pits are present in the posterior quarter of the mandible. Just four widely spaced pores connect the canal to the medial surface of the dentary.

Meckel’s cartilage (mec, Fig. 9b,f) is ossified as a single element along the entire length of the jaw. Endochondral ossification is present in all regions within the perichondral lining, although is best developed in the anterior and posterior regions. Anteriorly, Meckel’s is positioned between the coronoid and the dentary and does not expand ventral to the level of the coronoid series. It opens as unfinished bone at the anterior tip of the jaw, though there is perichondral bone present in this region ventrally, in the gap between the coronoid and dentary. In the articular region (art, Fig. 9a,e,f), Meckel’s cartilage is well ossified and rises dorsal to the dentary tooth row. The adductor fossa (add.fo, Fig. 9c,e) is an elongate ovoid in shape and narrows slightly anteriorly. Only a few small patches within the adductor fossa lack perichondral coating. The two large articular cotyles (cot, Fig. 9e) are almost circular, but slightly anteroposteriorly elongate. Anterior to the adductor fossa, Meckel’s cartilage extends well ventral to the prearticular, almost reaching the lower margin of the mandible, before becoming more dorsally restricted. In this region, Meckel’s cartilage is predominantly just a perichondral shell, with only sporadic patches of endochondral bone. Immediately anterior to the level of the adductor fossa, the medial portion of this shell is pierced by two foramina for the mandibular branch of the trigeminal nerve (f.mand.V, Fig. 9f). A groove, housing the external mandibular branch of the facial nerve (e.mand.VII, Fig. 9b,f), runs along the lateral junction of Meckel’s with the dentary; it extends from the posterior tip of the jaw to just posterior to the anterior tip of the adductor fossa. Within this groove, below the adductor cotyles, are several foramina for the internal mandibular branch of the facial nerve.

#### 3.4.4. Raynerius splendens

Both lower jaws are present in MGL 1245, the holotype and only known specimen of *Raynerius splendens* (Fig. 10), and were originally described via X-ray CT by Giles et al. (2015b). We add to this description by reexamining the existing tomograms. The left lower jaw is less damaged overall, but its anterior portion is missing. The right lower jaw shows substantial damage along most of its lateral margin and articular region but is complete anteriorly (total length: 22 mm).

**Fig. 10:**
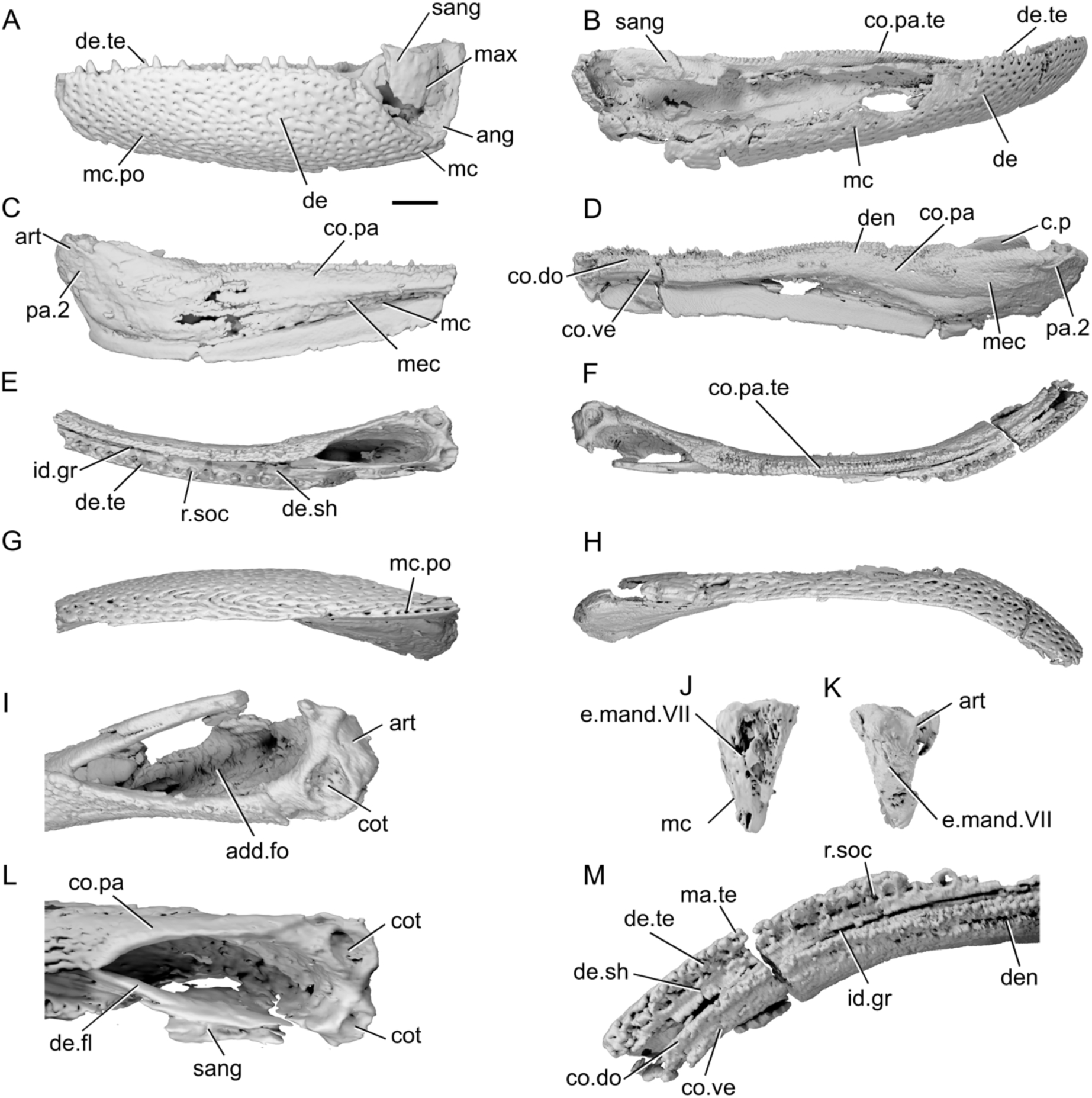
Mandibles of *Raynerius splendens* MGL. 1245**. a**, Left mandible in lateral view. **b**, right mandible in lateral view. **c**, left mandible in medial view. **d**, right mandible in medial view. **e**, left mandible in dorsal view. **f**, right mandible in dorsal view. **g**, left mandible in ventral view. **h**, right mandible in ventral view. **i**, articular region of right mandible in dorsal view. **j**, left mandible in posterior view. **k**, right mandible in posterior view. **l**, articular region of left mandible in dorsal view. **m**, anterior region of right mandible in dorsal view. Scale bar = 2 mm. Panels i, l, m not to scale. *Abbreviations*: add.fo, adductor fossa; ang, angular; art, articular; c.p, coronoid process; co.do, dorsal part of the coronoid; co.pa, coronoid-prearticular; co.pa.te, coronoid-prearticular teeth; co.ve, ventral part of the coronoid; cot, cotyle; de, dentary; de.fl, dentary flange; de.sh, dentary shelf; de.te, dentary teeth; den, denticles; e.mand.VII, external mandibular branch of the facial nerve; id.gr, interdental groove; ma.te, marginal teeth; max, area overlapped by the maxilla; mc, mandibular canal; mc.po, pores connecting to the mandibular canal; mec, Meckel’s cartilage; pa.2, posterior prearticular; r.soc, replacement socket; sang, surangular.

The ventral margin of the mandible is almost completely straight but inclined anterodorsally. Close to the anterior end, it turns upwards strongly. The dorsal margin of the mandible is also largely straight, although the articular region is raised above the toothrow. Overall, the jaw is stout and does not taper significantly at the anterior end, as noted by Giles et al. (2015b). In dorsal view the jaw is concave, most strongly so at the anterior end of the jaw. Close to the posterodorsal margin of the mandible, a shallow embayment corresponds with the overlap of the maxilla (max, Fig. 10a). This area is triangular, with the apex pointing ventrally; it has a nearly vertical posterior margin and a sloping anterior margin. The surangular is entirely contained within this region, but bone preservation is poor.

The dentary (de, Fig. 10a,b) is the largest dermal element on the lateral surface of the jaw. It covers much of the lateral jaw surface as far as the area of the maxillary overlap. In the dorsal part of this area, a short, unornamented flange (de.fl, Fig. 10l) projects from the dentary to form the anterior and lateral margin of the adductor fossa and underlies the surangular. Ventral to this flange, the posteroventral margin bounds the ventral half of the maxillary overlap area. This sloping posterior margin of the dentary terminates a point at the posteroventral corner of the jaw. A broad medial shelf (de.sh, Fig. 10e) is present along the entire length of the dentary, reaching to the inner surface of the coronoid-prearticular plate and resulting in a substantial interdental groove (id.gr, Fig. 10e,m) between the inner and outer tooth rows.

Two infradentaries are present, though both are incompletely preserved. The angular (ang, Fig. 10a) is evident in the left jaw. It is narrow and rod-like, restricted to the ventral and posterior margin of the jaw, and predominantly formed as a tube surrounding the mandibular canal. It is underlapped by the dentary and extends anteriorly to approximately the anterior margin of the adductor fossa. A small, rhomboidal surangular (sang, Fig. 10a,b) is present in the area overlapped by the maxilla. It overlies the posterodorsal flange of the dentary, but incomplete preservation means its contact with the angular is unclear. The dorsal margin of the surangular appears to extend some way above the adductor fossa, forming a rudimentary coronoid process (c.p, Fig. 10d).

The damage to both jaws prevents a full count of the tooth number. However, the number of functional teeth is quite low, with most of the single row of teeth on the dentary (de.te, Fig. 10a,b,e,m) separated by a pair of replacement sockets, some teeth by one socket, and one pair of adjacent teeth. The teeth are large, robust, vertical cones and acrodin caps are present on the largest of them. As the teeth leave from the medial shelf of the dentary, a substantial portion of each is obscured in lateral view. A single row of small, blunt teeth (ma.te, Fig. 10m) sits atop the lateralmost edge of the dentary.

The dentary ornament is smooth and punctured by numerous large, regularly spaced pores. These pits are mostly circular, but some are elongate, and drawn out into short grooves. Generally, they are scattered across the lateral jaw surface with no particular pattern. However, the pores that mark the path of the mandibular canal (mc.po, Fig. 10a) tend to be more densely spaced and many are chevron shaped.

The path of the mandibular canal (mc, Fig. 10a,b,c) is completely preserved until the posterodorsal corner of the jaw, running close to the posterior and then ventral margin. Around a third of the way from the posterior margin of the mandible, the canal angles anterodorsally and is carried some way from the ventral jaw margin. It opens in a large pore at the anterior margin of the jaw, just dorsal to the midpoint.

The medial tooth-bearing dermal bones are continuously ossified (co.pa, Fig. 10c,d,l), and so the number of coronoids cannot be discerned. The series extends from the very anterior tip of the jaw to the anterior margin of the articular cotyles, and has a rounded posterior margin. It is restricted to the dorsal half of the jaw along its entire length. The plate consists of a large, medially facing, gently convex lamina on which teeth and denticles are borne. A short, thin lateral shelf extends some way beneath the medial shelf of the dentary. Similarly to *M. durgaringa* (Fig. 9b), a small, ovoid posterior prearticular (pa.2, Fig. 10c,d) lies posterodorsal to the main coronoid-prearticular plate; this was not identified by Giles et al. (2015b).

Very small, blunt teeth (co.pa.te, Fig. 10b,f) are present along the dorsal margin of the medial dermal bones. These are closely packed and form two to three rows. Medial to this toothrow is a narrow groove, below which lies is an additional, single row of blunt teeth. This groove tapers and terminates posteriorly, and the two toothrows merge behind the midpoint of the jaw. An anterior continuation of the groove divides the coronoids into dorsal (co.do, Fig. 10d,m) and ventral (co.ve, Fig. 10d,m) components. Much smaller denticles (den, Fig. 10d,m), which are at the limit of the scan resolution, are present on the ventral portions of the prearticular region.

Meckel’s cartilage (mec, Fig. 10c,d) is ossified as a single element that spans the length of the jaw. For most of its extent, it comprises a thin sheath of perichondral bone bridging the ventral gap between the dentary and the ventral margin of the medial dermal bones. It is restricted to the dorsal half of the mandible in the anterior half of the jaw but deepens posteriorly and extends ventral to the margin of the coronoid-prearticular plate. Endochondral ossification is more extensive at the anterior portion and in the articular region. The articular region (art, Fig. 10c,i,k) is somewhat elevated relative dorsal to the dentary tooth row, although to a lesser extent that in *Moythomasia durgaringa* (Fig. 9a). Two rounded articular cotyles are present and face dorsally and slightly medially. The adductor fossa (add.fo, Fig. 10i,l), of which the articular forms the posterior margin, is teardrop shaped; it is wide posteriorly, with a rounded posterior margin, and it tapers strongly anteriorly. A short, shallow, narrow groove is present at the very posterior tip of Meckel’s cartilage for the external mandibular branch of the facial nerve (e.mand.VII, Fig. 10j,k). Several additional small holes are present on the surface of Meckel’s cartilage close to the posterior end of the jaw, though as the surface is also incomplete in this area it is difficult to identify which may represent foramina.

#### 3.4.5. “*Moythomasia*” *devonica*

The lower jaw of “*Moythomasia*” *devonica* (Fig. 11) is described based on X-ray CT of BSNS E22113, a complete but isolated mandible (length: 30 mm) that does not appear to have undergone any significant distortion or compression. However, extensive pyrite intrusion through the centre of the specimen means that some details are obscured. The external surface of the mandible has previously been figured (under the original binomial *Rhadinichthys devonicus*: Hussakof & Bryant, 1918: pl. 62, 65) but has not been described in any detail. It was subsequently placed in *Moythomasia* by Gardiner (1963: 297), based on examination of sparse material at NHMUK and previous literature. However, the mandible differs conspicuously from that of *Moythomasia* and similar forms like *Raynerius*, calling into question this assignment. Consequently, we refer to this taxon as “*Moythomasia*” *devonica* here.

**Fig. 11:**
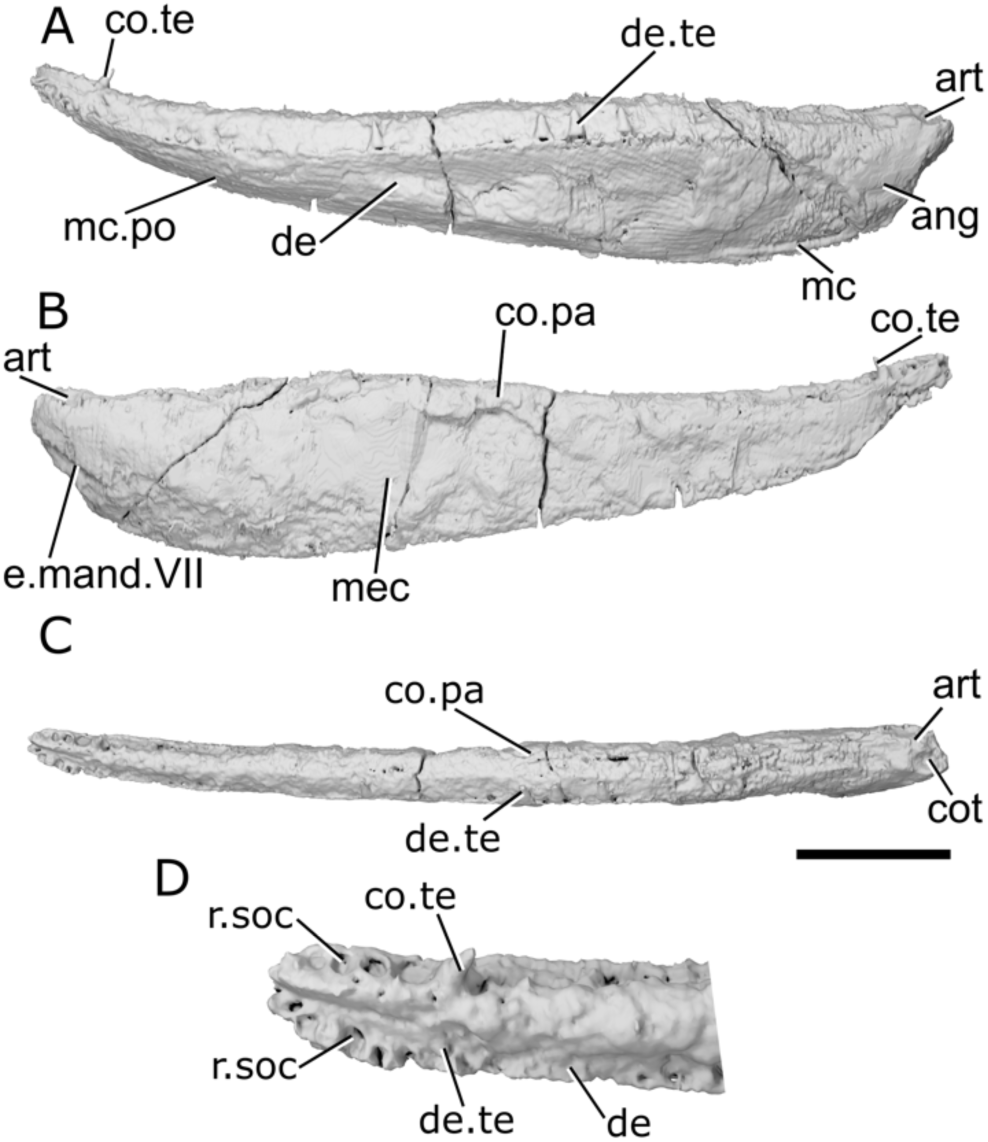
Mandible of *?Moythomasia devonica* BSNS E22113. **a**, Left mandible in lateral view. **b**, medial view. **c**, dorsal view. **d**, anterior region in dorsal view. Scale bar = 5 mm. Panel d not to scale. *Abbreviations*: ang, angular; art, articular; co.pa, coronoid-prearticular; co.te, coronoid teeth; cot, cotyle; de, dentary; de.te, dentary teeth; e.mand.VII, external, mandibular branch of the facial nerve; mc, mandibular canal; mc.po, pores connecting to the mandibular canal; mec, Meckel’s cartilage; r.soc, replacement socket.

The ventral margin of the jaw is convex overall, with a steeper anterodorsal orientation in the front third of the mandible and a with a sharp dorsal upturn at the posterior margin. The dorsal margin of the jaw is generally straight, but inclined upwards in its anterior quarter such that the jaw tapers in thickness anteriorly. As the dorsal articular region (art, Fig. 11a,b) extends some way posterior to the lateral dermal bones, the posterior margin of the mandible is sinusoid. There is no depression to indicate an area of overlap for the maxilla. In dorsal view the profile of the jaw is only very slightly concave, but it appears to have been somewhat flattened.

Almost the entire lateral surface of the jaw is made up of the dentary (de, Fig. 11a,d), which bears a sharp ridge along its upper lateral margin, just ventral to the tooth row. An angular (ang, Fig. 11a) is present, but is restricted to the very posterior end of the jaw; it is crescent shaped and very narrow. Ventrally, the margins of the angular are not preserved. There is no distinctly ossified surangular. Extensive pyritization within the jaw has obscured any details of the inner face of the dentary, such as the presence or extent of a medial dentary shelf. The external surface of the dentary is ornamented with fine, elongate, parallel ridges. The ridges are oriented horizontally in the ventral part of the jaw, but anterodorsally to posteroventrally in the dorsal part of the posterior two-thirds of the jaw.

Only part of the dentary dentition is preserved, with four teeth present within their corresponding sockets (de.te, Fig. 11a,c,d) and the impressions of several more visible. Each tooth is vertically oriented, conical, and robust. Marginal dentition appears to be absent, although this may be due to specimen preparation. A row of smaller replacement sockets (r.soc, Fig. 11d) in the anterior quarter of the dentary suggests that tooth size decreased anteriorly.

The medial dermal tooth-bearing bones are ossified as a single series (co.pa, Fig. 11b,c), and sutures between individual coronoids or the prearticular cannot be identified. Although it is not possible to tell the condition of any lateral coronoid-prearticular shelf, the coronoid-prearticular plate is in close contact with the dentary tooth row, with no interdental groove between the inner and outer dental arcade. The coronoid-prearticular plate is broad in dorsal view and very high, with its dorsal margin sitting some way above the tips of the dentary teeth. Very small, blunt, denticle-like tooth cusps, as well as more infrequent larger and pointed cusps, are distributed across the dorsal surface of the coronoid-prearticular plate. In the anterior fifth of the jaw, likely corresponding to the anteriormost coronoids, the dorsomedial margin of the coronoid develops a distinct ridge, along which sits a regular row of very small, conical teeth. A single larger, posteriorly curved coronoid tooth (co.te, Fig. 11a,b,d) the same size as the dentary teeth sits close to the anterior margin of the mandible. Four empty tooth sockets (r.soc, Fig. 11d) lie anterior to this tooth, becoming successively smaller in size towards the front of the jaw.

Meckel’s cartilage (mec, Fig. 11b) is ossified along the entire length of the jaw. Its depth along the mandible is difficult to determine due to pyrite growth. Similarly, the extent of endochondral and perichondral ossification cannot be determined with confidence, but at least the anterior portion is endochondrally ossified. The articular region (art, Fig. 11a,b,c) is fairly narrow and, as previously mentioned, extends further posteriorly than the lateral dermal bones. Two small, sub-circular cotyles (cot, Fig. 11c) are present, positioned close together. These are offset: the more lateral is posteriorly positioned and faces dorsally, and the more anterior is more anteriorly positioned and faces dorsomesially. A deep groove for the external mandibular branch of the facial nerve (e.mand.VII, Fig. 11b) is present on the posterior face of the articular.

In the posterior and anterior portions of the mandible, the mandibular canal (mc, Fig. 11a) has become infilled with pyrite, allowing its path to be traced with confidence. Small dorsal and ventral continuations of the pyrite indicate the presence of narrow tubules that would have connected to regular pores on the surface of the dermal jaw bones. Through the middle part of the dentary, the path can be traced by the continuation of these small pore openings (mc.po, Fig. 11a). The mandibular canal closely follows the ventral margin of the jaw. It is straight and rises gradually dorsally, away from the ventral margin, and opens via a rounded pore at the extreme anterior tip of the jaw.

#### 3.4.6. Osorioichthys marginis

*Osorioichthys marginis* is known from a single specimen, IRSNB P 1340, described by Taverne (1997), in which observations of the mandibles were restricted to the external surfaces. The specimen is dorsoventrally compressed and is sheared on one side, although this distortion affects only the left mandible and not the right. However, the density of the specimen, its large size (dentary: 48 mm), and the presence of highly attenuating mineral inclusions limit the level of detail that can be drawn from tomograms. In addition, the ventral margin of the dentary is exposed at the edge of the specimen and cannot be completely reconstructed (Fig. 12).

**Fig. 12:**
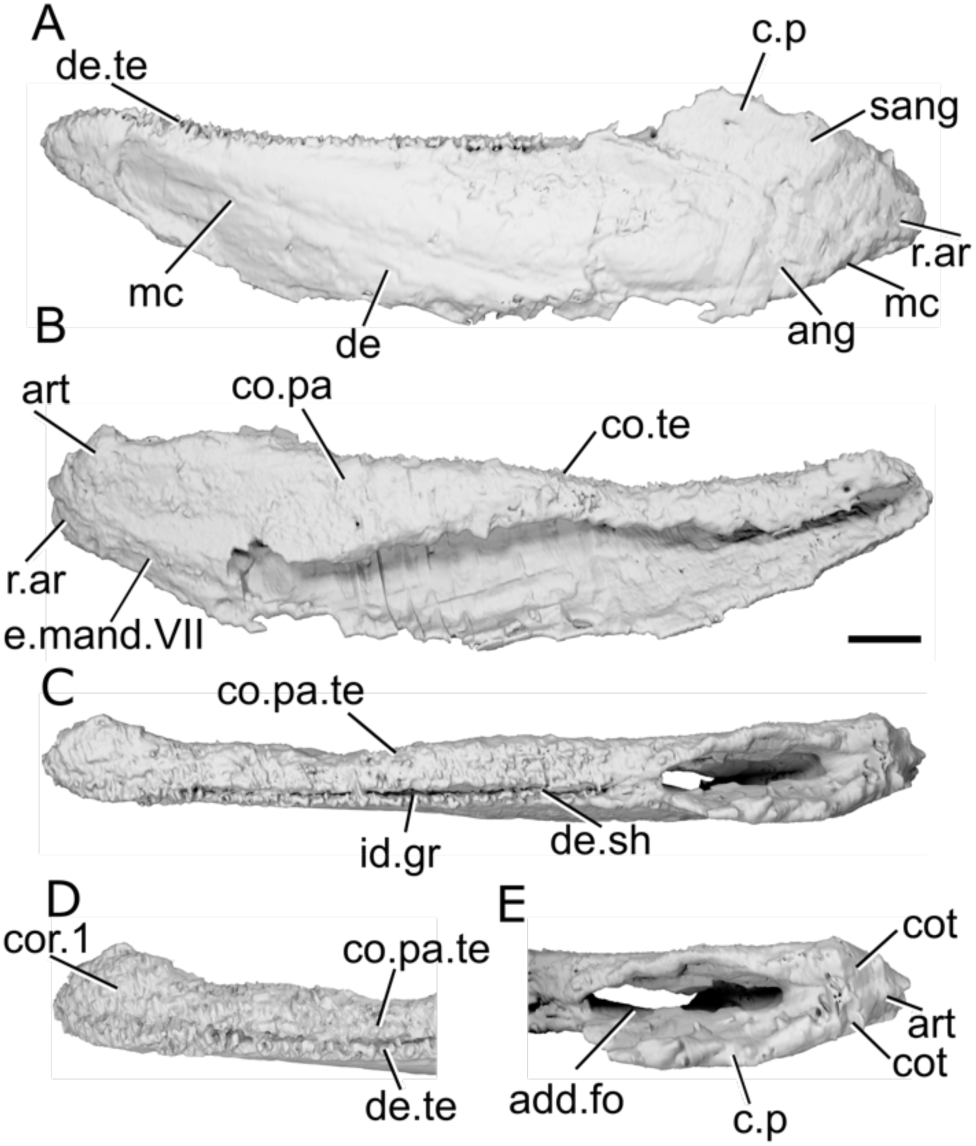
Mandible of *Osorioichthys marginis* IRSNB. **P** 1340**. a**, Right mandible (mirrored) in lateral view. **b**, medial view. **c**, dorsal view. **d**, anterior region in dorsal view. **e**, articular region in dorsal view. Scale bar = 5 mm. Panels d, e not to scale. *Abbreviations*: add.fo, adductor fossa; ang, angular; art, articular; c.p, coronoid process; co.pa, coronoid-prearticular; co.pa.te, coronoid-prearticular teeth; cor, coronoid; cot, cotyle; de, dentary; de.sh, dentary shelf; de.te, dentary teeth; e.mand.VII, external mandibular branch of the facial nerve; id.gr, interdental groove; mc, mandibular canal; r.ar, retroarticular process; sang, surangular.

In dorsal view the jaw appears mostly straight but broadens towards the midline at the symphysis. The ventral margin is convex along the whole length of the jaw. The dorsal margin is concave, though more gently, resulting in the jaw narrowing and turning slightly upwards towards the anterior end. Lateral to the adductor fossa, the dorsal margin of the jaw is raised into a rounded extension, and there is no indication of a depressed area for the ventral extension of the maxilla. The adductor fossa (add.fo, Fig. 12e) extends along approximately 20% of the length of the jaw and is straight sided with rounded anterior and posterior ends.

The lateral surface of the jaw comprises a dentary and two infradentaries. The dentary (de, Fig. 12a) makes up the entire depth of the jaw in the anterior two thirds, and its triangular posterior margin terminates in line with the midpoint of the adductor fossa. The angular (ang, Fig. 12a) is large and sub-rectangular, forming the posteroventral part of the jaw, and has a narrower anterior ramus. The surangular (sang, Fig. 12a) is small and projects dorsally above the rest of the jaw, forming a distinct and rounded coronoid process (c.p, Fig. 12a,e) that may also be contributed to by the dentary. The mode of preservation means that the ornament cannot be observed. Similarly, the path of the mandibular canal (mc, Fig. 12a) is difficult to trace, but appears to be marked by a slight ridge near the posteroventral corner of the jaw and a straight, shallow, anterodorsally oriented groove in the anterior half of the jaw.

Numerous small, sharp teeth (de.te, Fig. 12a,d) sit above the narrow dorsal surface of the dentary. The teeth are densely spaced, with few replacement sockets in between. A thin, short medial shelf (de.sh, Fig. 12c) extends towards, but does not reach, the medial tooth-bearing bones, resulting in a narrow interdental groove (id.gr, Fig. 12c) that tapers out anteriorly. A smaller row of lateral dentition appears to be absent on the dentary.

The medial dermal tooth-bearing bones appear to be ossified as a single coronoid-prearticular plate (co.pa, Fig. 12b). They form a slightly domed, toothed dorsal surface and a flat, medially facing shelf, which connect along a gently rounded margin. In the anterior half of the jaw the plate is very shallow, but deepens slightly posterior to the midpoint to extend a little over halfway down the medial surface of the jaw before shallowing again posteriorly. In dorsal view, the region corresponding to the anteriormost coronoid (cor.1, Fig. 12d) increases in width medially, and teeth become much smaller and more numerous.

The dorsal surface of the coronoid-prearticular plate and the anterior part of the medially facing shelf are covered with small, irregularly arranged teeth (co.pa.te, Fig. 12c,d). These teeth are only slightly smaller than the dentary teeth and their tips are at the same height, forming a continuous, broad field of dentition.

Meckel’s cartilage comprises separate articular and mentomeckalian ossifications. In the articular (art, Fig. 12b,e) region, it forms the posterior, lateral and medial walls of the adductor fossa (add.fo, Fig. 12e), with ossification ending just posterior of the anterior margin of the adductor fossa. Two cotyles (cot, Fig. 12e) are present behind the fossa: a slightly more anterior, dorsally facing cotyle, and a smaller cotyle that faces predominantly laterally. Both are extremely shallow. Unusually, Meckel’s cartilage extends behind the articular cotyles and lateral dermal ossifications, forming a sub-triangular retroarticular process (r.ar, Fig. 12a,b). A shallow groove along the ventral region of the articular accommodated the (emand.VII, Fig. 12b). The mentomeckelian is narrow and cyclindrical, and ossification is restricted to the anterior fifth of the jaw.

#### 3.4.7. Palaeoneiros clackorum

Parts of both jaws are present in the type and only specimen of *Palaeoneiros clackorum*, MCZ VPF-5114 (Fig. 13). The right mandible is the more complete of the two (preserved length: 22 mm), though neither retains the anterior portion of the jaw. The jaws were originally described by Giles et al. (2023) using X-ray CT, and here we supplement that description based on reexamination of the existing tomograms. In the preserved parts of the jaws, the ventral margin is nearly completely straight.

**Fig. 13:**
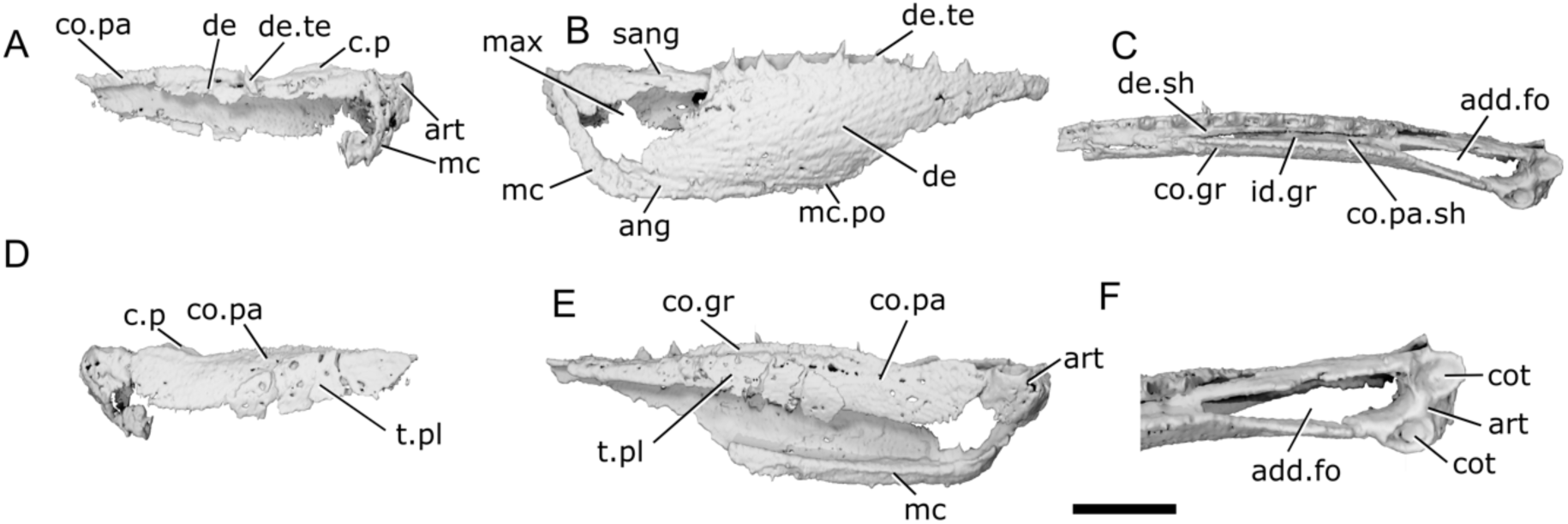
Mandibles of *Palaeoneiros clackorum* MCZ VPF-5114. **a**, Left mandible in lateral view. **b**, right mandible in lateral view. **c**, right mandible in dorsal view. **d**, left mandible in medial view. **e**, right mandible in medial view. **f**, articular region of right mandible in dorsal view. Scale bar = 5 mm. Panel f not to scale. *Abbreviations*: add.fo, adductor fossa; ang, angular; art, articular; c.p, coronoid process; co.gr, groove on the coronoid; co.pa, coronoid-prearticular; co.pa.sh, coronoid-prearticular shelf; cot, cotyle; de, dentary; de.sh, dentary shelf; de.te, dentary teeth; id.gr, interdental groove; max, area overlapped by the maxilla; mc, mandibular canal; mc.po, pores connecting to the mandibular canal; sang, surangular; t.pl, toothplate.

The dorsal margin is sinusoid. It is slightly convex in the region of the adductor fossa, forming a very low coronoid process. Anterior to this it forms a small trough, with the remaining margin, supporting the toothrow, gently convex. The adductor fossa is long and narrow, tapering anteriorly in dorsal view. A large, triangular area lateral and ventral to the adductor fossa indicates the position overlapped by the maxilla (max, Fig. 13b). In dorsal view the mandibles curve gently medially along their entire length. Much of the preserved lateral surface is made up of the dentary (de, Fig. 13a,b). Its ventral margin is thickened into a ridge for the mandibular canal on its medial surface.

Dorsally, beneath the toothrow, a medial shelf (de.sh, Fig. 13c) extends towards the medial tooth-bearing bones. This ridge is narrow posteriorly, barely projecting from the dentary, but becomes more pronounced anteriorly. It abuts the corresponding coronoid-prearticular shelf (co.pa.sh, Fig. 13c) along its entire length but is separated from the toothed portions by a substantial gap (id.gr, Fig. 13c). The angular (ang, Fig. 13b), forms the posterior jaw margin. It is narrow and rod-like, with a slim anterior ramus that underlies the dentary. Although the posterodorsal region of the lateral surface of the mandible is incomplete in both jaws, a shallow surangular (sang, Fig. 13b) forms a low, rounded coronoid process (c.p, Fig. 13a,d).

The lateral surface of the jaw is lightly ornamented with long, shallow, parallel grooves and ridges. In the dorsal half of the jaw the grooves slope anterodorsally to posteroventrally, and in the ventral half of the jaw they run parallel to the jaw ventral margin. The path of the mandibular canal is traceable by irregularly spaced pores (mc.po, Fig. 13b). It follows the ventral and posteroventral margin of the jaw very closely.

Eight well-preserved teeth (de.te, Fig. 13a,b) are present on the right jaw. An additional two partial teeth are preserved on both ends of this series. Each tooth is tall and narrow, and laterally flattened in cross section rather than conical. The teeth sit right on the dorsolateral margin of the dentary, with no lateral marginal dentition. The preserved dentary teeth almost entirely alternate with replacement tooth sockets, though the last two teeth are adjacent and there is a series of adjacent replacement sockets anteriorly.

The medial dermal tooth-bearing bones appear to be co-ossified as a single coronoid-prearticular series (co.pa, Fig. 13a,d,e). This series comprises a shallow vertical medial face that occupies less than half of the jaw depth along its entire length, and a lateral shelf (co.pa.sh, Fig. 13c). The lateral shelf is wide posteriorly and narrows anteriorly, mirroring the change in width of the corresponding medial shelf of the dentary. The dorsal margin of the medial face is raised into a continuous narrow crest that extends to slightly below the top of the dentary teeth. A closely packed series of small, sharp, medially directed teeth project from the dorsomedial surface of this crest. Immediately below this, a shallow groove (co.gr, Fig. 13c,e) extends approximately halfway along the anterior length of jaw. This is reminiscent of the groove in *Gogosardina, Mimipiscis, Moythomasia durgaringa* and *Raynerius* that separates the coronoid series into dorsal and ventral components, but in *Palaeoneiros* it terminates well short of the anterior margin of the mandible and there appear to be components of the coronoid series dorsal to its termination. Minute denticles are present immediately ventral to the groove. The more ventral components of the coronoid-prearticular shelf do not bear denticles; instead, their medial surface is covered by a series of three sub-ovoid toothplates (t.pl, Fig. 13d,e) that themselves bear denticles.

Meckel’s cartilage is unossified anterior to the articular region (art, Fig. 13a,e,f), and even here endo- and perichondral ossification is limited to the posterodorsal third of the adductor fossa. The two articular cotyles (cot, Fig. 13f) are close to circular in shape. Both articular cotyles face dorsally, although the medial cotyle is positioned anterior relative to the lateral cotyle.

#### 3.4.8. “Kentuckia” hlavini

The genus *Kentuckia* was established for *Kentuckia deani*, which is represented by three-dimensionally preserved material hosted in concretions of early Carboniferous age (Rayner, 1952). Older Devonian specimens have been assigned to this genus as “*Kentuckia*” *hlavini*, although this placement has been questioned (e.g., Friedman & Blom, 2006). The mandible of “*Kentuckia*” *hlavini* was originally briefly described by Dunkle (1964) based on external observations of the right mandible of an articulated specimen, in which the posterior margin is obscured by overlying dermal bones. Here we supplement this description through X-ray CT of CMNH imaging, an isolated left mandible (Fig. 14). This specimen, like the holotype, is strongly laterally compressed, although otherwise complete (24 mm long) and as a result the mandible appears extremely narrow and straight in dorsal view. Overall, the mandible is long and shallow, roughly one-fifth as deep as it is long. It is deepest close to the posterior end, and tapers gently towards the anterior end. The dorsal margin is close to straight, but bulges very slightly around the midlength.

**Fig. 14:**
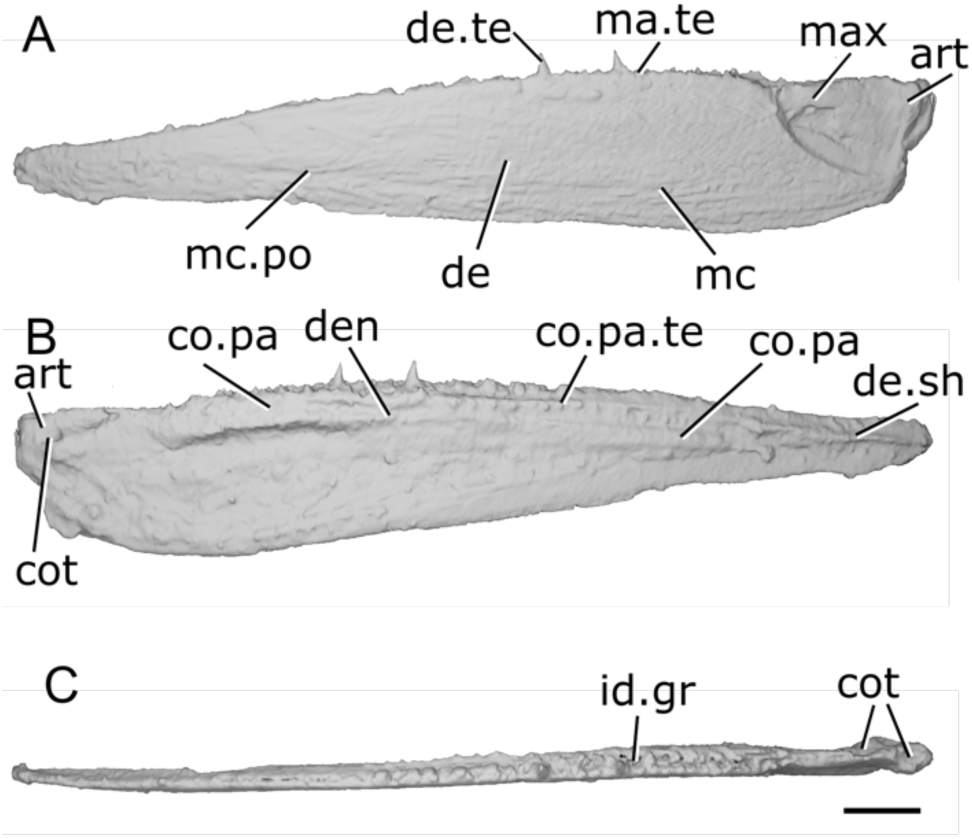
Mandible of *Kentuckia hlavini* CMNH. 9562**. a**, Left mandible in lateral view. **b**, medial view. **c**, dorsal view. Scale bar = 2 mm. *Abbreviations*: art, articular; co.pa, coronoid-prearticular; co.pa.te, coronoid-prearticular teeth; cot, cotyle; de, dentary; de.sh, dentary shelf; de.te, dentary teeth; den, denticles; id.gr, interdental groove; ma.te, marginal teeth; max, area overlapped by the maxilla; mc, mandibular canal; mc.po, pores connecting to the mandibular canal.

The lateral surface of the mandible has been interpreted as comprising a dentary and angular (Dunkle, 1964). However, a suture between the dentary and any postdentary bones is not apparent in tomograms, although this may be due to heavy ornamentation. The lateral surface of the mandible is depressed in the region of the maxillary overlap area (max, Fig. 14a) and there does not appear to be a separate surangular. This surface of this area is smooth and lacks ornamentation. This unornamented region is near-triangular, with the apex positioned posteroventrally. Its dorsal margin is straight, while the posterior margin is straight and almost vertical and the anteroventral margin is gently convex.

Although the heavy lateral compression of the specimen makes interpretation of the tomograms difficult, a distinct medial shelf of the dentary appears to be absent except in the anteriormost part of the mandible (de.sh, Fig. 14b).

Long ridges of ornament cover almost the entire surface of the mandible and mostly extend anteroposteriorly, although posteriorly they are oriented anterodorsally to posteroventrally. A small number of elongate, low tubercles are present close to the dorsal margin, at the midpoint of the length of the mandible, but otherwise the ridges are consistent in thickness and shape across the jaw.

Only two teeth appear to be completely preserved on the primary tooth row of the dentary (de.te, Fig. 14a). They are each straight, narrow, very sharp, and oriented dorsally. Empty sockets are evident along the mandible, and a series of stumps anterior to the two well-preserved teeth likely represent broken crowns. The size of the teeth appears to decrease anteriorly. Teeth generally alternate with empty replacement sockets, although this pattern is slightly irregular: two of the seven identified teeth are adjacent, and there are multiple adjacent sockets. Lateral to these, a series of short, sharp bumps likely represent a lateral row of marginal dentition (ma.te, Fig. 14a), confirmed by the description of two rows of dentary teeth by Dunkle (1964), but the scan is of insufficient resolution to be certain.

The mandibular canal (mc, Fig. 14a) traces the ventral margin of the jaw in the posterior third, but anterior to this is deflected upwards and gradually rises in a straight line to the dorsoventral midpoint of the jaw close to the anterior tip. Close to the anterior tip, the path of the canal levels out. Posteriorly, the position of the canal is only faintly indicated on the lateral surface. However, it is clearer in the anterior half, where it is marked by a line of small, dense and irregularly but closely spaced pores (mc.po, Fig. 14a) on the lateral surface. A small line of pits is also visible on the medial surface of the jaw, close to the dorsal margin of the lateral dermal ossification near the posterior end of the jaw, in the region of the lateral jaw surface that lacks ornamentation.

The medial dermal bone series (co.pa, Fig. 14b) spans from the posterior margin of the adductor fossa and appears to terminate at a point approximately a fifth of the way from the anterior margin of the mandible. The entire series is fused, so the number of coronoids and division with the prearticular cannot be discerned. Its lateral surface is very closely associated with the lateral dermal bones due to compression, and the underlying endochondrally ossified Meckel’s cartilage. The horizontal medial face consists of two surfaces: one mediodorsally facing and one medioventrally facing, separated by a sharp, medially oriented ridge. The medial dermal bones are separated from the dentary by a very narrow groove, but lateral compression makes the original width and extent of this gap difficult to determine.

A series of small, dorsomedially oriented teeth (co.pa.te, Fig. 14b) runs along the ridge separating the two faces of the medial dermal bone. They are widely spaced and restricted to the middle portion of the bone, and it is unclear if their orientation has been distorted during preservation. A scattering of much smaller denticles (den, Fig. 14b) is present more ventrally on the prearticular region.

Ossification of Meckel’s cartilage appears to be as separate articular and mentomeckelian elements. The mentomeckelian is limited anteriorly, although the full extent of the ossification is difficult to determine in the tomograms. In the articular region (art, Fig. 14a,b), Meckel’s cartilage forms two small cotyles (cot, Fig. 14b,c). The more lateral of the cotyles is positioned on the dorsal margin of the jaw. It is teardrop shaped, with the apex pointed posteriorly. Posterior to this, the jaw bulges slightly. The medial cotyle is positioned beneath the dorsal margin of the jaw, in a bulge on the medial surface of Meckel cartilage. It is oval shaped and faces dorsally and slightly medially. Lateral compression of the mandible has resulted in the adductor fossa becoming almost completely crushed, but it appears to have been very narrow and relatively elongate.

#### 3.4.9. Actinopterygii n. gen. n. sp. CMNH 9560

As indicated in Friedman & Blom (2006: pg. 1186), the Late Devonian Cleveland Member of the Ohio Shale yields a third, as-yet-unnamed actinopterygian taxon. This is represented by multiple isolated elements, as well as disarticulated but associated individuals, and is currently being formally described (Carr & Sallan, pers. comm.). An isolated right mandible that can be identified as belonging to this taxon on the basis of ornament and overall shape, CMNH 9560 (Fig. 15), is described here using X-ray CT. It has been laterally flattened during preservation, and some pyrite is present within the specimen.

**Fig. 15.**
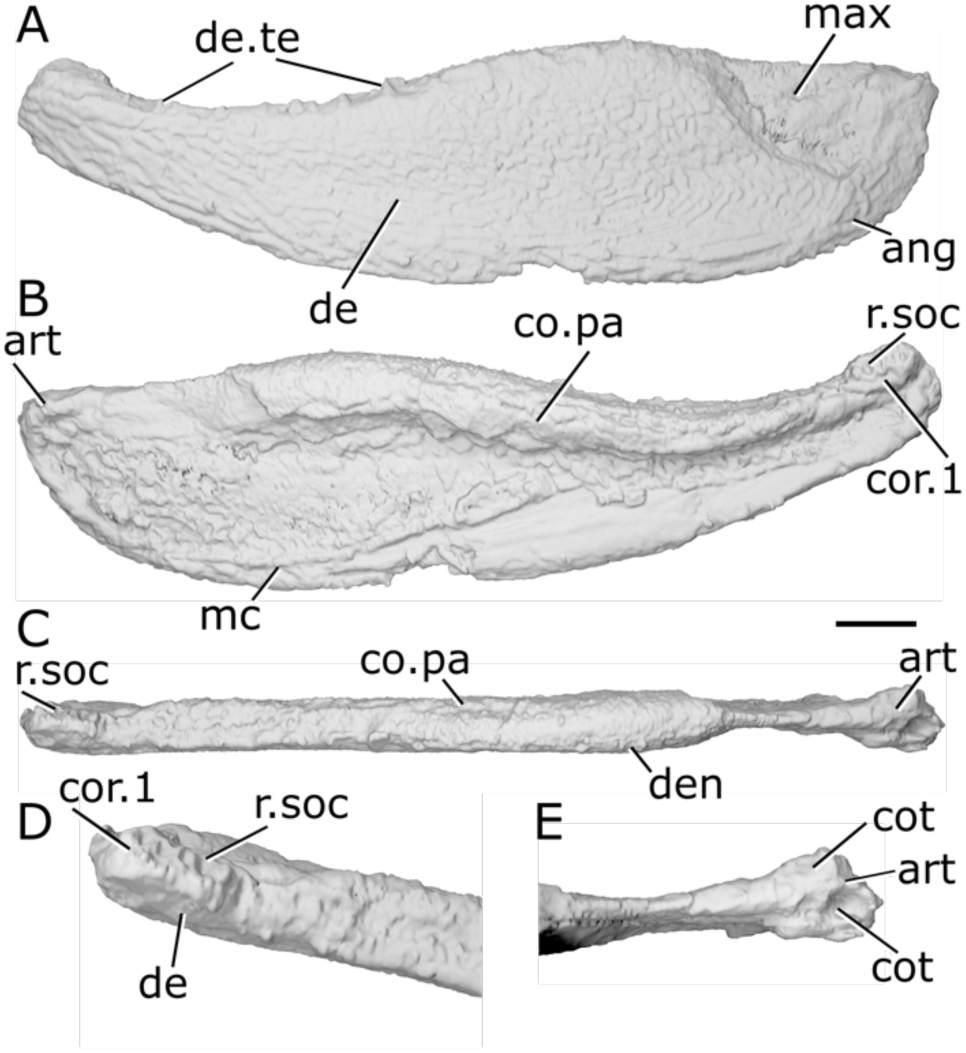
Mandible of Actinopterygii n. gen. n. sp. (CMNH. 9560**). a**, Right mandible (mirrored) in lateral view. **b**, medial view. **c**, dorsal view. **d**, anterior region in dorsal view. **e**, articular region in dorsal view. Scale bar = 2 mm. Panels d, e not to scale. *Abbreviations*: ang, angular; art, articular; co.pa, coronoid-prearticular; cor, coronoid; cot, cotyle; de, dentary; de.te, dentary teeth; den, denticles; max, area overlapped by the maxilla; mc, mandibular canal; r.soc, replacement socket.

The ventral margin is convex along its whole length, though more strongly towards the anterior and posterior ends. The dorsal margin of the jaw is sigmoid; it peaks with a gentle convex shape in the posterior half of the jaw, and the anterior half is strongly concave. The jaw is medially thickened at its anterior end, resulting in a bulbous tip. The adductor fossa extends for just under a quarter of the jaw in length, but due to the flattened nature of the specimen, little else can be said about its shape, including its dorsal profile.

The dentary (de, Fig. 15a,d) is the largest element of the lateral dermal jaw, forming almost the entire mandible depth along almost the entire length of the jaw. Its medial surface is developed as stubby, thickened medial shelf. The wedge-shaped angular (ang, Fig. 15a) forms the posteroventral jaw margin. There is no separate surangular. A thickened ridge for the mandibular canal (mc, Fig. 15b) runs along the posterior and ventral margin of the angular and ventral margin of the dentary, angling more sharply dorsally in the anterior half of the jaw.

The lateral surface of the jaw is strongly ornamented with irregular tubercles and pits. Close to the ventral margin of the jaw, the ornament is drawn out into longer ridges and grooves that run parallel to the ventral margin. The area that would have been overlapped by the maxilla in life (max, Fig. 15a) is depressed relative to the rest of the lateral surface and lacks ornament.

Short, sharp, conical teeth (de.te, Fig. 15a) are present at regular intervals along the dorsolateral margin of the dentary. Replacement sockets are difficult to identify due to pyrite infill along the jaw margin, but their presence can be inferred by larger gaps between some dentary teeth. Teeth appear to decrease in size towards the anterior margin of the dentary, and are entirely absent from the anteriormost portion, lateral to the expanded coronoid. There is no evidence for marginal dentition lateral to the main dentary toothrow.

The medial tooth-bearing dermal bones appear to be co-ossified as a single series (co.pa, Fig. 15b,c). They are consistent in depth along the jaw and do not deepen posteriorly, being restricted to near the dorsal margin of the jaw and following its sinusoidal outline. The plate has an inflated dorsal margin that wraps around the medial shelf of the dentary and reaches as high as the tips of the dentary teeth. Much of the dorsal and some of the medial surface of the coronoid-prearticular plate is covered with minute, rounded denticles that forms a continuous field of dentition. As the dorsal extension of the coronoid-prearticular plate completely overlies the medial shelf of the dentary and abuts the dentary toothrow, there is no interdental groove. Anteriorly, in the region likely corresponding to the anterior coronoid (cor.1, Fig. 15b,d), the medial tooth-bearing series broadens towards the midline and its upper surface becomes dorsomedially directed. A row of four-five empty tooth sockets (r.soc, Fig. 15b,d), which indicate the presence of teeth of a similar size to those borne on the dentary, are borne on this surface. Minute denticles (den, Fig. 15c) line a ridge lateral to this empty row.

Meckel’s cartilage is ossified in at least the articular region of the jaw (art, Fig. 15b,c,e), but is difficult to trace further anteriorly. Two cotyles (cot, Fig. 15e) are present on the articular, although laterally compressed. The more medial cotyle is deep, oval, and faces dorsally and slightly laterally. The medial more cotyle is also oval, though is smaller and positioned further anteriorly. It faces dorsomedially. The mandibular canal (mc, Fig. 14b) lies close to the ventral margin of the jaw and runs parallel to it along the entire length of the jaw.

#### 3.4.10. Tegeolepis clarki

The mandible of *Tegeolepis clarki* has previously been described by Gardiner (1963) and Dunkle & Schaeffer (1973) based on examination of the exposed lateral surface, and by Figueroa et al. (2021) based on X-ray CT imaging of a partial mandible. We supplement these descriptions based on reexamination of the surface files of CMNH 8124 (Fig. 16a-c,g) produced by Figueroa et al. (2021) and new X-ray CT imaging of NHMUK PV P 45312 (Fig. 16d-f,h,i). The latter mandible (173 mm long) is from a complete, articulated, though heavily laterally compressed cranium, belonging to the same individual as a postcranium cataloged as a separate specimen (NHMUK PV P 9402). In addition to postmortem compression, the skull is very dense, fractured and contains x-ray dense mineral inclusions, so the level of detail that can be drawn from the tomograms is limited.

**Fig. 16:**
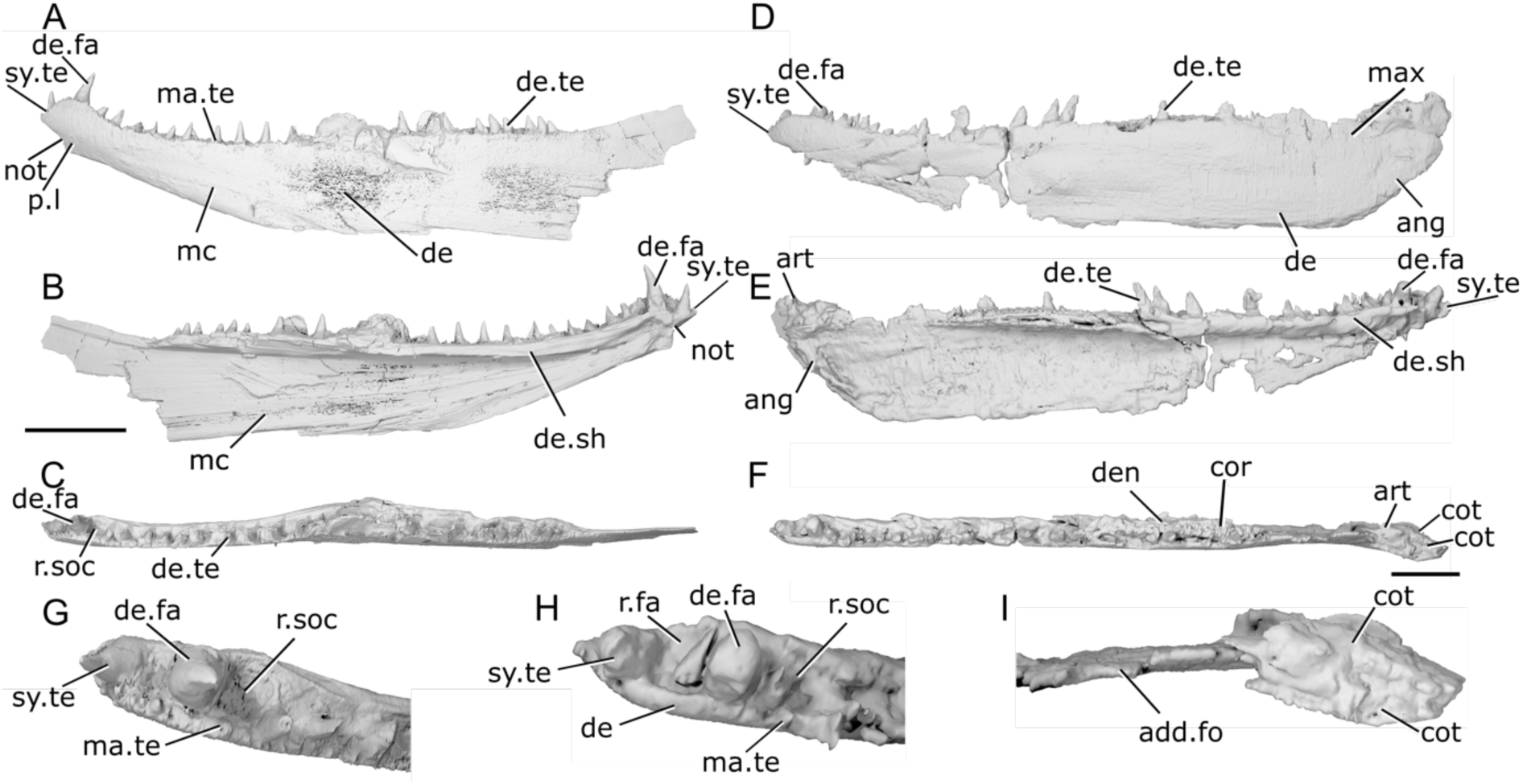
Mandibles of *Tegeolepis clarki*. **a**, Right mandible (mirrored) of CMNH 8124 in lateral view. **b**, mandible of CMNH 8124 in medial view. **c**, mandible of CMNH 8124 in dorsal view. **d**, right mandible (mirrored) of NHMUK PV P 45312 in lateral view. **e**, mandible of NHMUK PV P 45312 in medial view. **f**, mandible of NHMUK PV P 45312 in dorsal view. **g**, anterior region of CMNH 8124 in dorsal view. **h**, anterior region of NHMUK PV P 45312 in dorsal view. **i**, articular region of NHMUK PV P 45312 in dorsal view. Scale bar: a-c = 20 mm. d-f = 15 mm. Panels g, h, i not to scale. *Abbreviations*: ang, angular; art, articular; cor, coronoid; cot, cotyle; de, dentary; de.fa, dentary fang; de.sh, dentary shelf; de.te, dentary teeth; den, denticles; ma.te, marginal teeth; max, area overlapped by the maxilla; mc, mandibular canal; not, notch; p.;, pit line; r.fa, replacement fang; r.soc, replacement socket; sy.te, symphysial teeth.

The dorsal and ventral margins are straight and parallel along much of the jaw. In the posterior half of the jaw, the margins are near-horizontal, with the posterodorsal margin sloping upwards at a roughly 45-degree angle. In the anterior half of the mandible, the ventral margin turns sharply anterodorsally. The dorsal margin also slopes upwards with a concave margin in its anterior third. The jaw bulges in thickness at its anterior tip; this is more pronounced in CMNH 8124, which also bears a concavity (not, Fig. 16a,b) on the anteroventral margin of the mandible. The adductor fossa (add.fo, Fig. 16e,i) extends for approximately 20% of the length of the jaw. Posteriorly, the articular region (art, Fig. 16e,f,i) is slightly raised relative to the rest of the mandible, sitting level with the top of the largest dentary teeth.

The dentary (de, Fig. 16a,d,h) comprises most of the lateral surface of the jaw. Its medial surface bears a robust, extensive medial shelf (de.sh, Fig. 16b,e; the ‘internal longitudinal laminae’ of Dunkle & Schaeffer, 1973: p. 157) with a slightly convex dorsal surface. The shelf tapers out posteriorly, but anteriorly becomes deeper and flattens against the medial surface of the dentary. A ventral ridge is present along the anterior third of the mandible. It may have carried the mandibular canal (mc, Fig. 16a,b), and merges with the anterior extremity of the medial dentary shelf to form a sideways ‘v’ shape. A crescentic angular (ang, Fig. 16d) is present along the posterior and part of the ventral jaw margin. The angular is narrow along its whole length, projecting anteriorly level with the midpoint of the adductor fossa. We cannot corroborate presence of the surangular reported by Gardiner (1963: p.302), but this may be an artefact of scan quality.

Ornamentation is only visible on the surface of CMNH 8124, and comprises extremely fine tubercles and anteroposteriorly elongate ridges. A small, shallow depression lateral to the adductor fossa of NHMUK PV P 45312 (max, Fig. 16d) indicates the area that would have been overlapped by the maxilla.

Twenty one large, robust teeth (de.te, Fig. 16a,c,d,e) are preserved on the dentary, with their bases borne on the medial dentary shelf. They are generally straight and oriented vertically, but some of the teeth in the anterior half of the mandible of both specimens slope posteriorly, and the posterior six teeth in CMHN 8124 slope anteriorly. These six posteriormost teeth are also smaller and more closely spaced than other teeth on the dentary. The anteriormost tooth, situated on the jaw symphysis (sy.te, Fig. 16a,b,d,e,g,h) is squatter in form and faces anteriorly. The second tooth in CMNH 8124 and third tooth in NHMUK PV P 45312, which is situated on the thickened portion of the dentary, is fang-like (de.fa, Fig. 16a,b,c,d,e,g,h). It is twice the size of the teeth immediately posterior to it in both height and width. The third tooth position in CMNH 8124, and second tooth position in NHMUK PV P 45312, is open as a replacement socket (r.soc, Fig. 16g,h), and a minute replacement fang (r.fa, Fig. 16h) is visible in the latter specimen. Along much of the toothrow, replacement sockets alternate with teeth. However, the posterior six teeth form two short series of three consecutive teeth, with a single replacement socket in between in CMNH 8124, and in NHMUK PV P 45312 there are also several adjacent sockets at the posterior end of the row. A single row of marginal dentition (ma.te, Fig. 16a,g,h) is present along the lateral surface of the dentary, lateral to the main toothrow. These teeth are minute, sharp, and oriented dorsally.

The medial tooth-bearing series is not preserved in CMNH 8124, and only a single coronoid element (cor, Fig. 16f) can be confidently identified in NHMUK PV P 45312. It is narrow and elongate, and sits entirely on top of the medial shelf of the dentary, as in *Cheirolepis* (Fig. 3b,f,g). The coronoid is positioned very close to the dentary toothrow, and as such there is no interdental groove. Its dorsal and medial surface is covered with small, pointed, randomly arranged denticles (den, Fig. 16f), other than an area at the anterior margin that may have contacted a neighboring coronoid.

The path of the mandibular canal (mc, Fig. 16a,b) cannot be traced fully; it is evident at the posterior end of the jaw in NHMUK PV P 45312 and the anterior third in CMNH 8124. In NHMUK PV P 45312, a thickened ridge in the angular and posterior end of the dentary indicate that the mandibular canal was very close to the posteroventral jaw margin in this area. In CMNH 8124, a short line of approximately six very small pores is present close to the ventral margin of the dentary on its lateral surface, positioned roughly in line with the positions of teeth seven and eight. The pores form a line that slopes gently anterodorsally to posteroventrally. These are aligned with a faint groove close to the ventral margin of the jaw. Close to the anterior margin of the mandible, beneath the largest dentary tooth, the mandibular canal turns sharply dorsally into a curved pit line (p.l, Fig. 16a), reminiscent of that seen in *Mimipiscis toombsi* (Fig. 7A).

Meckel’s cartilage is only ossified posteriorly, in the articular region (art, Fig. 16e,f,i), and even here ossification does not extend past the middle of the adductor fossa. It is thickened on the medial surface, forming a ridge down the posterodorsal margin of the dentary to support the two cotyles (cot, Fig. 16f,i). The lateral cotyle is ovoid and faces dorsally. The medial cotyle is also ovoid and faces dorsally, but is positioned more ventrally.

#### 3.4.11. Limnomis delaneyi

The mandible of *Limnomis delaneyi* has been previously described by Daeschler (2000) based on surface examination. We supplement this description through X-ray CT of ANSP 23721, an articulated skull containing both lower jaws. Although most specimens of *Limnomis* are preserved as highly flattened within green-to-gray mudstones, this specimen is preserved in a coarser reddish matrix and is less severely flattened. The left mandible is complete except for its posteroventral corner and parts of the ventral margin (preserved length: 8.2 mm). The right mandible is complete posteroventrally but the anterior tip is missing; the dentary is also fractured in several places and has been retrodeformed (Fig. 17, Fig. S3). In both specimens, the coronoids are slightly disarticulated and sit dorsal to life position.

**Fig. 17:**
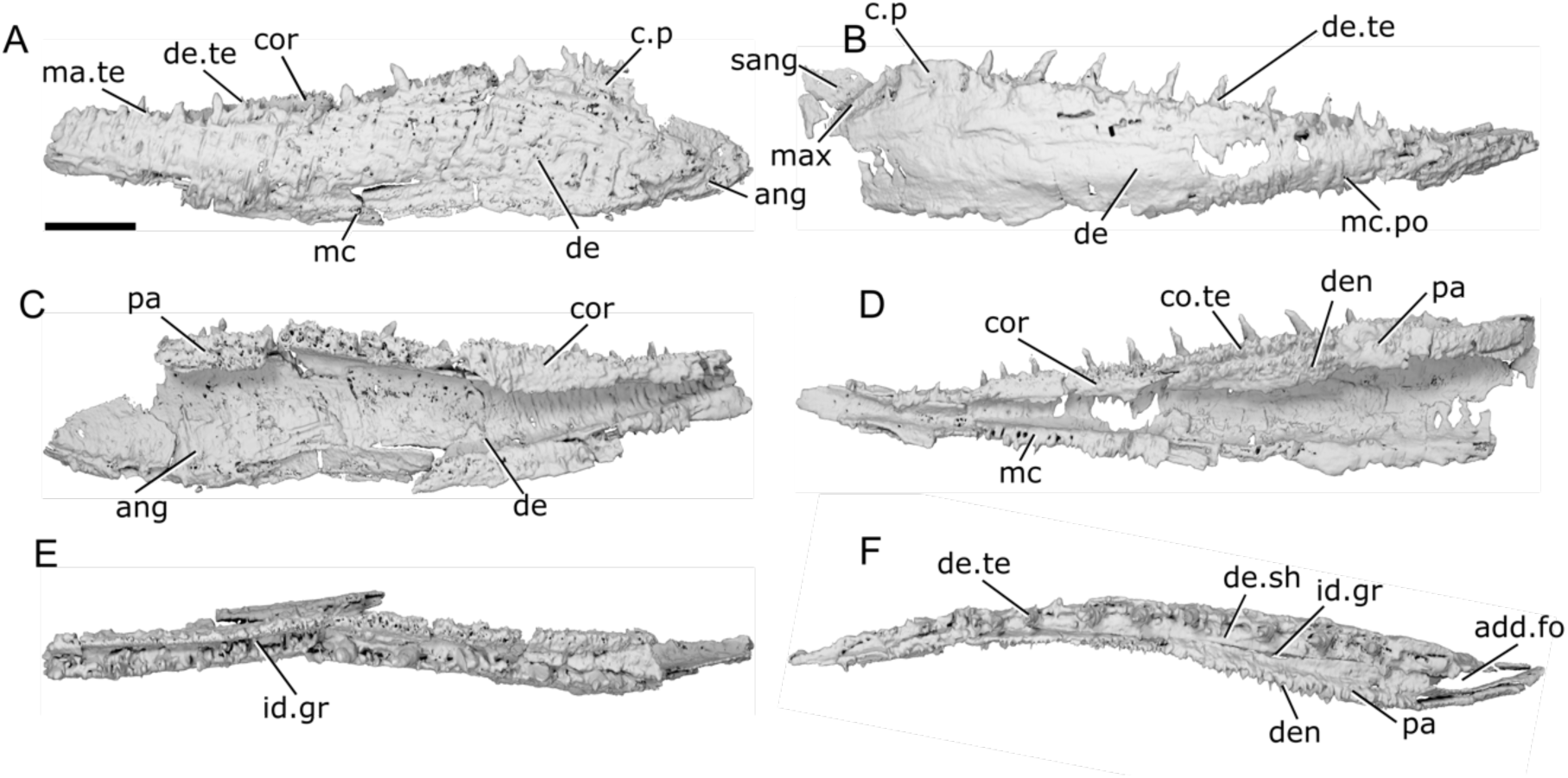
Mandibles of *Limnomis delaneyi* ANSP 23721. **a**, Left mandible in lateral view. **b**, right mandible in lateral view. **c**, left mandible in medial view. **d**, right mandible in medial view. **e**, left mandible in dorsal view. **f**, right mandible in dorsal view. Scale bar = 1 mm. *Abbreviations*: add.fo, adductor fossa; ang, angular; c.p, coronoid process; cor, coronoid; co.te, coronoid teeth; de, dentary; de.sh, medial dentary shelf; de.te, dentary teeth; den, denticles; id.gr, interdental groove; ma.te, marginal teeth; max, area overlapped by the maxilla; mc, mandibular canal; mc.po, pores connecting to the mandibular canal; pa, prearticular; sang, surangular.

The ventral margin of the mandible is very gently convex, and the dorsal margin is very gently concave. The jaw is deepest posteriorly, at the level of the adductor fossa, and tapers anteriorly with a gentle upturn at the anterior margin. In dorsal view, the posterior half and anterior quarter of the jaw are straight, but the jaw curves strongly medially in between these sections.

Most of the lateral surface of the jaw is made up of the dentary (de, Fig. 17a,b,c), which terminates posteriorly in a rounded triangular point. The dentary forms the deepest part of the mandible, and, along with a portion of the surangular, forms a broad, low coronoid process (c.p, Fig. 17a,b). A concave depression in the posterior half of this region indicates the area overlapped by the maxilla (max, Fig. 17b). The dentary bears a robust but shallow medial shelf (de.sh, Fig. 17f) that partially underlies the medial tooth-bearing bones. A deep, ovoid angular (ang, Fig. 17a,c) is partially overlapped by the dentary and forms the posterior and ventral margin of the dentary. It has an elongate angular arm that extends approximately halfway along the length of the dentary, narrowing slightly anteriorly. Unlike in generalized Devonian actinopterygians such as *Gogosardina*, there is no posterodorsal arm to the angular: instead, its dorsal extent reaches barely halfway up the dentary and the mandibular canal exits from its posteroventral margin. A small surangular (sang, Fig. 17b) is present, something noted as uncertain by Daeschler (2000). However, it has shifted somewhat ventral to life position and was likely originally situated above the angular.

The ornament is only partially visible, as a combination of exposure of the surface of the specimen and mineral inclusions within the fossil have obscured large parts of the lateral surface of both mandibles. It is best preserved in the posterior region of the left mandible, and is formed of elongate, broad ridges. These are oriented anterodorsal-posteroventrally in the posterodorsal portion of the jaw, and become aligned with the ventral margin more ventrally.

The dentary toothrow comprises large, narrow, sharp teeth that curve slightly medially (de.te, Fig. 17a,b,f). The teeth become smaller anteriorly. They generally alternate with replacement sockets, with small replacement teeth visible in some sockets. The teeth are slightly recessed from the lateral margin of the dentary, with their bases situated on the medial shelf. An additional row of small, more numerous pointed teeth (ma.te, Fig. 17a) are present lateral to the main dentary toothrow.

The dermal medial tooth bearing bones are preserved in both jaws, but are more complete on the right. Coronoids and the prearticular are separately ossified on both sides. The entire series comprises a flat, medially facing surface and a narrow lateral shelf that overlaps the corresponding medial shelf of the dentary. The lateral shelf increases in width posteriorly. It is widest just anterior to the adductor fossa, at which point a lateral projection curves around the front of the fossa. As a result, the interdental groove (id.gr) between the teeth of the lateral and medial teeth bearing bones increases in width posteriorly. The right jaw preserves at least three coronoids (cor, Fig. 17a,c,d), with a possible fourth preserved at the preserved anterior limit of the mandible. The left jaw is less completely preserved, but at least two coronoids are present. Each is connected to its neighbour via a jagged suture, with a narrow projection on the posterior margin underlying the more posterior coronoid. A similar feature connects the posteriormost coronoid and the prearticular. The prearticular (pa, Fig. 17c,d,f) is long, spanning just under half the length of the mandible, and is slightly deeper than the coronoids. It forms the anterior, most of the medial, and a small part of the margins of the adductor fossa (add.fo, Fig. 17f).

Two rows of moderately sized teeth are present on the dorsal edge of the medial surface of the prearticular. They are very closely spaced, with medially directed tips. A groove ventral to these two rows of teeth continues anteriorly into the coronoid series, but gradually tapers out approximately halfway along the way. Only a single row of teeth is present above the groove on the coronoid elements. Beneath the groove, a dense field of denticles, which generally decrease in size ventrally, extends down the prearticular and coronoids.

There is no evidence in either jaw of any ossification of Meckel’s cartilage, either in the mentomeckelian or articular regions. Consequently, the shape and condition of the articular facets is unknown.

The mandibular canal (mc, Fig. 17a,d) can be traced along much of the ventral margin of the mandible. It was housed in a large, thin-walled canal and connected to the lateral surface of the jaw by regularly spaced pores (mc.po, Fig. 17b). the position of these pores indicates that the mandibular canal did not arch anteriorly, instead remaining close to the ventral margin, and that it reached the anterior end of the jaw.

#### 3.4.12. “Gonatodus” brainerdi

The genus *Gonatodus* was established for material from the early Carboniferous of Scotland previously assigned to *Amblypterus* (for a review, see Gardiner, 1967: p. 147). The Devonian form reviewed here was originally identified as a new species of *Paleoniscum* (Thomas, 1853) and subsequently placed in *Gonatodus* by Newberry (1889) with no clear justification. This assignment is almost certainly incorrect, so we refer to the Devonian species as “*Gonatodus*” *brainerdi* here.

The lower jaw of “*Gonatodus*” *brainerdi* has not been previously described. The description here (Fig. 18) is based on ANSP 6232, an articulated specimen (mandible length: 35 mm) that has undergone lateral compression. The dorsal margin of the lower jaw is almost completely straight, with a slight convexity along the posterior half of the mandible. Posterior to the dentary toothrow, the jaw margin rises just above the tips of the dentary teeth, forming a long, low coronoid process (c.p, Fig. 18a,b) with a flattened dorsal margin. The posterior margin of the lower jaw is relatively straight, and close to vertical.

**Fig. 18:**
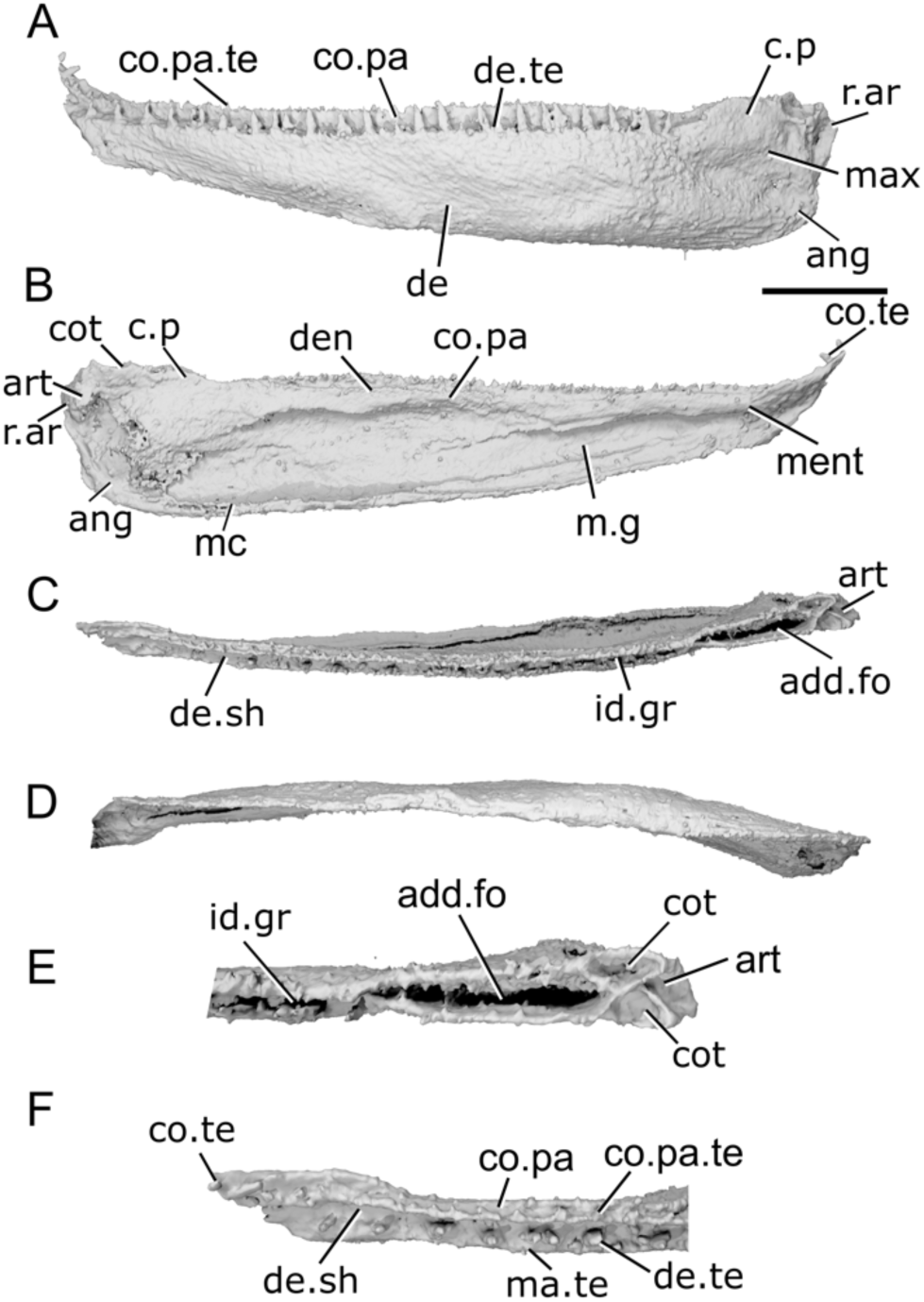
Mandible of *Gonatodus brainerdi* ANSP. 6232**. a**, Left mandible of ANSP 6232 in lateral view. **b**, medial view. **c**, dorsal view. **d**, ventral view. **e**, articular region in dorsal view. **f**, anterior region in dorsal view. Scale bar = 5 mm. Panels e,f not to scale. *Abbreviations*: add.fo, adductor fossa; ang, angular; art, articular; co.pa, coronoid-prearticular; co.pa.te, coronoid-prearticular teeth; co.te, coronoid tooth; cot, cotyle; c.p, coronoid process; de, dentary; de.sh, dentary shelf; de.te, dentary teeth; den, denticles; id.gr, interdental groove; m.g, groove housing the mandibular canal; ma.te, marginal teeth; max, area overlapped by the maxilla; ment, mentomeckelian ossification; r.ar, retroarticular process.

The ventral margin of the jaw is straight and horizontal in its posterior half and rises gently from the midpoint. The anterior tip of the lower jaw turns sharply upwards and medially, and tapers to a point. A deep, bowl-shaped concavity on the lateral surface of the posterior region of the jaw corresponds to the overlap area of the maxilla (max, Fig. 18a). The jaw gently curves along its length in dorsal view.

Sutures are difficult to identify because of the strongly developed dermal ornament, but the lateral surface of the mandible appears to comprise the dentary and a single infradentary, the angular. The dentary (de, Fig. 18) is by far the largest component, with the angular (ang, Fig. 18a,b) restricted to a very narrow strip along the posterior and ventral margins of the jaw; the posteroventral corner of the angular projects slightly posterior to the more dorsal portion of the jaw margin. The medial shelf of the dentary (de.sh) is barely developed, and it abuts the nearly flat lateral face of the inner dental series. A narrow but very deep interdental groove (id.gr, Fig. 18e) is present along the entire mandible.

Much of the lateral surface of the lower jaw is ornamented with a series of long, shallow, parallel grooves. No pits or pores are evident on the lateral surface. In the ventral half of the jaw, the grooves run approximately parallel with the ventral lower jaw margin. In the dorsal half of the jaw, they slope anterodorsally to posteroventrally. No ornamentation is present in the region overlapped by the maxilla (max, Fig. 18a).

A row of closely spaced teeth (de.te, Fig. 18a,f) is present along the dorsal margin of the dentary, positioned very close to its lateral edge. Approximately twenty-two teeth are present, all vertically oriented, tall, and narrow. They are mostly consistent in size, though the anterior and posterior-most teeth are slightly smaller. The teeth generally alternate with replacement sockets, but there is one pair of adjacent teeth, and three pairs of adjacent replacement sockets. Additionally, multiple smaller pointed teeth (ma.te, Fig. 18f) are positioned lateral to the main toothrow, right on the lateral edge of the jaw.

The medial dermal tooth-bearing bones are co-ossified as a single series (co.pa, Fig. 18a,b,f), so the number of coronoids cannot be determined. The medial dermal complex is shallow and restricted to the anterior half of the mandible, although it deepens slightly medial to the adductor fossa. Along most of its length, this element is formed of a tall vertical surface extends dorsally as high as the tops of the dentary teeth. A shallow groove and corresponding medial ridge approximately a third of the way from the top of the shelf separates it into dorsal and ventral regions. The dorsal region is narrow, and bears a single row of long, sharp, medially-directed teeth (co.pa.te, Fig. 18a,f) along its upper surface. These are closely spaced and become shorter and more medially directed near the anterior margin of the dentary. One to two irregular rows of denticles (den, Fig. 18b) leave the ventral region of the prearticular-coronoid series immediately ventral to the ridge. These cusps are more rounded than those in the upper region, becoming more so anteriorly. Scattered denticles are also present in the middle of the deeper, prearticular plate. In the anteriormost region of the mandible, the ridge and groove separating the dorsal and ventral regions of the coronoid-prearticular series twists dorsally such that the tooth rows of the two region become parallel. The inner dental series continues anterior to the anterior margin of the dentary and turns sharply dorsally and laterally: the ridge between the two regions is visible in lateral view in this part. A single, moderately-sized tooth (co.te, Fig. 18b,f) is present on the anteriormost tip of what likely corresponds to the anteriormost coronoid (cor.1), its tip pointing posteriorly.

The mandibular canal (mc, Fig. 18b) is only fully enclosed within the angular, where it runs along its posterior margin in a very narrow canal. It exits the posterior of the mandible approximately halfway up on the posterior margin, where the posterior extension of the angular begins to narrow again. There are no pores on the lateral surface of the dentary, although several are present within the angular. No fully enclosed mandibular canal is evident in tomograms of the lower jaw, nor are there any pores on the lateral surface of the lower jaw that might mark the path of a mandibular canal. There is, however, a low but sharp ridge running along much of the medial surface of the dentary in the lower half. This ridge results in a deep groove on the medial surface which may have housed a canal m.g, Fig. 18b). The groove continues until almost the posterior end of the jaw.

Meckel’s cartilage is unossified throughout much of the jaw. A few patches of perichondral bone right at the anterior end of the jaw represent the mentomeckelian ossification (ment, Fig. 18b). Meckel’s cartilage is also ossified in the region of the articular (art, Fig. 18b,c,e). It does not extend past the posterior end of the adductor fossa, although its rear margin bulges behind the rest of the jaw margin. The cartilage is mostly ossified as perichondral bone, but small amounts of endochondral bone are present sporadically throughout the element. Two cotyles (cot, Fig. 18b,e) are present on the dorsal surface of the articular. The more lateral cotyle is hemispherical and faces dorsolaterally. The more medial is positioned more dorsally and slightly anteriorly. It is deeper and more elongate than the lateral cotyle, and the medial-more cotyle is more elongate, and forms a very deep groove. It is positioned higher and further forwards than the lateral-more cotyle, and oriented dorsally. Behind the cotyles, a short, shallow groove extends laterally across the articular. Though the jaw has been flattened, the adductor fossa is clearly narrow and elongate, becoming narrower towards its anterior end.

#### 3.4.13. Cuneognathus gardineri

The mandible of *Cuneognathus delaneyi* has previously been described by Friedman & Blom (2006) based on external description of specimens preserved in lateral view. We add to this description based on X-ray CT of NHMD-1235389 (Fig. 19), a complete but laterally compressed cranium. Both the left and right mandibles are preserved (each 12 mm in length), but compression has led to them appearing straight in dorsal view. The ventral margin of the left jaw and ventral portion of the medial surface of the right jaw are also partially incomplete where they are exposed on the surface.

**Fig. 19:**
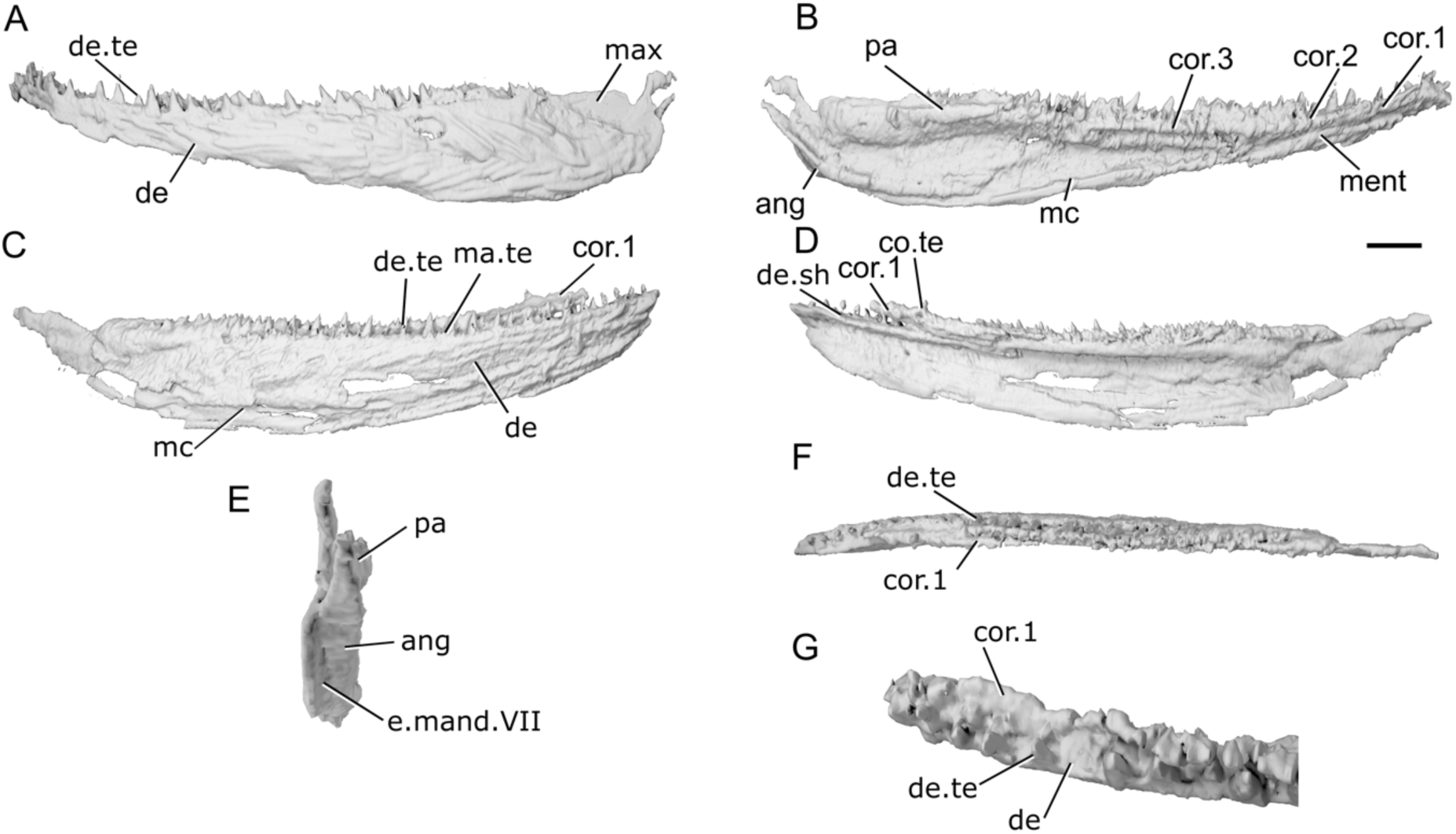
Mandibles of *Cuneognathus gardineri* NHMD-1235389. **a**, left mandible in lateral view. **b**, left mandible in medial view. **c**, right mandible in lateral view. **d**, right mandible in medial view. **e**, left mandible in posterior view. **f**, right mandible in dorsal view. coronoid region of right mandible in dorsal view. **g**, anterior region of left mandible in dorsal view. Scale bar a-d, f = 1 mm. Panels e, g not to scale. *Abbreviations*: ang, angular; cor, coronoid; co.te, coronoid teeth; de, dentary; de.sh, dentary shelf; de.te, dentary teeth; e.mand.VII, external mandibular branch of the facial nerve; max, area overlapped by the maxilla; ma.te, marginal teeth; mc, mandibular canal; ment, mentomeckelian ossification; pa, prearticular.

The dorsal margin of the mandible is mostly straight and horizontal. It begins to curve very gently dorsally in its anterior third, curving most strongly dorsally at the anterior tip of the jaw. The ventral margin is deepest just anterior to the adductor fossa. Posterior to the deepest point, it curves strongly dorsally. Anterior to this point, it curves very gently dorsally for the rest of the jaw length. A small area posteriorly was overlapped by the maxilla (max, Fig. 19a). This reaches almost to the posterior end of the jaw, and was triangular in shape with the apex facing posteroventrally. In dorsal view, the jaw curves very gently medially, but this is likely less strong than it would have been in life due to taphonomic flattening of the specimen.

Ornamentation is present across the entire lateral surface of the dentary. It consists of long, wavy ridges. The ridges are strongest in the posteriorly half of the jaw, where they show two orientations. In the ventral half of the jaw the ridges are horizontal, and in the dorsal half of the jaw they are oriented diagonally, sloping from anterodorsal to posteroventral. In the anterior half of the jaw, the ornamentation is lighter, and has no particular pattern of orientation. Specimen photos show that the ridges break up into tubercles in the dorsalmost part of the jaw (Friedman & Blom 2006: fig. 3A).

The dentary (de, Fig. 19a,c,) makes up the entire lateral surface of the mandible of *Cuneognathus*. It bears a medial shelf (de.sh, Fig. 19b,d) level with its dorsal margin that partially underlies the medial tooth-bearing dermal bones. This shelf flattens against the medial surface of the dentary posteriorly, and tapers out some distance anterior to the adductor fossa, before the prearticular. Friedman & Blom (2006) were not able to identify an angular, but tomograms show that it is present (ang, Fig. 19b) along the very posterior margin of the mandible and is barely visible in lateral view. The angular is narrow along its length, primarily formed as a tube surrounding the mandibular canal (mc, Fig. 19b,c), and extends anteriorly as far as the adductor fossa before terminating abruptly. Further anteriorly, the mandibular canal is carried within a narrow canal along a ventral ridge at the ventralmost edge of the dentary. Its path through the anterior quarter of the jaw is uncertain, but it may have been exposed above a ventral ridge as in “*Gonatodus*” *brainerdi* (Fig. 18b). A few small pores, indicating the presence of the mandibular canal, are present at the posterior end of the jaw, some posteromedially on the angular and some lateral on the dentary.

The dentary possesses a single row of sharp, vertically oriented, conical teeth (de.te, Fig. 19a,c,f,g) in the anterior two-thirds of the jaw, situated on its dorsal surface. The row includes approximately eighteen teeth in the right jaw, and twenty in the left. A row of numerous, irregularly spaced small teeth (ma.te, Fig. 19c) lie lateral to the main tooth row. These are present along the entire margin of the jaw, including lateral to the anterior portion of the adductor fossa where larger teeth are absent. Most of the dentary tooth row consists of evenly spaced teeth, with one tooth alternating with one replacement tooth socket. However, there are a small number of adjacent teeth or replacement sockets. Additionally, a continuous row of five slightly smaller teeth are present at the anterior end of the left jaw, continuing right to the anterior tip of the dentary.

The medial tooth-bearing series is present as separately ossified prearticular and coronoids, all restricted to the dorsal portion of the mandible along the entire length of the jaw. The prearticular (pa, Fig. 19b) reaches almost to the posterior margin, terminating just anterior to the angular, and extends forward for approximately one-third of the jaw length. It consists of a single, bulbous shelf. Three separate coronoids (cor, Fig. 19b) are present, each roughly equal in length, extending from the anterior tip of the prearticular to almost the anterior tip of the mandible. Each consists of a single curved shelf, very closely applied to the dentary with no gap in between, though it is possible that this is a result of the extreme lateral compression of the jaws. Each coronoid partially overlaps the element posterior to it. Teeth and denticles are absent from the prearticular until its anterodorsal corner, although the apparent lack of denticles may be an artefact of scan resolution. A single row of teeth (co.te, Fig. 19d), slightly smaller than the dentary teeth, runs along the dorsalmost part of the prearticular and onto the third and second coronoid. The third and second coronoids also possess a more ventral field of additional, slightly smaller, randomly placed teeth. The tooth-bearing surface of the anteriomost coronoid (cor.1, Fig. 1b,d,f,g) faces dorsally, rather than dorsomedially. A single row of three-four teeth sit on this surface, parallel to the dentary tooth row. The anteriormost is larger, and its tip points posteriorly. No denticles are visible, but as with the prearticular, this may be an artefact of scan resolution.

Meckel’s cartilage is ossified as a mentomeckelian (ment, Fig. 19b) along the anterior quarter of the jaw length. There is no endoskeletal ossification in the posterior region of either jaw, so it is not possible to describe the articular region or cotyles.

## 4. DISCUSSION

### 4.1. Morphological variation in Devonian actinopterygian mandibles

Variation in the morphology of the mandible of early actinopterygians, and its potential utility for informing relationships—as well as ecological role—has long been recognized. Seminal papers that laid out phylogenetically informed arguments recognised trends in the ossification of Meckel’s cartilage, the number of external dermal elements, and the coronoid process (e.g. Gardiner, 1984; Gardiner & Schaeffer, 1989; Gardiner et al., 2005). Many qualitative observations have been used to feed into diagnostic treatments of taxa (e.g. Dunkle & Schaeffer, 1973) and to support informal higher taxonomic groupings (e.g. Gardiner & Schaeffer, 1989), as well as to suggest possible affinities for isolated mandibular material (e.g. Friedman & Blom, 2006; Figueroa et al., 2021). As a result, an appreciable number of characters in recent phylogenetic character-by-taxon matrices capture mandibular morphology.

The anatomical information reported here (Figs 20, 21) allows us to clarify and revise morphological character-state data for many early actinopterygians, as well as to propose additional characters to capture newly recognized variation. We review these traits below by updating the lower jaw characters from a recent character-by-taxon matrix targeting early osteichthyans, particularly actinopterygians (Giles et al., 2023). Although we do not attempt a mandible-based phylogenetic analysis (due to acknowledged problems with restricted character sets), we anticipate that future studies may incorporate some or all of these traits as discrete characters.

**Fig. 20.**
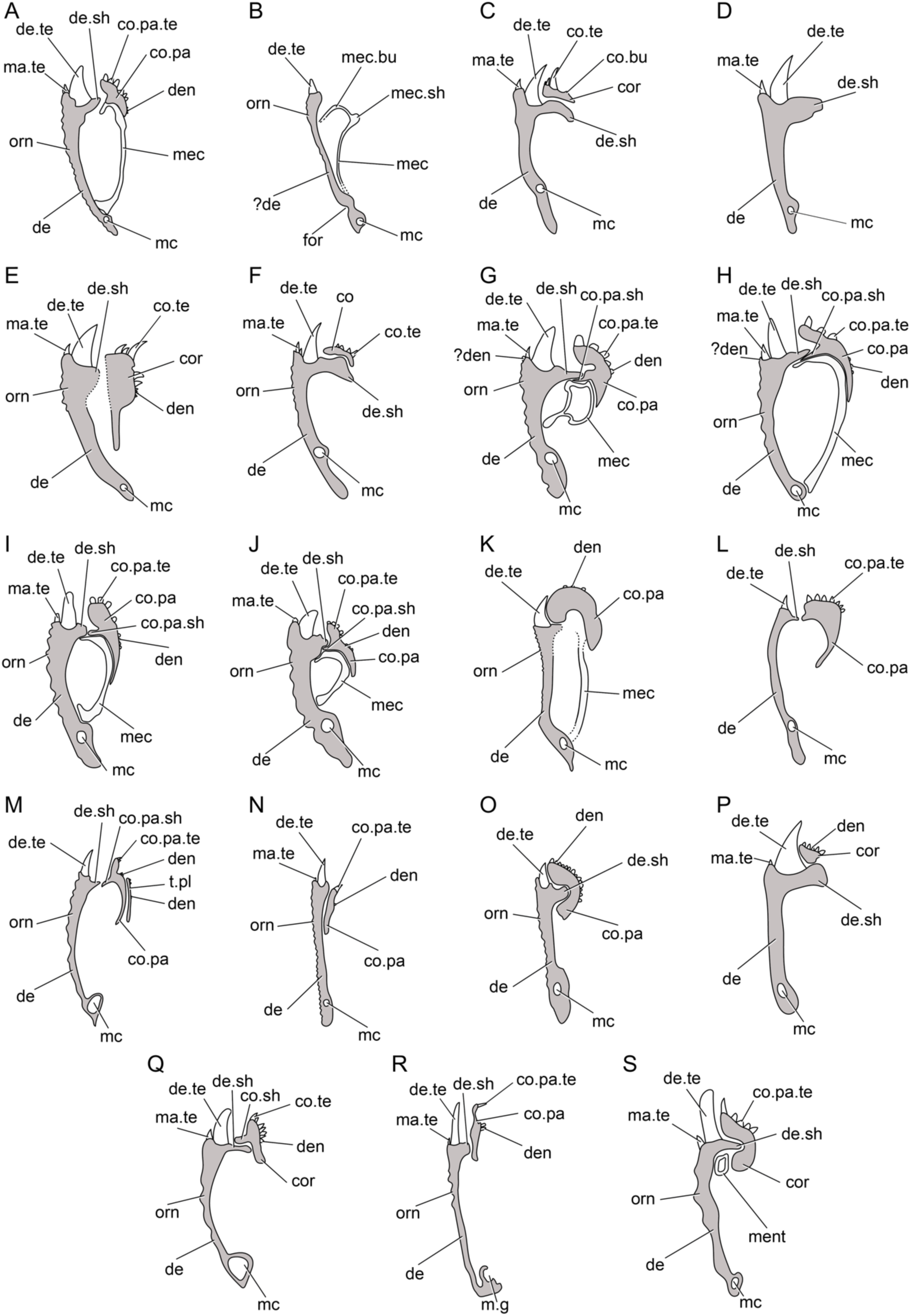
Schematic axial sections through mandibles of all taxa included in this study, positioned anterior to the midpoint of the jaw. a, *Gogosardina coatesi*. b, *Meemannia eos*. Internal dermal bones unknown. c, *Cheirolepis trailli*. Only articular region of Meckel’s cartilage ossified. d, *Austelliscus ferox*. Internal dermal bones and Meckel’s cartilage unknown **e**, *Howqualepis rostridens*. Only articular region of Meckel’s cartilage ossified; inner surface of jaw unknown. **f**, *Cheirolepis jonesi.* Meckel’s cartilage unossified. g, *Mimipiscis toombsi*. h, *Mimipiscis bartrami.* i, *Moythomasia durgaringa*. j, *Raynerius splendens*. **k**, ?*Moythomasia devonica*. l, *Osorioichthys marginis*. Only mentomeckelian and articular region of Meckel’s cartilage ossified. m, *Palaeoneiros clackorum*. Only articular region of Meckel’s cartilage ossified. **n**, *Kentuckia hlavini*. Jaw laterally compressed. Only mentomeckelian and articular region of Meckel’s cartilage ossified. **o**, Actinopterygii n gen n sp (CMNH 9560). Jaw laterally compressed. Only articular region of Meckel’s cartilage ossified. p, *Tegeolepis clarki*. Only articular region of Meckel’s cartilage ossified. q, *Limnomis delaneyi.* Meckel’s cartilage unossified. r, *Gonatodus brainerdi*. Only mentomeckelian and articular region of Meckel’s cartilage ossified. s, *Cuneognathus gardineri*. Only mentomeckelian region of Meckel’s cartilage ossified. Not to scale. *Abbreviations*: cor, coronoid; co.bu, bump on coronoid; co.pa.te, coronoid-prearticular teeth; co.pa, coronoid-prearticular: co.pa.sh, coronoid-prearticular shelf; co.sh, coronoid shelf; co.te, coronoid teeth; de, dentary; de.sh, dentary shelf; de.te, dentary teeth; den, denticles; for, foramen; m.g, groove housing the mandibular canal; ma.te, marginal teeth; mc, mandibular canal; mec, Meckel’s cartilage; mec.bu, bumps on Meckel’s cartilage; mec.sh, shelf on Meckel’s cartilage; ment, mentomeckelian ossification; orn, ornament; t.pl, toothplate.

#### 4.1.1 General morphology of jaw

**– *Relative length of dentary****: (0) long (constitutes most of the length of the lower jaw); (1) short (constitutes less than half of jaw length) (Giles et al., 2023 character 79, and references therein).* The dentary generally forms most of the length of the lower jaw in actinopterygians (with the possible exception of *Meemannia*, although the absence of sutures between the dentary and any infradentary bones make this difficult to assess). It is short in at least a subset of sarcopterygians, most notably coelacanths and lungfishes, where the infradentaries contribute to a greater proportion of the mandible.
**– *Number of infradentaries****: (0) more than two; (1) two (angular and surangular); (2) one (angular only) (Giles et al., 2023 character 89, and references therein).* Absence of a surangular (supra-angular of Gardiner, 1984) was historically noted in only a small number of Devonian actinopterygians and the extant *Polypterus*, and inferred to be lost multiple times. Here, we report the presence of a surangular in *Gogosardina*, *Limnomis* and *Palaeoneiros*. Genuine absence appears to characterize *Mimipiscis*, “*Moythomasia*” *devonica*, and Actinopterygii n. gen. n. sp (CMNH 9560). The number of infradentaries cannot be determined in some taxa (e.g “*Kentuckia*” *hlavini*) because sutures are not apparent or are obscured by dermal ornament.
**– *Posterior infradentary reduced to thin strip in lateral view****: (0) absent; (1) present (new character).* In most Devonian actinopterygians, and some Carboniferous taxa (e.g. *Gonatodus punctatus*: Gardiner, 1967), the angular extends along the lateral surface of the mandible, with both anterior and dorsal arms forming the posteroventral region. In a subset of Devonian (e.g. *Palaeoneiros* and *Limnomis*) and some stratigraphically younger taxa (e.g. *Carboveles*: Moy Thomas, 1938), it is reduced to a thin strip surrounding the mandibular canal and barely visible in lateral view, with the dentary forming most of the lateral surface of the mandible. We caution that whether the angular occupies more than a ‘thin strip’ is subjective.
**– *Dentary with conspicuously reflexed distal tip****: (0) absent; (1) present (Giles et al., 2023 character 81, and references therein).* In profile, the mandible of most Devonian actinopterygians is deep posteriorly and tapers anteriorly, sometimes with a greater or lesser degree of dorsal inflection to the dorsal margin (e.g. “*Moythomasia*” *devonica*, Actinopterygii n. gen. n. sp. [CMNH 9560]). In some taxa, the anteriormost portion of the dentary is turned distinctly upwards into a reflexed distal tip. This morphology is present in the largest Devonian actinopterygian taxa included in this study: *Austelliscus*, *Tegeolepis* and *Howqualepis*. We note that a different morphology, where the dorsal margin of the mandible curves ventrally, is seen in “*Kentuckia*” *hlavini*; this shape is also seen in stratigraphically younger taxa such as *Beagiascus* (Mickle et al., 2009) and may be of future relevance as a phylogenetic character state.
**– *Concavity in anteroventral margin of dentary****: (0) absent; (1) present (Figueroa et al., 2021 character 266).* A slight concavity is present on the anteroventral margin of the dentary in *Austelliscus* and *Tegeolepis*.
**– *Coronoid process of lower jaw****: (0) absent; (1) present (Giles et al., 2023 character 93, and references therein).* In some actinopterygians, the posterior portion of the mandible is raised into a distinct, convex crest, commonly referred to as a ‘coronoid process’. Historically, the presence of this process has been considered a neopterygian synapomorphy (Gardiner, 1984, Gardiner & Schaeffer, 1989). However, raised processes in structurally homologous positions—although often contributed to by different ossifications—are present in a range of non-neopterygian taxa (see discussion in Coates & Tietjen, 2018). Given the range of possible configurations, we keep the broad term ‘coronoid process’ to refer to the development of a dorsal crest on the mandible and consider the specific ossifications contributing to such a process in a separate character. Here, we recognize the presence of coronoid processes in *Howqualepis*, *Raynerius*, *Palaeoneiros*, *Osorioichthys*, *Limnomis*, and “*Gonatodus*” *brainerdi*.
**– Coronoid process contributed to by**: (0) prearticular only; (1) surangular only; (2) dentary plus postdentary bones; (3) angular only (Giles et al., 2023 character 94, and references therein). The broadly defined ‘coronoid process’ can be composed by different sets of bones. The most common condition amongst Devonian (e.g. Raynerius; Palaeoneiros) and Carboniferous (e.g. Trawdenia: Coates & Tietjen, 2018; Mecca Quarry Shale ‘elonichthyid’: Gottfried, 1992) appears to be a coronoid process formed exclusively by the surangular. In some taxa (e.g. Osorioichthys; Limnomis, Aesopicthys: Poplin & Lund, 2000), the dentary may contribute alongside one or more postdentary bones; this condition is also common in neopterygians. A coronoid process formed exclusively by the prearticular is a synapomorphy of polypterids and scanilepiforms (Claeson et al., 2007; Giles et al., 2017).
– **Jaw margins overlain by lateral lamina**: (0) absent; (1) present (Giles et al., 2023 character 85, and references therein). The maxillary dentition is obscured in lateral view by a thin lamina of bone in Styracopterus, Fouldenia and Amphicentrum (Sallan & Coates, 2013).

#### 4.1.2. Mandibular canal

**– Mandibular canal**: (0) primarily carried by infradentaries; (1) primarily carried by dentary (Giles et al., 2023 character 78, and references therein). The mandibular canal is transmitted largely within the dentary in actinopterygians (with the possible exception of Meemannia, although the absence of sutures between the dentary and infradentaries make this difficult to assess), and within the infradentaries in sarcopterygians.
**– Course of mandibular canal**: (0) traces ventral margin of jaw along entire length; (1) arches dorsally in anterior half of jaw (Giles et al., 2023 character 76, and references therein). In the Devonian actinopterygians Cheirolepis, Austelliscus, Howqualepis, Gogosardina, Mimipiscis, Moythomasia and Raynerius (as well as some stratigraphically younger taxa: e.g. Australosomus: Nielsen, 1949), the mandibular canal is confined to the ventral margin of the mandible along the posterior half to two thirds of the mandible and arches dorsally in the anterior third. In some taxa (e.g. Osorioichthys), the canal is oriented anterodorsally along much of its length; this may be of future relevance as a phylogenetic character.
**– Mandibular canal reaches anterior margin of mandible**: (0) present; (1) absent (Giles et al., 2023 character 77, and references therein). Most commonly, the mandibular canal reaches the anterior extremity of the dentary. In some taxa, however (e.g. Mimipiscis toombsi), the canal opens into the dorsal margin some way posterior to the anterior margin.

#### 4.1.3. Tooth characters

**– Teeth on dentary**: (0) present; (1) absent (Giles et al., 2023 character 80, and references therein).
**– Teeth of outer dental arcade**: (0) several rows of disorganized teeth; (1) two rows, with large teeth lingually and small teeth labially; (2) single row of teeth (Giles et al., 2023 character 84, and references therein). Teeth on the dentary of early actinopterygians are most commonly arranged into a lingual row of larger teeth and a labial row of smaller, marginal teeth (Fig. 20; e.g. Gogosardina, Mimipiscis toombsi). However, the teeth appear to be more irregular in Cheirolepis canadensis (Arratia & Cloutier, 1996) and Cheirolepis jonesi, with several poorly organized rows on the mandible. Marginal dentition is absent (or reduced to the point of functional absence) in a handful of Devonian taxa (e.g. Meemannia, Osorioichthys, Palaeoneiros, the unnamed Cleveland Shale form); this morphology is also seen in some early sarcopterygians (e.g. Psarolepis; Yu, 1998) and stratigraphically younger actinopterygians (e.g. Aesopichthys: Poplin & Lund, 2000).
**– Base of dentary teeth**: (0) level with dorsal rim of dentary; (1) recessed ventrally (new character). The position of the base of the teeth of the main dentary tooth row is variable among early actinopterygians (Fig. 20). They may be level with the dorsal margin of the dentary, as in Mimipiscis bartrami, “Moythomosia” devonica, Osorioichthys, “Kentuckia” hlavini and “Gonatodus” brainerdi. In other taxa, however (e.g. Austelliscus, Tegeolepis, Raynerius, Howqualepis), the base of the tooth is positioned more ventrally, such that the bottom portion of the tooth is obscured by the dentary when viewed laterally.
**– Enlarged series of teeth on anterior (parasymphysial) region of dentary**: (0) absent; (1) present (modified from Giles et al., 2023 character 82, and references therein). The teeth in the anteriormost region of the dentary may be enlarged relative to those seen elsewhere on the dentary. This is most noticeable in Austelliscus and Tegeolepis.
**– Procumbent parasymphysial teeth**: (0) absent; (1) present (new character). In most taxa, the teeth borne on the dentary terminate prior to its anterior margin. In taxa such as Meemannia and Tegeolepis, however, a number of procumbent teeth are present on the anteriormost symphysial margin of the dentary.
**– Parasymphysial tooth whorl (or articulation facet for whorl on anterior dentary)**: (0) present; (1) absent (modified from Giles et al., 2023 character 83, and references therein). In early sarcopterygians, a parasymphysial tooth whorl is present at the anterior extent of the mandible, attaching to its medial surface via a distinct facet. Given the disarticulated nature of many early osteichthyan fossils, the attachment area for the whorl is known in more taxa than the whorl itself (Zhu et al. 1999, Zhu et al. 2009; these taxa are sometimes interpreted as stem osteichthyans), although articulated parasymphysial tooth whorls are known in onychodonts (Andrews et al., 2006) and porolepiforms (Jarvik 1972). The homologies of this whorl to other parts of the coronoid dentition have been debated at length (e.g. Zhu & Yu 2004; Friedman & Blom 2006). Here, we distinguish between a parasympyhsial tooth whorl and other elements of the inner dental series (which may or may not bear whorled teeth) based on topology and anatomy. We consider a parasymphysial tooth whorl to be a distinct element at the anterior extent of the mandible, discontinuous with the rest of the coronoid series and separated from it by a gap, housed in its own facet, and with a continuous and distinctly curved base. Parasymphysial tooth whorls appear to be primitive for sarcopterygians but have not been observed in any actinopterygians (see below for discussion of anterior coronoids in Cheirolepis and Howqualepis). The anterior portion of the mandible in Meemannia bears a deep concavity (Fig. 2), but this is oriented medially and appears to represent the space occupied in life by the unossified mentomeckelian portion of Meckel’s cartilage.
**– Acrodin caps on teeth**: (0) absent; (1) present (Giles et al., 2023 character 86 and references therein). In many actinopterygians, hypermineralised tissue is restricted to a narrow cone at the apex of the teeth, bounded by a morphologically distinct collar region. This cap is referred to as ‘acrodin’. We express caution in interpreting the absence of acrodin from tomograms: for example, acrodin has been described in thin section on the teeth of Mimipiscis (Gardiner, 1984), but is not always easily observable in our CT data, despite the relatively high resolution. However, acrodin can sometimes be seen in tomograms, for example in Gogosardina, and has also been reported previously in CT-based studies (Friedman et al., 2024).
**– Plicidentine**: (0) absent; (1) present (Giles et al., 2023 character 87, and references therein). Complex infolding of the dentine around the base of teeth, referred to as plicidentine, is widespread in early sarcopterygians (Jarvik 1980; Zhu & Yu 2004). It is more sporadically distributed among early actinopterygians, being reported in Cheirolepis canadensis (Meunier et al., 2018) and some Carboniferous large-toothed forms (e.g. Friedman et al., 2024).

#### 4.1.4. Inner dental series

**– Coronoids (sensu stricto, excluding parasymphysial tooth whorl)**: (0) present; (1) absent (Giles et al., 2023 character 90, and references therein). The presence of coronoids can now be confirmed in taxa such as *Osorioichthys* and *Limnomis*, in which the medial surface was previously unknown. As discussed above, we exclude the parasymphysial tooth whorl from this series.
***– Number of coronoids****: (0) five or more; (1) four; (2) three; (3) two; (4) one (Giles et al. 2022 character 91, and references therein).* A higher number of coronoids appears to be plesiomorphic for osteichthyans (Zhu & Yu, 2004; Friedman, 2007). We confirm that the number of individual coronoids is variable in early actinopterygians, ranging from three in *Cuneognathus* to nine in *Cheirolepis trailli*. We note that the coronoids (and prearticular) are commonly fused into a single series, and this character cannot be coded in those taxa.
**– *Enlarged series of teeth on anterior coronoid, excluding parasymphysial tooth whorl****: (0) absent; (1) present (new character).* As the anterior coronoid in the taxa studied here is continuous with the remainder of the coronoid series and has a flat base that articulates with the dentary in the same way as the more posterior coronoids, we consider it part of the coronoid series rather than a separate parasymphysial element. The anterior coronoid of several Devonian actinopterygians bears enlarged teeth relative to those seen on more posterior portions of the inner dental series. The form of this varies from two to three teeth resembling miniature fangs (e.g. *Gogosardina*, *Moythomasia durgaringa*) to a longer row of larger teeth (e.g. *Cheirolepis trailli*) to a single posteriorly directed cusp (e.g. “*Gonatodus*” *brainerdi*). This variance may be of future relevance as a phylogenetic character.
**– *Whorl-like teeth on anterior coronoid, excluding parasymphysial tooth whorl****: (0) absent; (1) present (new character).* The anterior coronoid of some taxa (*Cheirolepis jonesi*, *Howqualepis*) bears a series of teeth that are strongly recurved and aligned in a row such that each partially overlaps the tooth posterior to it. As this anterior coronoid is continuous with the remainder of the coronoid series and has a flat base that articulates with the dentary in the same way as the more posterior coronoids, we consider it as part of the coronoid series rather than a separate parasymphysial element.
**– *Broad anterior coronoid, excluding parasymphysial tooth whorl****: (0) absent; (1) present (new character).* In most actinopterygians, the anteriormost coronoid is the same width as those positioned more posteriorly in the series. In contrast, in *Cheirolepis jonesi*, *Howqualepis*, *Osorioichthys*, and Actinopterygii n. gen. n. sp. (CMNH 9560), the anterior coronoid is expanded medially, forming a broad, flat surface. An additional variation on this, where the ventromedial surface of the anterior coronoid twists to form a dorsomedial surface, is seen in “*Gonotadus*” *brainerdi*; if observed in other taxa, this unusual arrangement may be a useful as a separate character.
**– *Position of coronoids****: (0) restricted above dorsal shelf of mandible; (1) ventral extension down medial surface of mandible (new character).* In most actinopterygians (e.g. *Gogosardina*, *Mimipiscis*), the coronoid elements extend ventrally, to a greater or lesser extent, down the medial surface of the mandible (Fig. 20). However, in *Cheirolepis* and *Tegeolepis*, the coronoid series is dorsally restricted, sitting entirely on top of the medial shelf of the dentary, with no ventral extension. Although coronoids are not preserved in *Meemannia*, a similar morphology—with the series sitting on top of the Meckelian shelf—is inferred here and in the original description (Zhu et al., 2010). This dorsal restriction of the coronoids, with a more ventrally extensive prearticular, characterizes many sarcopterygians (Zhu & Yu, 2004).
**– *Coronoid series divided into dorsal and ventral regions****: (0) absent; (1) present (new character).* In *Gogosardina*, *Mimipiscis toombsi*, *M. bartrami*, *Moythomasia durgaringa* and *Raynerius*, the coronoids are divided into two distinct tooth-bearing surfaces: a dorsal region, oriented dorsally; and a ventral region, oriented medially, that does not extend to the anteriormost margin of the mandible. Other variation seen in coronoid anatomy, such as a ventral region that extends further anteriorly in *Limnomis*, may prove to be of future phylogenetic relevance if identified in other taxa.
**– *High coronoid-prearticular shelf****: (0) absent; (1) present (new character)*. In some Devonian actinopterygians examined here, the coronoid-prearticular series forms a high shelf that extends almost as high as (e.g. *Moythomasia durgaringa*, *Palaeoneiros*), or in some cases higher than (e.g. “*Moythomasia*” *devonica*, Actinopterygii n. gen. n. sp. (CMNH 9560), “*Gonatodus*” *brainerdi*) the tips of the dentary teeth. There is also variation in the height, width, and tooth-bearing nature of this shelf, which may carry additional signal.
**– *Prearticular****: (0) absent; (1) present (Giles et al., 2023 character 106, and references therein)*.
**– *Posterior prearticular plate****: (0) absent; (1) present (new character).* A second, small prearticular plate, lying posterior to the primary prearticular, is present in *Moythomasia durgaringa* (Gardiner 1984) and *Raynerius*.
**– *Palatal bite****: (0) absent; (1) present (Giles et al., 2023 character 98, and references therein).* Teeth on the dentary represent the primary means by which early actinopterygians interact with food items and bite. However, some Carboniferous (e.g. *Notacmon [=Eurynotus]*, Friedman et al., 2019) and stratigraphically younger taxa process food by means of toothplates or other surfaces developed on the palate and corresponding inner surface of the mandible.
**– *Medial mandibular toothplates****: (0) absent; (1) present (new character).* A series of three rhomboid toothplates, which collectively form an ovoid shape, lie medial to the inner surface of the mandible in *Palaeoneiros* (Giles et al., 2022). These have previously only been identified in *Pteronisculus stensiö* (Nielsen, 1942), although we caution that their presence may have been overlooked in previous studies and it may only be possible to confidently assess their presence or absence in taxa described via serial sectioning or X-ray CT imaging of articulated specimens.

#### 4.1.5. Meckel’s cartilage

***– Ossification of mentomeckelian region:*** *(0) present; (1) absent (Giles et al., 2023 character 88, and references therein).* Mineralisation of Meckel’s cartilage is highly variable in Devonian actinopterygians. It is most commonly ossified along the entire jaw as a continuous element (e.g. *Meemannia*, *Mimipiscis*, *Raynerius*, *?Moythomasia devonica*), or the articular and mentomeckelian may be ossified independently (e.g. *Osorioichthys*, “*Kentuckia*” *hlavini*, “*Gonatodus*” *brainerdi*). The articular alone is ossified in a handful of taxa (e.g. *Cheirolepis trailli*, *Howqualepis*, *Tegeolepis*, undescribed Cleveland Shale taxon), and Meckel’s is completely unossified in *Cheirolepis jonesi* and *Limnomis*.
***– Retroarticular process****: (0) present; (1) absent (Giles et al., 2023 character 97, and references therein).* In most early actinopterygians, the posterior margin of the dermal bones of the mandible and the posterior margin of the endoskeletal elements of the mandible are approximately level, with the articular forming a vertical surface ventral to the articular cotyles. However, in *Osorioichthys* and “*Gonatodus*” *brainerdi*, the endoskeletal element extends posterior to both the articular cotylar surface and the dermal elements, forming a modest retroarticular process. It is not clear whether any ligaments or bones of the hyoid arch articulate with this surface, as is the case for the (non-homologous) retroarticular processes of some sarcopterygians (e.g. coelacanths: Friedman, 2007).
**– Raised articular region**: (0) absent; (1) present (new character). In most taxa, the articular region of Meckel’s cartilage accommodating the adductor fossa and articular cotyles lies level with the rest of the upper margin of the mandible. In *Moythomasia durgaringa*, the articular region is elevated dorsally relative to the rest of the mandible, above the tips of the dentary teeth. This morphology is also seen in some Carboniferous durophagous (Friedman et al., 2019) and large-toothed ‘macrodont’ (Friedman et al., 2024) actinopterygians.

### 4.2 Patterns of jaw evolution in early actinopterygians

The lack of a robust phylogenetic framework with which to examine the distribution and optimization of the above characters makes it difficult to assess their significance for relationships: indeed, it is hoped that they will feed into more stable phylogenetic hypotheses. Nevertheless, comparison with sarcopterygians and stratigraphically younger actinopterygians with affinities to extant radiations allow some preliminary conclusions to be drawn.

In the earliest diverging actinopterygian (Lu et al., 2016) *Meemannia*, the mandible resembles that of many early sarcopterygians, with a blunt and rounded profile, infradentary foramina, and abundant pore openings matching its original description as a lobe finned fish (Zhu et al., 2010). Although not preserved, the prearticular appears to be positioned ventrally and the coronoids dorsally, the latter sitting atop Meckel’s cartilage. A similar dorsal position for the coronoids, albeit atop a medial shelf of the dentary rather than directly on Meckel’s cartilage is also seen in *Cheirolepis* and *Tegeolepis*, and may also have been the condition in *Austelliscus*. In these taxa, the coronoid series lacks the ventral extension down the medial face of the dentary seen in almost all other early actinopterygians (Fig. 20). As previously noted by Friedman and Blom (2006), the mentomeckelian region of Meckel’s cartilage is unossified in *Tegeolepis* and *Howqualepis*; we can additionally confirm its absence in *Cheirolepis*. Other mandibular characters are also shared across some or all of these taxa, including a reflexed anterior dentary region with teeth that are enlarged relative to the rest of the tooth row (*Tegeolepis*, *Howqualepis*, *Austelliscus*), a notch on the anteroventral region of the dentary (*Tegeolepis*, *Austelliscus*), and whorl-like teeth on the anterior coronoid (*Howqualepis*, *Cheirolepis jonesi*). Recumbent teeth also extend onto the anteriormost symphysial region in *Meemannia* and *Tegeolepis*.

Several previous studies have suggested a close relationship between some or all of these taxa near the base of the actinopterygian tree. A clade comprising *Howqualepis* and *Tegeolepis* was first proposed by Friedman & Blom (2006), supported by features of the pectoral fin and mandible, as well as the large size and elongate rostrum of these taxa. Long et al. (2008) resolved an alternative clade comprising *Donnrosenia* and *Howqualepis* to the exclusion of *Tegeolepis*, but subsequent studies have generally either recovered a closer relationship between *Howqualepis* and *Tegeolepis* (Swartz, 2009; sometimes including *Donnrosenia* within this clade: Choo, 2012) or failed to resolve the relationships between these and other taxa (e.g. Giles et al., 2015b). Surprisingly, the mandible of *Osorioichthys*, which is often recovered as sister taxon to almost all other actinopterygians, sometimes in association with *Tegeolepis*, shows a strikingly different mandibular morphology. This taxon has a blunt dentary lacking enlarged teeth and marginal dentition, a completely ossified Meckel’s cartilage, a coronoid process formed by the dentary and infradentary bones, and a broad prearticular-coronoid series that forms a continuous field of dentition together with the dentary. At present, stratigraphically younger taxon with close affinities to *Osorioichthys* are difficult to identify, but we also note the presence of a suborbital, representing an unusual feature for a Devonian actinopterygian (Giles et al., 2022).

A posterior prearticular plate is uniquely shared between *Moythomasia durgaringa* and *Raynerius splendens*. General similarities between these taxa were recognized when the latter was erected (Giles et al., 2015), and a sister group relationship has been recovered between *Raynerius* and moythomasiids in some subsequent analyses (e.g. Giles et al., 2017; Caron et al., 2023). We note that the mandible of “*Moythomasia*” *devonica* displays striking differences to that of *Moythomasia durgaringa*: the former has a slender jaw that tapers anteriorly, a low articular region, a high and expanded coronoid-prearticular plate, and a narrow angular, more closely resembling stratigraphically younger actinopterygians loosely referred to as ‘rhadinichthyids’. In agreement with Choo (2015), we regard its inclusion in *Moythomasia* as in need of review.

One hallmark of more derived actinopterygian fishes, the coronoid process, is present among Devonian taxa, although its connection to that process in living taxa is unclear. In neopterygians (and, derived independently, polypteriforms), the coronoid process increases the surface area of the lateral wall of the adductor fossa, creating a larger attachment area for the adductor mandibular musculature, and raises the posterior margin of the mandible, essentially turning the jaw into a bent lever arm. Together, these increase torque and facilitate a stronger bite (Schaeffer & Rosen, 1961; Lauder, 1980).

Previous studies have unearthed hints of a coronoid process in Carboniferous fishes, formed either from a dorsal extension of the surangular (Mecca Quarry Shale ‘elonichthyid’: Gottfried, 1992; *Coccocephalichthys wildi*: Poplin & Veran, 1996; *Trawdenia planti*: Coates & Tietjen, 2018, in which it is referred to as a ‘surangular process’); a narrow, high extension of the dentary and angular (*Aesopichthys*: Lund & Poplin, 2000); a broad, low extension of the dentary and angular (*Paramblypterus duvernoyi*: Dietze, 2001); or the prearticular (*Notacmon* [=*Eurynotus*] *crenatus*: Friedman et al., 2019). We identify two types of incipient coronoid processes in Devonian taxa: a process on the dorsal margin formed only by the surangular (present in *Howqualepis*, *Raynerius*, *Palaeoneiros*, and “*Gonatodus*” *brainerdi*); and an overall dorsal expansion of the mandible formed by the dentary and postdentary ossifications (present in *Osoriochthys* and *Limnomis*). The evolution of a coronoid process is frequently treated as part of a linear sequence, representing outdated concepts of “levels of organization” (Schaeffer & Rosen, 1961). Given its clear functional significance, and the origin of mechanically comparable structures in many other vertebrate groups, it seems likely that coronoid processes evolved multiple times independently beyond the examples in extant polypterids and neopterygians.

We note that the mandibles of “*Gonatodus*” *brainerdi* and “*Kentuckia*” *hlavini* bear little similarity to the Carboniferous type species of these genera, lending further support for the argument that these genera are both polyphyletic (Traquair, 1907; Dunkle, 1964; Gardiner, 1967; Giles & Friedman, 2014). Given the amenability of this material for CT-based investigation, they both represent priorities for future taxonomic revision.

### 4.3 Functional considerations

Devonian actinopterygians are often perceived as functionally or ecologically homogenous, especially in comparison to coeval sarcopterygians and placoderms or stratigraphically younger ray-fins. Quantitative analyses of functional traits relating to jaws and teeth (Anderson et al., 2011) as well as overall mandibular shape (Hill et al., 2018) add support to this conventional view. While we agree that the overall variation in gross jaw structure—and likely function—is lower in Devonian actinopterygians than other groups, our study exposes previously unrecognized variation within the earliest ray-finned fishes. These new anatomical details complement past evidence for trophic diversity among Devonian actinopterygians. Direct support for diets comes in the form of gut contents for three Late Devonian species that were either examined directly or assigned to genera for which we studied other species: *Cheirolepis canadensis* (specimens including the acanthodian *Homalacanthus* or smaller individuals of *Cheirolepis*; Chevrenais et al., 2017), *Gogosardina coatesi* (one individual containing remains of the conodonts *Oulodus* and *Icriodus*; Nicoll, 1977; Choo et al., 2009), and *Tegeolepis clarki* (one specimen containing parts of a small arthrodire similar to *Selenosteus* plus a chondrichthyan fin spine; Williams, 1990). Although these three taxa are associated with distinct environments (estuarine: *Cheirolepis*; reef: *Gogoardina*; offshore, but potentially shallow, marine: *Tegeolepis*; Long and Trinajstic, 2010; Cloutier et al., 2011; Alsharani and Evans, 2014), in each case prey items represent nektonic vertebrates. The size of these prey items varies, pointing to a major piece of circumstantial evidence for dietary diversity among Devonian actinopterygians: predator size. The taxa surveyed in this contribution span an order of magnitude in jaw length (Fig. 21), ranging from *Limnomis* (jaw length = 8.2 mm) to *Tegeolepis* (jaw length = 173 mm). This substantial variation implies differences in diet since, unsurprisingly, larger fishes with larger gapes can consume larger prey (Scharf et al., 2000). Formal analyses have also suggested some dietary variation stemming from differences in the dentition of Devonian actinopterygians (Gauchey et al., 2014).

**Fig. 21.**
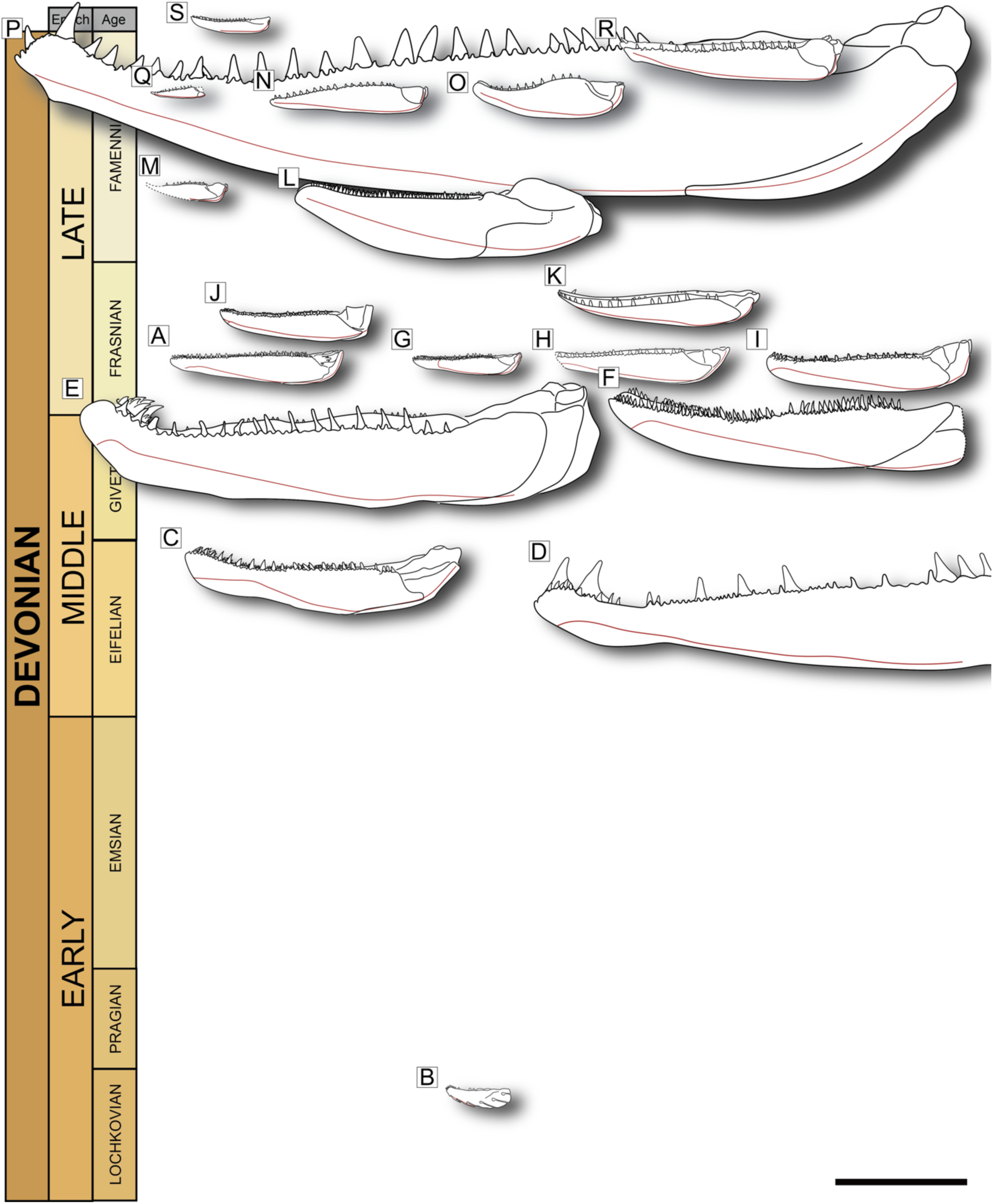
Mandibles of Devonian taxa included in this study, arranged stratigraphically. **a**, Gogosardina coatesi. **b**, Meemannia eos. **c**, Cheirolepis trailli. **d**, Austelliscus ferox. **e**, *Howqualepis rostridens*. **f**, Cheirolepis jonesi. **g**, Mimipiscis toombsi. **h**, Mimipiscis bartrami. **i**, Moythomasia durgaringa. **j**, Raynerius splendens. **k**, ?*Moythomasia devonica*. **l**, Osorioichthys marginis. **m**, Palaeoneiros clackorum. **n**, *Kentuckia hlavini*. **o**, Actinopterygii n. gen. n. sp. (CMNH 9560). **p**, Tegeolepis clarki. **q**, Limnomis delaneyi. **r**, Gonatodus brainerdi. **s**, Cuneognathus gardineri. Red line indicates path of mandibular canal. Scale bar = 20 mm.

Here we examine the possible functional consequences of the differences we observed between early ray-fin mandibles, recognizing that the lower jaw represents just one component of the complex actinopterygian feeding mechanism. However, and in contrast to extant ray-fins that show substantial variation in mobility of upper jaw bones, early forms are relatively homogeneous in sharing a premaxilla and maxilla bound to other parts of the dermal skull and palate (Schaeffer & Rosen, 1961; Lauder, 1980). This does not mean that non-neopterygian actinopterygians had no cranial kinesis (see Pearson & Westoll, 1979 for *Cheirolepis* and Whitlow et al., 2022 for *Polypterus*), but rather that the overall range in patterns of skull mobility was substantially narrower than in extant taxa (Westneat, 2005). Consequently, modifications to the lower jaw and its dentition might represent the key path for trophic diversification among the earliest ray-fins. These observations are largely qualitative and informed by the rich body of literature on fish ecomorphology and functional anatomy. Quantitative analysis is beyond the scope of the present study but will inform future comparisons between Devonian and Carboniferous actinoptergyians (cf. Sallan & Friedman, 2012).

#### 4.3.1 Functional consequences of differences in mandibular structure

Most work on the function of actinopterygian lower jaws has anticipated a generalised biting morphotype for all Devonian taxa (Schaeffer & Rosen, 1961; Pearson & Westoll, 1979; Lauder,1980; Lauder, 1981; Gottfried, 1992), and this feeding mode does indeed appear the most common in Devonian actinopterygians. Extant fish that employ biting tend to have a longer gape than those that do not (Markey et al., 2006), and Devonian actinopterygians typically had a proportionally large gape. Variations in overall jaw shape impact jaw mechanics (e.g., jaw-opening and -closing lever arms; Westneat, 2004), the physical properties of the mandible (e.g., stiffness as governed by second moment of area; Huie et al., 2022), and patterns of dental occlusion (e.g., simultaneous versus sequential occlusion as a function of position of the jaw joint; Ramsay & Wilga, 2007). While most Devonian actinopterygians have broadly similar jaw shapes, there are some notable differences with probable functional consequences. Particularly deep jaws characterize *Meemannia*, *Osorioichthys*, and the unnamed Cleveland Shale taxon. This can confer additional stiffness in dorsoventral compression, permitting more forceful bites.

*Osorioichthys* combines a stout mandible bearing the most substantial coronoid process found among Devonian actinopterygians, further distinguishing the taxon from its contemporaries.

Variation is also seen in the posterior region of the lower jaw. Many species have jaw joints that are effectively in line with the dentary teeth. However, *Moythomasia*, *Raynerius*, *Howqualepis*, and, to a lesser degree, *Cheirolepis trailli*, have articulars that are raised above the dorsal tips of the primary dentition (Figs 9, 21i). This arrangement impacts occlusion, with the entire toothed length of the mandible brought into contact with the upper jaw—and thus the prey item—at the same time (Ramsay & Wilga, 2007).

Comparable displacements of the jaw joint are found in many post-Devonian actinopterygians, including multiple macrodont taxa from the Carboniferous and Permian (summarized by Friedman et al., 2024). *Osorioichthys* also shows articular offset suggesting similar functional consequences, but here the jaw joint lies ventral to the dentary tooth row. This arrangement is unusual among Paleozoic actinopterygians, but is common among early sarcopterygians including lungfishes (Miles, 1977) and coelacanths (Forey, 1997), as well as many extant ray-fins (Gregory, 1933).

#### 4.3.2. Functional consequences of differences in dentition

Differences in teeth, in terms of their shape, size, and distribution, point to contrasts in feeding among Devonian actinopterygians. In modern reef fishes, regardless of phylogenetic affinities, taxa with similar dentition have similar predation strategies and diets (Mihalitsis & Bellwood, 2019a, b; 2021; Muruga et al., 2022). However, extant fish with superficially similar morphologies can still exhibit somewhat different feeding behaviours (Porter & Motta, 2004). Work on extant fishes shows that denticles are more suited to gripping prey, whereas a sharp toothrow is more suited to damage and post-capture processing of large prey (Muruga et al. 2022). Past quantitative work suggests diversification in feeding strategies during the Late Devonian based on variation in the shape of isolated actinopterygian teeth (Gauchey et al., 2014). However, different tooth shapes often occur in single individuals, and the distribution of tooth forms across the jaws can have important functional implications (Mihalitsis & Bellwood, 2019a, b; 2021). The taxa we surveyed show noteworthy differences in tooth size and shape, as well as differences in the degree to which dentition appears regionalized along the mandible. Although most variation pertains to dentary teeth, there are also clear differences in inner dentition and the coronoid bones themselves with likely functional contrasts.

One major contrast relates to the arrangement of the teeth themselves. Most taxa bear a single row of principal dentary teeth that is often flanked by a smaller lateral row and supplemented mesially by small teeth or denticles on the coronoids and prearticular. The principal dentary teeth, which can be quite large in some taxa, suggest a role in piercing and retaining prey items. A contrasting feeding morphology appears to be present in the Middle Devonian *Cheirolepis*, which displays several features in common with the villiform functional morphotype of extant teleosts (Mihalitsis & Bellwood, 2019). *Cheirolepis* bears numerous small-medium teeth across both its inner and outer dental series, with a medially enlarged anterior coronoid contributing to a broad field of teeth. Living villiform taxa utilise their teeth for post-capture gripping of prey, with minimal damage to the prey item. This feeding modality in *Cheirolepis* is supported by essentially intact acanthodians found as gut contents in some specimens of *C. canadensis* (e.g., Chevrinais et al., 2017: fig. 1).

Among other taxa, there are significant differences in the relative size, distribution, and orientation of the largest mandibular teeth. *Austelliscus* and *Tegeolepis* are noteworthy in having extremely large, widely spaced, principal teeth and a reflexed anterior dentary tip bearing larger fangs. Both taxa also show clear differences in tooth-crown orientation along the jaw, with anterior teeth reclined posterodorsally and those at the rear of the jaw inclined anterodorsally. A similar gradient of tooth angles is apparent in *Howqualepis*. *Austelliscus* and *Tegeolepis*, the largest Devonian actinopterygians known to date, and their collection of traits resembles that of the macrodont functional morphotype in extant fishes (Mihalitsis & Bellwood, 2019). Similarities are strongest with front-fanged macrodonts, which possess a row of large, widely spaced teeth and anterior fangs. Extant macrodont teleosts are typically piscivorous grabbers (Mihalitsis & Bellwood, 2021). Although their dentaries are morphologically similar to these Devonian taxa, most use enhanced suction feeding via highly kinetic upper jaws to engulf prey before employing their large teeth for post-capture processing (Mihalitsis & Bellwood, 2021). The mandibles of *Austelliscus* and *Tegeolepis* also recall those of extant shortnose gar (Meunier & Brito, 2017) in possessing a similarly shaped toothrow with enlarged teeth anteriorly and in the symphysial region. Gars are typically piscivorous predators that lack a mobile upper jaw, employing ram-feeding (rather than significant suction feeding) to ambush prey followed by grabbing with their teeth (Kammerer et al., 2006). Sideways head movements can also be used to bring prey in between the jaws during capture (Porter & Motta, 2004), and the well-developed marginal dentition in *Tegeolepis* and *Austelliscus* may have contributed to a similar function.

Several Devonian actinopterygians bear one or more enlarged teeth on the anterior coronoid that may resemble miniature fangs (e.g. *Cheirolepis trailli*, *Gogosardina coatesi*); these can be arranged into a whorl-like structure (*Cheirolepis jonesi*, *Howqualepis rostridens*) or even point posteriorly (e.g., “*Moythomasia*” *devonica*). Enlarged anterior coronoid teeth are reminiscent of the enlarged anterior dentary fangs of *Tegeolepis* and *Austelliscus*, although they are clearly non-homologous and often do not occur in conjunction with a reflexed tip of the jaw as they do in those two genera plus *Howqualepis*. However, it is possible these different kinds of expanded parasymphysial dentitions served a similar role in prey capture, acting principally to secure prey via grabbing and subsequent puncture (Mihalitsis & Bellwood, 2019). Like extant analogues, they likely employed ram-feeding, an effective strategy in the capture of elusive prey (Norton, 1991). Parasymphysial fangs, borne either by the dentary or on a tooth whorl in series with the coronoids, are widely distributed among many of the sarcopterygian contemporaries of the Devonian ray-fins considered here (Janvier, 1996).

More posterior coronoids and the prearticular generally show smaller teeth and denticles, although the arrangement and size of these varies considerably among taxa (Fig. 20). The prearticular dentition of *Howqualepis* is especially distinctive, bearing a row of enlarged, posteriorly curved teeth near its dorsal margin. While teeth of this size are generally directed dorsally such that their apices contact the prey during jaw closing, these enlarged prearticular teeth are strongly deflected to the horizontal plane, directing their tips toward the centre of the buccal cavity. This suggests a role in restraining prey once it was in oral chamber rather than initial capture. Horizontally directed teeth, presumably with a similar function, are borne by bones of the palate in some Paleozoic actinopterygians (e.g., *Daemodontiscus*; Friedman et al., 2024) and extant teleosts (e.g., *Esox*; Brocklehurst et al., 2019).

A final, and unanticipated, observation relates to relative position of the coronoid series and its consequences for the bones and teeth making major contributions to the bite. Typically, the dentary teeth of the mandible contact the food item and/or occlude with the upper jaw first. However, in several Devonian taxa the inner dental series—that is, the coronoids and/or prearticular—extends far enough dorsally that it lies level with, or sometimes even above, the tips of the dentary teeth (Fig. 20). As such, the inner dental series interacts with food items in conjunction with, or before, the dentary teeth. This trait is developed in several different ways in Devonian taxa. The coronoid-prearticular series may form a narrow plate that bears one or multiple rows of denticles on its medial surface, as in “*Moythomosia*” *devonica*, *Palaeoneiros*, *Gonatodus*, *Limnomis* and *Cuneognathus*; in these taxa, the dentary retains sizeable teeth, although the height of the shelf varies. In other taxa, such as *Osorioichthys* and the unnamed Cleveland Shale taxon, the coronoid-prearticular is also mediolaterally expanded to form a horizontal platform with denticles on its dorsal surface. The functional significance of this is a broad, flat inner dental series that interacts with food items in conjunction with, or before, the dentary teeth. At its extreme, the outer dentition would appear to be excluded from food processing. This shift from the outer to inner dental series is seen in multiple groups with palatal bites, an arrangement that is often thought to be associated with the processing of harder prey items (e.g., Cui et al., 2022). Among actinopterygians, palatal bites are well-developed in the Permo-Carboniferous eurynotiforms (Friedman et al., 2019; Bradley Dyne, 1939). Many of these taxa bear dental plates that appear to form from fusion of individual bulbous tooth cusps on the inner surface of the mandible and palate (Sallan & Coates, 2012; Elliott & Giles, 2024). The oldest eurynotiforms are early Carboniferous (Tournaisian) in age, and even the most generalised taxa like *Fouldenia* and *Styracopterus* already bear conspicuous specialisations (Sallan & Coates, 2012). We have no evidence of a relationship between the taxa in this study and these Carboniferous durophages, but Devonian actinopterygians with this enlarged coronoid shelf may provide a model of how this distinctive feeding strategy arose.

## 5. CONCLUSIONS

We provide a comprehensive examination of the lower jaw of Devonian actinopterygians, including detailed descriptions of the mandible for roughly two thirds of all known species. Consistent with previous commentary, we recognize overall similarity in gross shape and composition among the mandibles of the earliest ray-finned fishes. The lower jaw typically has a high aspect ratio, is deep posteriorly and tapers anteriorly. It is contributed to laterally by the canal-bearing dentary with one or two smaller infradentaries, internally by the variably ossified Meckelian element(s), and mesially by the prearticular and numerous coronoids. Despite this broadly conserved architecture, we find new morphological details and conspicuous differences between taxa that are not apparent from external examination of specimens or in previous descriptions. These include updates to the structure of the inner and outer dermal dental series, the ossification and position of Meckel’s cartilage, and the form and arrangement of the dentition.

We anticipate these new data will have implications for two major gaps in understanding of early actinopterygian evolution. First, they will expand phylogenetic character sets developed for inferring relationships among Palaeozoic actinopterygians. Existing phylogenetic hypotheses are generally unstable and often contain only a minor fraction of Devonian ray-fin diversity, partly due to limited anatomical data and a general perception that actinopterygians of this age are structurally homogenous. We review and revise existing mandibular characters, with the intention that these will be incorporated into future phylogenetic analyses. Our observations lead to 12 novel characters relating to general jaw form, tooth morphology, the inner dental series and Meckel’s cartilage. These, along with pre-existing and revised characters, contribute to a total of 44 mandibular characters for Devonian actinopterygians. In addition to providing more characters, we anticipate the greater detail available for some poorly known species will also permit expanded taxonomic sampling in future analyses.

Second, differences in jaw and dental structure revealed by this work bear on our understanding of function and feeding ecology in early actinopterygians. In total, we have generated 26 3D models, most of which represent complete or near-complete and largely undistorted mandibular models. Jaws represent a paleobiological model system for addressing questions related to ecological diversification (Gregory, 1933; Wainright & Richards, 1995; Friedman 2009). Beyond the qualitative considerations of function made here, we anticipate that formal quantitative analysis of discrete and continuous traits from this suite of mandibles will help better define ecological diversity in Devonian actinopterygians. This, in turn, provides necessary context for understanding the apparent proliferation of jaw and tooth structures in Carboniferous and younger taxa (Sallan & Friedman, 2012; Sallan, 2014; Friedman et al., 2018).

Our expanded descriptive and 3D data of the mandible of Devonian actinopterygians provides raw character data for establishing the genealogy of the earliest ray-fins as well as functional traits that can be examined in light of that refined systematic framework. Together, these will help illuminate evolutionary patterns deep in the history of one of today’s most successful vertebrate groups.

## AUTHOR CONTRIBUTIONS

Project conception: MF, SG

Funding acquisition: MIC, MF, SEP, SG

Data acquisition: KD, MF, RTF, SG, VF

Data curation: JL, MF, RRH, SEP, SG

Data segmentation: BI, MF, RTF, RRH, SG

Figure preparation: BI, SG

Manuscript writing: BI, SG

Comments and editing: BI, KD, EMT, JL, MIC, MF, MZ, RTF, RRH, SEP, SG, VF

## ACKNOWLEDGEMENTS

We thank the many collections staff who have made this work possible, including Alana Gishlick (AMNH), Annelise Foley (IRSNB), Bent Lindow and Laura Cotton (NHMD), Caitlin Colleary and Jeb Bugos (CMNH), Christina Byrd (MCZ), Emma Bernard (NHMUK), Kacey Page (BSNS), Ned Gilmore and Ted Daeschler (ANSP), Scott Johnston (MCZ), Stig Walsh (NMS), Thomas Mörs (NRM), Tim Ziegler (MV), and Yong Yi Zhen (AM). We thank Brett Clark (Imaging and Analysis Centre, NHMUK) and Liz Martin Silverstone (XTM Facility, Palaeobiology Research Group, University of Bristol) for assistance with CT scanning. SG and BI were supported by a Royal Society Dorothy Hodgkin Research Fellowship (no. DH160098) and the National Environment Research Council (NE/X016633/1) both to SG. SEP and RRH were supported by the National Science Foundation EAR (2219069) to SEP. MF and ET were supported by a grant from the National Science Foundation (EAR 2219007) to MF. MIC was supported by a grant from the National Science Foundation (NSF EAR 2218892). We acknowledge the European Synchrotron Radiation Facility (ESRF) for provision of synchrotron radiation facilities under proposal number ES1299 and we would like to thank the beamline staff for assistance and support in using beamline BM05.

---

**Table 1:** Overview of Devonian actinopterygians.

| **Name** | **Previous literature** | **Included in study** | **Reason for exclusion** | **Stage** | **Locality** | **Stratigraphy** |
| --- | --- | --- | --- | --- | --- | --- |
| *Meemannia eos* | Zhu et al., 2006, 2010; Lu et al., 2016 | Yes | n/a | Lochkovian | Xitun, Qujing, East Yunnan, China | Xitun Formation, Qujing |
| *Cheirolepis trailli* | Pearson & Westoll, 1979; Giles et al., 2015a | Yes | n/a | Eifelian | Tynet Burn; Clune; Lethen Burn; Orkey, Scotland | Achanarras Fish Bed Member, Lybster Flagstone Formation |
| *?Howqualepis youngorum* | Choo, 2009 | No | Lower jaw not preserved | Eifelian | New South Wales, Australia | Bunga Beds |
| *Austelliscus ferox* | Figuerola et al., 2021 | Yes | n/a | Eifelian- Givetian | Parana, Brazil | Ponta Grossa Formation |
| *Donnrosenia schaefferi* | Long et al., 2008 | No | Not amenable to CT scanning (size, density) | Givetian | Southern Victoria Land, Antarctica | Aztec Siltstone |
| *Howqualepis rostridens* | Long, 1998 | Yes | n/a | Givetian | Upper Howqua River, Mount Howitt, Australia | Lower mudstone unit, Avon River Supergroup |
| *Cheirolepis jonesi* | Newman et al., 2021 | Yes | n/a | Givetian | Estheriahaugen North, Spitsbergen, Svalbard | Fiskeløfta Member, Tordalen Formation, Mimerdalen Subgroup |
| *Cheirolepis schultzei* | Arratia & Cloutier, 2004 | No | Lower jaw incomplete | Givetian | Red Hill, Horse Creek Valley, Eureka County, central Nevada, USA | Red Hills Beds |
| *Stegotrachelus finlayi* | Gardiner, 1963; Swartz, 2009 | No | Not amenable to CT scanning (size) | Givetian | Ness of Sound & Dunrossness, Shetland, Scotland | Brindister Flags Formation, Middle Old Red Sandstone |
| *Moythomasia lineata* | Jessen, 1968; Choo, 2015 | No | Not amenable to CT scanning (size, density) | Frasnian | Heiligenstock Quarry, North Rhine-Westphalia, Germany | Oberer Plattenkalk Member, Ahrdorf Formation |
| *Moythomasia nitida* | Jessen, 1968; Choo, 2015 | No | Not amenable to CT scanning (flattened, density) | Givetian- Frasnian | Heiligenstock Quarry, North Rhine-Westphalia, Germany | Oberer Plattenkalk Member, Ahrdorf Formation |
| *Mimipiscis toombsi* | Gardiner, 1984; Choo, 2012 | Yes | n/a | Frasnian | Paddy's Valley, Mt. Pierre Station, Western Australia | Gogo Formation |
| *Mimipiscis bartrami* | Gardiner, 1984; Choo, 2012 | Yes | n/a | Frasnian | Paddy's Valley, Mt. Pierre Station, Western Australia | Gogo Formation |
| *Moythomasia durgaringa* | Gardiner, 1984; Choo, 2015 | Yes | n/a | Frasnian | Paddy's Valley, Mt. Pierre Station, Western Australia | Gogo Formation |
| *Gogosardina coatesi* | Choo et al., 2009a | Yes | n/a | Frasnian | Paddy's Valley, Mt. Pierre Station, Western Australia | Gogo Formation |
| *Pickeringius acanthophorus* | Choo et al., 2019 | No | Lower jaw incomplete | Frasnian | Paddy's Valley, Mt. Pierre Station, Western Australia | Gogo Formation |
| *Moythomasia perforata* | Gross, 1942 | No | Lower jaw not preserved | Frasnian | Kokenhusen (Koknese), Latvia | Snetnaya Group |
| *Cheirolepis canadensis* | Arratia & Cloutier, 1996; Arratia, 2009. | No | Not amenable to CT scanning (size, preservation) | Frasnian | River Ristigouche, Quebec, Canada | Escuminac Formation |
| *Raynerius splendens* | Giles et al., 2015b | Yes | n/a | Frasnian | La Parisienne quarry, Pas-de-Calais, France | Grey Member, Ferques Formation |
| "*Moythomasia devonica*" | Clarke, 1885; Hussakof & Bryant, 1918; Gardiner, 1963 | Yes | n/a | Frasnian | Folsomdale, New York State, USA | Rhinestreet Shale, West Falls Formation |
| *Krasnoyarichthys jesseni* | Prokofiev, 2002 | No | Lower jaw not preserved | Famennian | Preobrazhensky Village, Nazarovo City, Krasnoyarski Krai, Russia | Unknown |
| *Osorioichthys marginis* | Taverne, 1997 | Yes | n/a | Famennian | Philippeville-Mariembourg Le Fraity Highway, Belguim | Famenne Shales |
| *Palaeoneiros clackorum* | Eastman, 1907; Giles et al., 2022 | Yes | n/a | Famennian | Warren, Pennsylvania, USA | Chadakoin Formation |
| "*Kentuckia hlavini*" | Dunkle, 1964; Feldman, 1996 | Yes | n/a | Famennian | Cuyahoga County, Ohio, USA | Cleveland Shale Member, Ohio Shale |
| *Actinopterygii n gen n sp* | Friedman & Blom, 2006 | Yes | n/a | Famennian | Cuyahoga County, Ohio, USA | Cleveland Shale Member, Ohio Shale |
| *Tegeolepis clarki* | Gardiner, 1963; Dunkle & Schaeffer, 1973; Figueroa et al., 2021 | Yes | n/a | Famennian | Cleveland, Ohio, USA | Cleveland Shale Member, Ohio Shale |
| *Limnomis delaneyi* | Daeschler, 2000 | Yes | n/a | Famennian | Red Hill site, nr Hyner, Clinton County, Pennsylvania, USA | Duncannon Member, Catskill Formation |
| "*Gonatodus brainerdi*" | Newberry, 1889 | Yes | n/a | Famennian | Chagrin Falls, Ohio, USA | ?Berea Sandstone, Appalachian Basin |
| *Cuneognathus gardineri* | Friedman & Blom, 2006 | Yes | n/a | Latest Famennian/earliest Tournaisian | Celsius Bjerg, Ymer ÿ, East Greenland | Obrutschew Bjerg Formation |

**Table 2:**
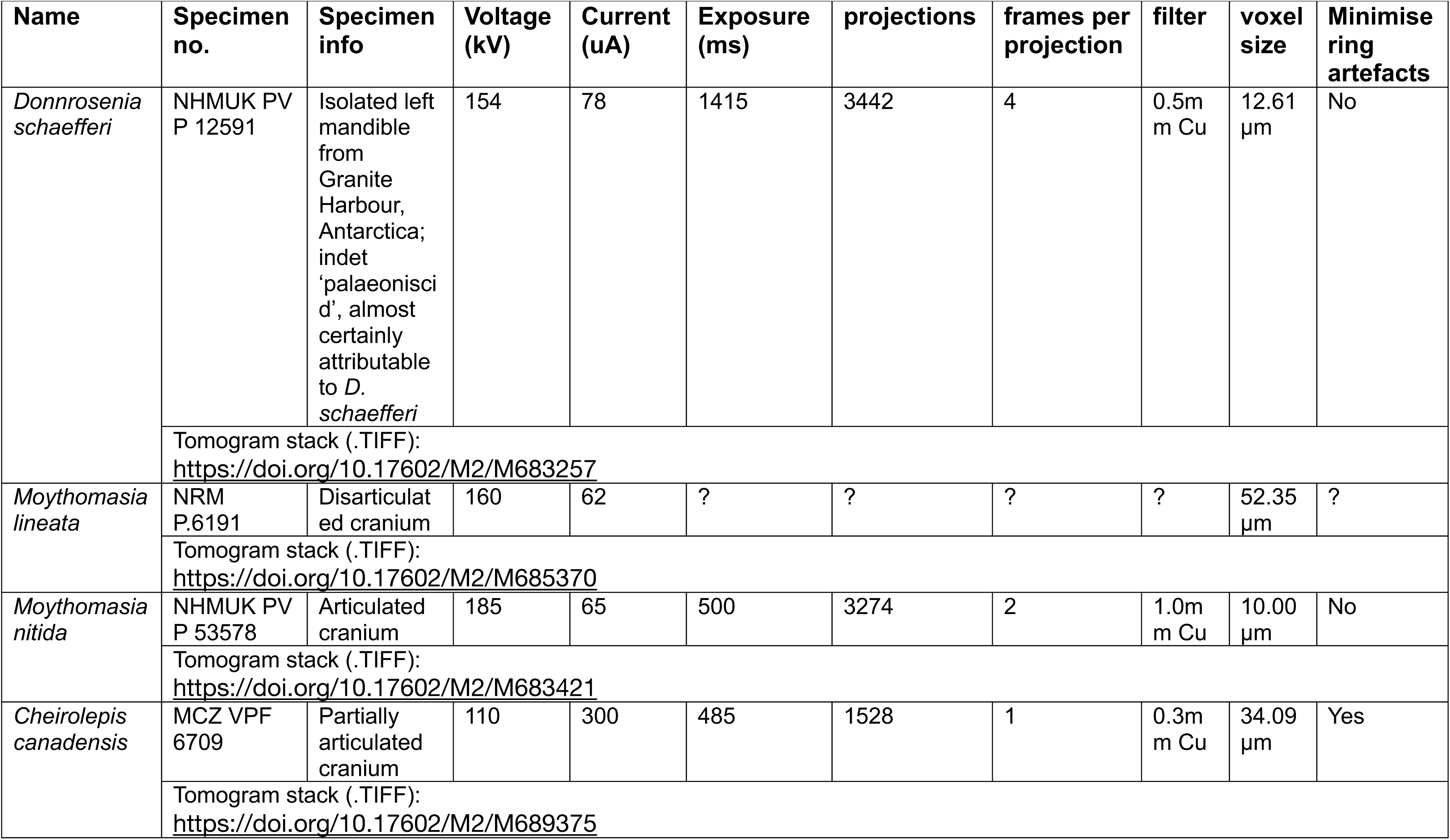
Taxa excluded from the study due to unsuitability of CT scanning.

**Table 3:** Taxa included in study and CT scanning parameters.

| Name | Specimen no. | Specimen info | Mandible | Voltage (kV) | Current (uA) | Exposure (ms) | projections | frames per proj | filter | voxel size | Min. ring artefacts |
| --- | --- | --- | --- | --- | --- | --- | --- | --- | --- | --- | --- |
| *Meemannia eos* | IVPP V14536.5 | Isolated mandible | Left | 110 | 120 | 1000 | 1440 | 2 | 1.00 mm Al | 8.62 μm | No |
| *Cheirolepis trailli* | NHMUK PV P 62908b | Semi-articulated specimen | Right | See materials and methods (synchrotron tomography). |  |  |  |  |  |  |  |
| | NHMUK PV P 1370 | Semi-articulated specimen | Right (and left articular) | 215 | 130 | 1000 | 4476 | 8 | 1.0 mm Sn | 27.33 μm | No |
| *Austelliscus ferox* | MCT890-P | Isolated mandible (mouldic) | Left | Scan data from Figueroa et al. (2021). |  |  |  |  |  |  |  |
| *Howqualepis rostridens* | AMF65495 | Semi-articulated specimen (mouldic) | Right | Scan data from Giles et al. (2015a). |  |  |  |  |  |  |  |
| | MV P.160801 | Poorly-articulated specimen; mouldic | Left (and right anterior partial) | 175 | 314 | 2000 | 2300 | 1 | 0.5 mm Sn | 55 μm | No |
| *Cheirolepis jonesi* | PMO 235.121 | Semi-articulated specimen | Both | Scan data from Newman et al. (2021). |  |  |  |  |  |  |  |
| *Mimipiscis toombsi* | NHMUK PV P 56495 | Isolated mandible (due to acid prep.) | Right | 110 | 50 | 1000 | 4476 | 1 | 0.1 mm Cu | 6.26 μm | No |
| | NHMUK PV P 53249 | Acid prepared cranium | Right | 80 | 238 | 500 | 4476 | 1 | 0.1 mm Cu | 20.26 μm | No |
| *Mimipiscis bartrami* | NHMUK PV P 53254 | Isolated mandible (due to acid prep.) | Right | 79 | 228 | 500 | 4476 | 1 | 0.1 mm Cu | 19.46 μm | no |
| *Moythomasia durgaringa* | AMNH FF 11598 | Articulated specimen (acid prepared) | Right | 150 | 165 | 708 | 3141 | 2 | None | 27.68 μm | no |
| *Gogosardina coatesi* | MV P.228269 | Isolated mandible (due to acid prep.) | Left | 150 | 107 | 1000 | 700 | 2 | 0.5 mm Al | 16 μm | No |
| *Raynerius splendens* | MGL 1245 | Articulated specimen | Both | Scan data from Giles et al. (2015b). |  |  |  |  |  |  |  |
| "*Moythomasia devonica*" | BSNS E22113 | Isolated mandible | Left | 155 | 105 | 1415 | 3141 | 2 | None | 18.13 μm | No |
| *Osorioichthys marginis* | IRSNB P 1340 | Articulated specimen | Right |  |  |  |  |  |  |  |  |
| *Palaeoneiros clackorum* | MCZ VPF-5114 | Articulated specimen | Both | Scan data from Giles et al. (2023). |  |  |  |  |  |  |  |
| "*Kentuckia hlavini*" | CMNH 9562 | Isolated mandible | Left | 145 | 130 | 1000 | 3141 | 1 | 0.1 mm Cu | 23.71 μm | no |
| *Actinopterygii* n. gen. n. sp. | CMNH 9560 | Isolated mandible | Right | 135 | 125 | 1000 | 3141 | 1 | 0.1 mm Cu | 17.86 $\mu$ | No |
| *Tegeolepis clarki* | CMNH 8124 | Isolated mandible and fragments | Right | Scan data from Figueroa et al. (2021). |  |  |  |  |  |  |  |
| | NHMUK PV P 45312 | Semi-articulated specimen | Right | 205 | 225 | 1415 | 4150 | 2 | 2.0 mm Cu | 122.1 μm | Yes |
| *Limnomis delaneyi* | ANSP 23721 | Semi-articulated specimen | Both | 80 | 80 | 4000 | 3141 | 2 | 1.4 mm Al | 6.53 μm | Yes |
| *"Gonatodus"*<br>*brainerdi* | ANSP 6232 | Articulated<br>specimen | Left | 155 | 155 | 2829 | 3141 | 4 | 1.25m<br>m Cu | 24.03<br>μm | Yes |
| *Cuneognathus*<br>*gardineri* | NHMD-1235389 | Semi-articulated<br>specimen | Both | 105 | 110 | 2829 | 3141 | 2 | 0.25<br>mm<br>Cu | 12.01<br>μm | Yes |

**Table 4:**
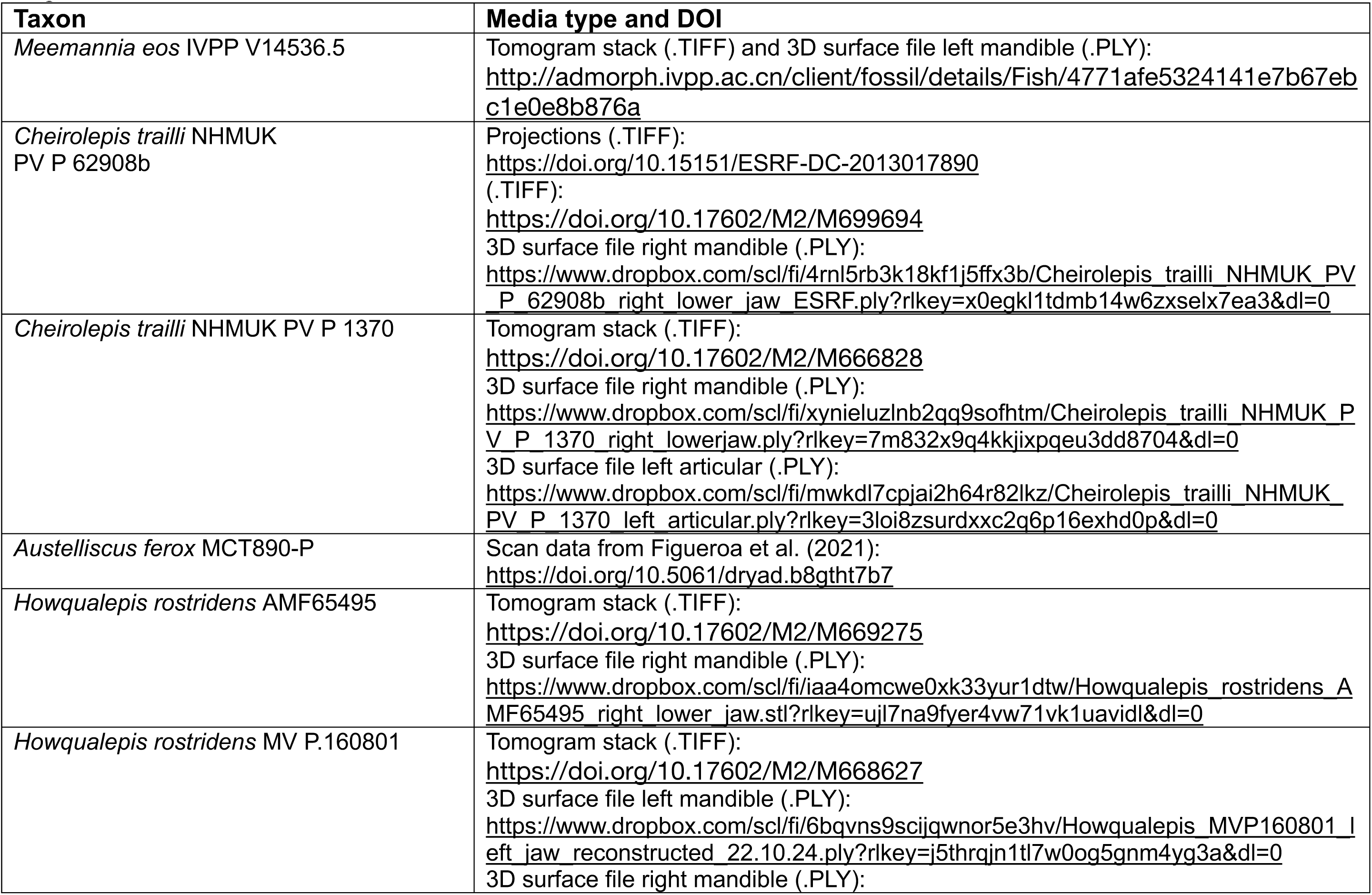
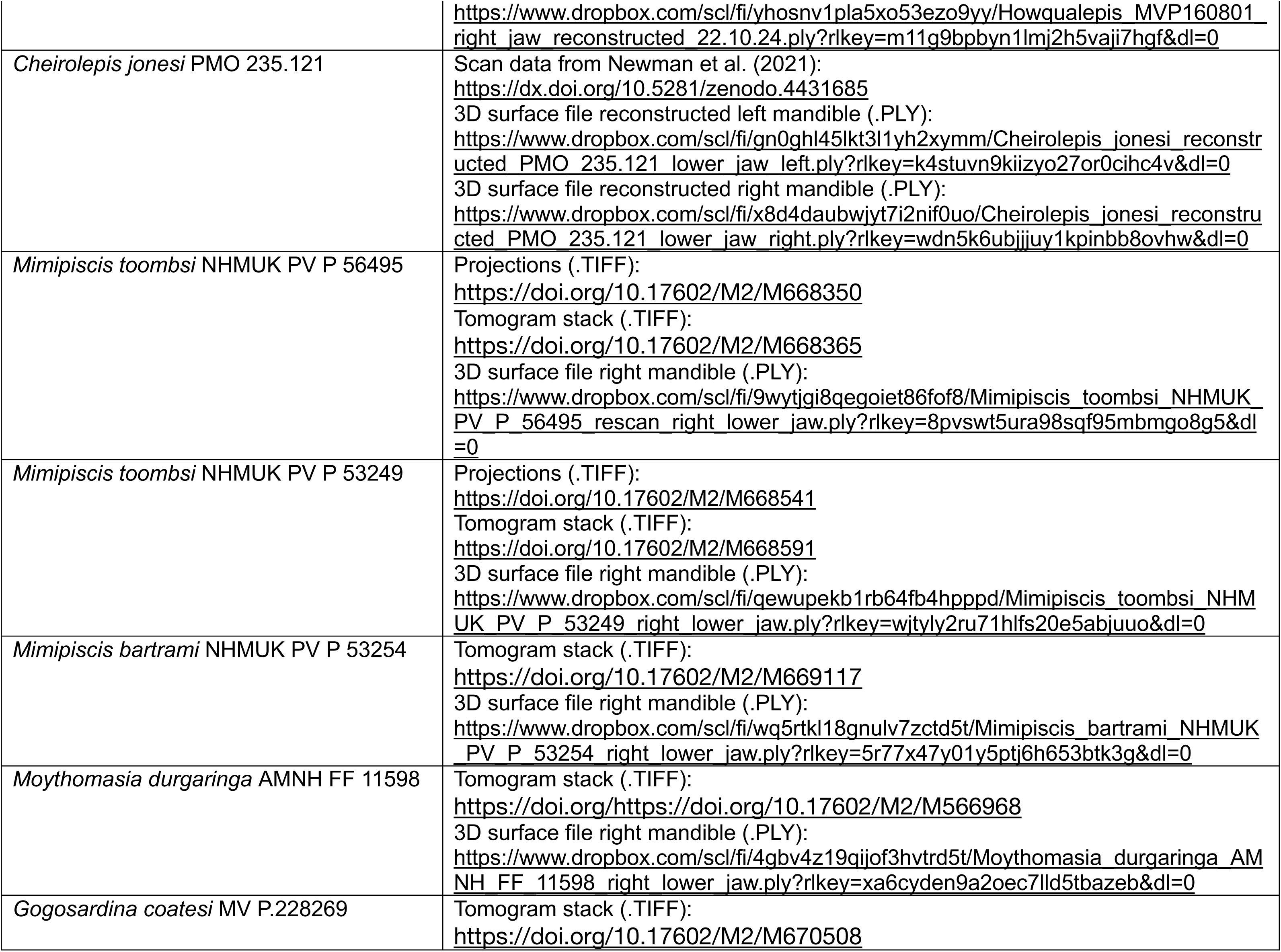
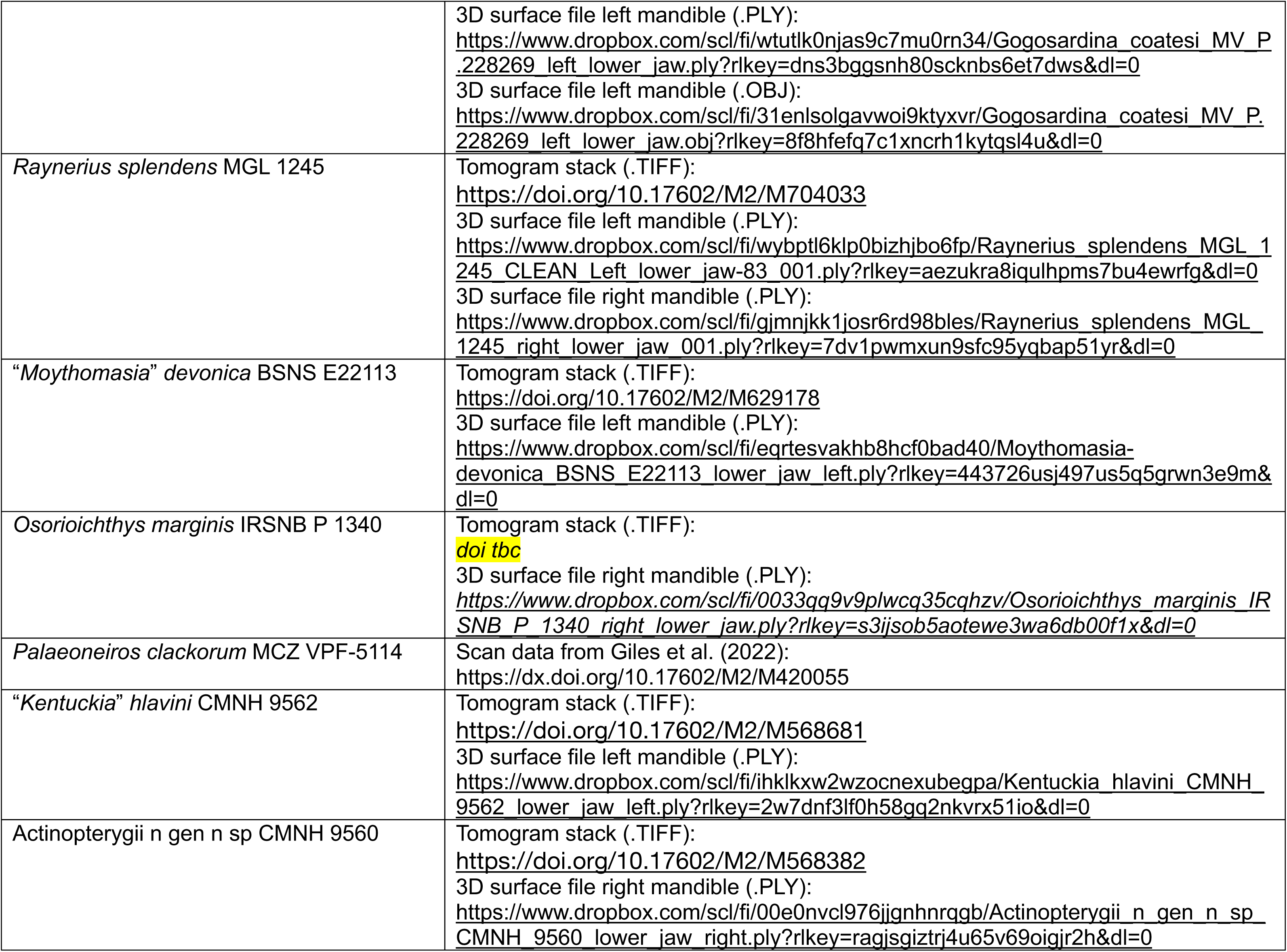
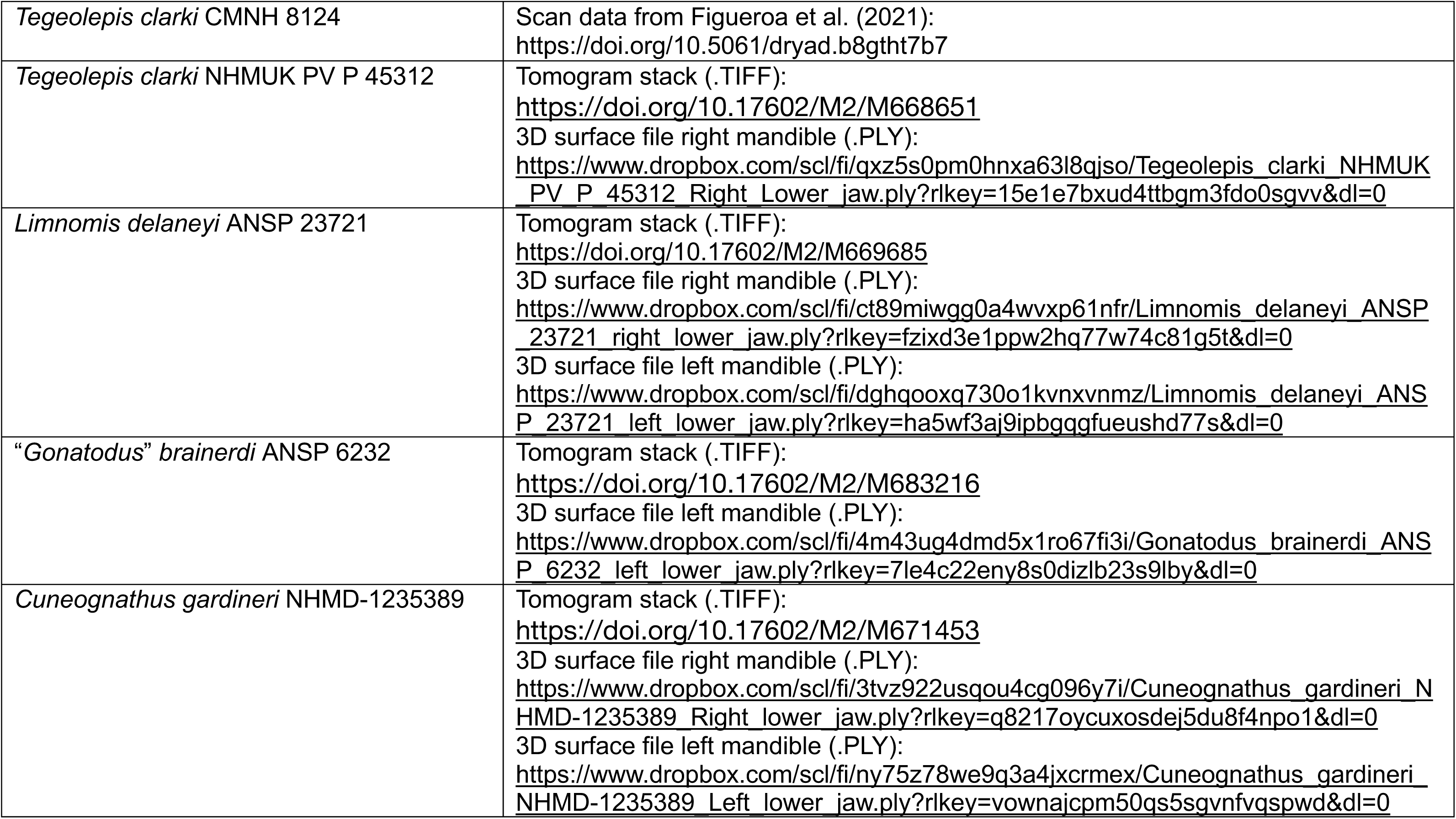
Data availability. Temporary Dropbox links are provided for the PLY files during the review process, and will be replaced with stable Morphosource DOIs upon acceptance.

## Supplementary figures

**Supplementary Figure 1.**
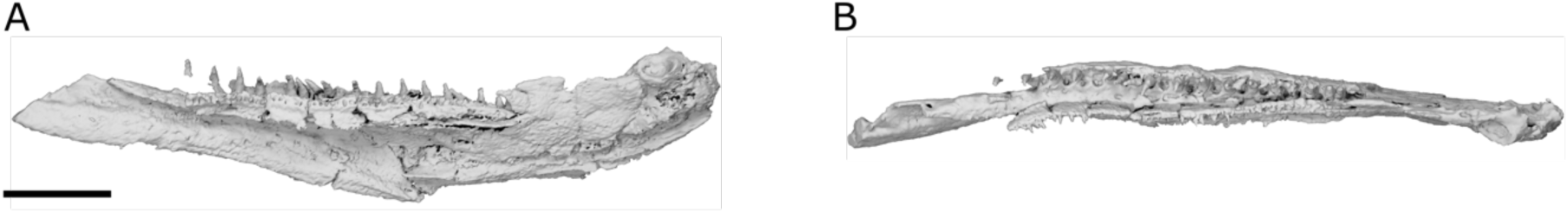
Mandibles of *Cheirolepis trailli* NHMUK PV P 1370 prior to retrodeformation. **a,** right mandible in medial view. **b,** right mandible in dorsal view. Scale bar = 5 mm.

**Supplementary Figure 2.**
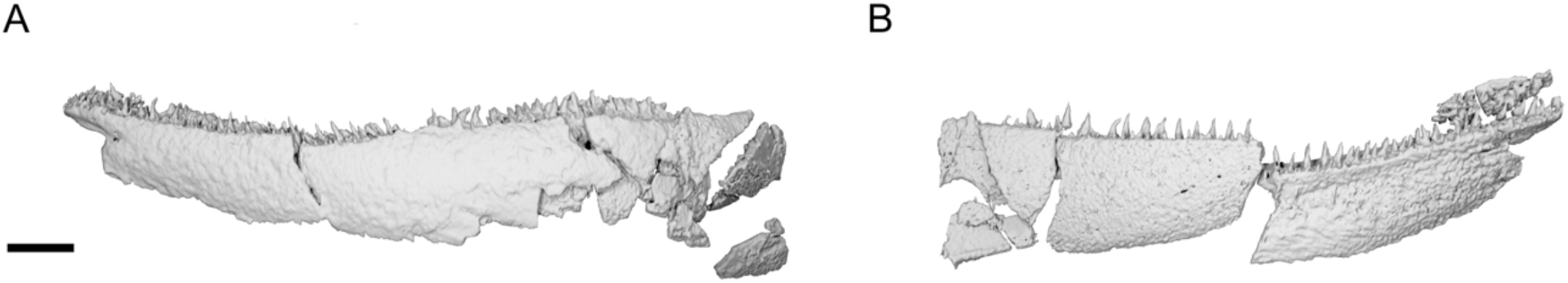
Mandibles of *Cheirolepis jonesi*, PMO 235.121 prior to retrodeformation. **a,** left mandible in lateral view. **b,** right mandible in lateral view. Scale bar = 5 mm.

**Supplementary Figure 3.**
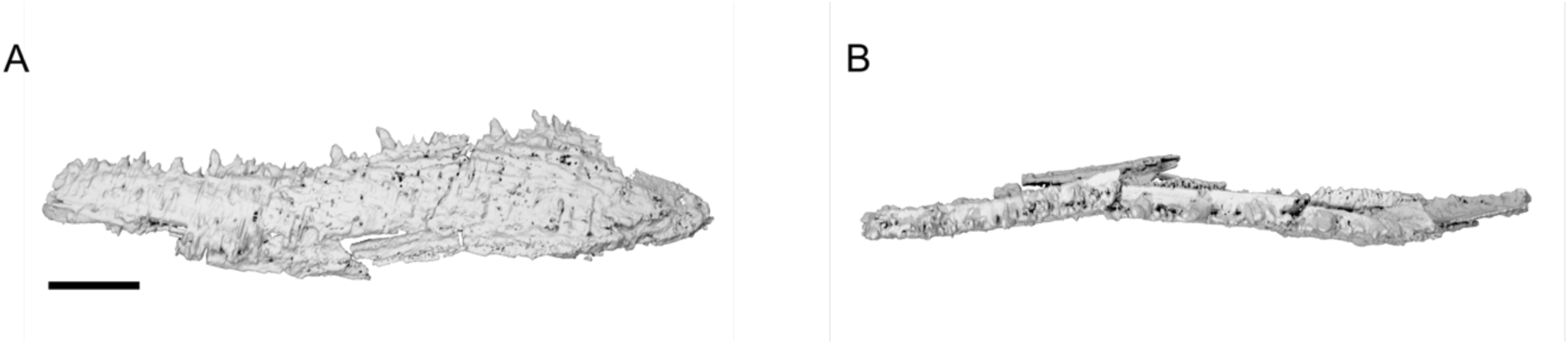
Mandibles of *Limnomis delaneyi* ANSP 23721 prior to retrodeformation. **a,** left mandible in lateral view. **b,** right mandible in lateral view. Scale bar = 1 mm.

